# The structural logic of insect olfactory receptor assembly and gating

**DOI:** 10.64898/2026.08.14.744483

**Authors:** Navid Paknejad, Dragana Nešić, Beth Graczyk, Jackson A. Rogow, Mackenzie A. Yedlin, Chidera Udedibia, Joel A. Butterwick, Vanessa Ruta

## Abstract

Insects detect the chemical world using a large family of odorant-gated ion channels, each formed from a variable odorant-binding subunit (OR) and a single conserved co-receptor Orco. This modular organization is thought to allow tuning ORs to diversify their chemical recognition while Orco provides structural stability to the heteromer. Yet Orco can be autonomously activated by synthetic agonists, suggesting that it may contribute to channel gating rather than serving solely as a structural scaffold. Here, we combine cryo-electron microscopy with analyses of receptor stoichiometry and function to define the structural logic underlying Orco-OR assembly and gating. We show that Orco retains the canonical ligand-binding pocket of ORs but is chemically insulated from environmental odorants by a phospholipid that occupies this site. The Orco agonist VUAA4 instead binds a membrane-accessible crevice adjacent to the gate, defining a distinct site of allosteric modulation. We further demonstrate that Orco-OR heteromers can assemble in multiple stoichiometries through shape complementarity within the intracellular anchor domain and resolve structures with both a 3:1 and 2:2 architecture. Receptors constrained to a 2:2 stoichiometry are functional but productive gating requires cooperative engagement of multiple subunits within the heteromer. Comparison with the distinct gating states of a basal homomeric olfactory receptor suggests that existing Orco-OR structures capture nonconductive intermediates within the broader conformational landscape of this receptor family. Together, these findings suggest how Orco can flexibly assemble and function with highly divergent ORs, acting not simply as a structural scaffold but as an integral partner in cooperative channel gating, thereby enabling the extraordinary diversification of insect olfactory receptors.

## Introduction

Olfactory systems must detect and discriminate among an immense diversity of volatile chemicals. Many species, from insects to mammals, have converged on a common solution to contend with the high-dimensional chemical world, relying on the orchestrated activity of large families of olfactory receptors to combinatorially encode odor identity^1–3^. Central to this coding strategy is that individual receptors must be tuned to different regions of chemical space, placing strong constraints on olfactory receptor architecture and evolution^4^.

Insects use a distinct family of olfactory receptors for chemical detection. Rather than signaling through second messenger cascades as in vertebrate receptors^5,6^, they form heteromeric odorant-gated ion channels^7,8^ comprised of two types of subunits: a highly conserved co-receptor, Orco^9,10^ and a highly divergent odorant receptor (OR) that harbors the ligand binding site and confers chemical specificity to the complex. Each species expresses a unique repertoire of tens to hundreds of divergent ORs, whose chemical tuning has evolved to meet the distinct demands of its ecological niche. By contrast, nearly all insects possess just a single Orco gene that is highly conserved across distant clades, reflecting an essential role in olfactory transduction. While ORs from basal insects can function autonomously^11,12^, accompanying the dramatic radiation of neopteran species, ORs became dependent on Orco to assemble, traffic, and function^7,10,13,14^. Orco thus serves as an essential component of insect olfactory receptors, where it has been proposed to confer structural stability to the heteromeric complex, thereby relaxing evolutionary constraints on the ORs and facilitating the diversification of millions of receptors with distinct chemical tuning^10,15^.

Recent structural studies have begun to shed light on the organization of Orco-OR heteromers through snapshots of receptor architecture captured by single particle cryo-electron microscopy^16,17^. To date, all resolved heteromer structures display a 3:1 assembly, in which three Orco subunits and a single OR encircle the central ion conduction pathway and contribute to the hydrophobic gate that constricts its extracellular entryway. While the OR undergoes conformational rearrangements upon odorant binding, the three Orco subunits remain largely stationary, leading to the proposal that Orco serves primarily as a rigid scaffold to support odorant signaling through the OR^16,17^. This asymmetric gating mechanism contrasts with the symmetric gating observed in homomeric ORs from basal insects^12^ and gustatory receptors^18–20^, in which all four subunits work in concert to dilate the pore. Whether existing structures of the Orco-OR heteromer capture the full range of functional receptor assemblies—or only a subset of the conformational states sampled during gating—remains unclear.

The modular architecture of Orco-OR heteromers poses a fundamental question: how can a single conserved co-receptor support the function of a large and evolving repertoire of ORs while preserving their distinct chemical tuning? Two features of Orco are essential to this role. First, Orco must remain chemically inert to preserve the fidelity of OR-dependent signaling. Although Orco is insensitive to odorants, it can function as an autonomous channel activated by synthetic agonists like VUAA1 and VUAA4^21,22^, implying an intrinsic capacity to gate not captured by existing structures. Second, a single Orco must be able to assemble and function with tens to hundreds of highly divergent ORs^23,24^ that share little (<20%) sequence conservation. Heteromer assembly must therefore rely on flexible modes of interaction to enable Orco to act as a universal co-receptor for this receptor family.

Here, we combine structural and functional analyses to show that Orco-OR complexes can assemble in multiple configurations, including both the previously reported 3:1 Orco-OR stoichiometry as well as a novel 2:2 arrangement. We identify a conserved Trp-His-Tyr (WHY) structural motif that anchors Orco-OR interactions, allowing highly divergent receptors to assemble through shape complementarity. Despite such flexible assembly, we find that productive odorant signaling requires concerted gating rearrangements from multiple subunits within the tetramer. We further show that Orco is insulated from environmental odorants by a phospholipid that occupies its canonical ligand-binding pocket, whereas the synthetic agonist VUAA4 binds at a distinct membrane-accessible site beneath the gate. Unexpectedly, however, Orco remains stationary, even when bound to VUAA4 and assembled with multiple odorant-bound ORs, suggesting that existing Orco-OR structures represent snapshots of nonconductive states. By examining the conformational states of a homomeric OR from a basal insect, we show that the 2:2 heteromer resembles a lower-symmetry intermediate gating state, suggesting that key features of cooperative gating appear to be shared across ancestral homomeric receptors and derived Orco-OR complexes. Together, these results suggest that Orco is not merely a static scaffold for assembly, but an integral participant in a conserved cooperative gating mechanism.

### Selective odorant recognition by CpOR9

While ORs and Orco share a conserved architecture^10,12,15,25^, they fulfill distinct roles in the heteromeric complex: ORs selectively recognize odorants, whereas Orco must remain unresponsive to them to preserve the fidelity of OR-driven signaling. Yet, Orco retains an intrinsic capacity to gate, as evidenced by its autonomous activation by synthetic agonists such as VUAA4^22^. To determine how Orco can be activated by VUAA4 while remaining insensitive to odorants, we elucidated the structure of an Orco-OR heteromer bound to both types of ligands simultaneously, allowing for direct comparison of their binding modes. For the tuning receptor, we chose OR9 from the mosquito *Culex pipiens* genome (*Cp*OR9), an unusually conserved OR selective for indole and 3-methyl indole (skatole)—microbially derived volatiles with broad ecological salience across species^26,27^. The striking evolutionary conservation^28,29^ and narrow tuning of the *Cp*OR9 clade^27^ suggest that these receptors occupy a constrained region of sequence space, providing a useful structural counterpoint for distinguishing odorant recognition by ORs from allosteric activation of Orco. Leveraging the ability of Orcos to assemble and function with evolutionarily distant ORs from other species^30,31^, we co-expressed and purified *Cp*OR9 with the Orco from the fig wasp, *Apocrypta bakeri* (*Ab*Orco)^15^. Both subunits carried the same high-affinity nanobody tag and were purified in the presence of saturating concentrations of indole and VUAA4 for single-particle cryo-EM. The most abundant class of particles was composed of one OR and three Orco subunits, as observed in other heteromeric structures^16,17^, which we resolved to 2.2 Å, allowing unambiguous modeling of the entire assembly (**Fig. 1a; Extended Data Fig. 1**).

**Fig. 1.**
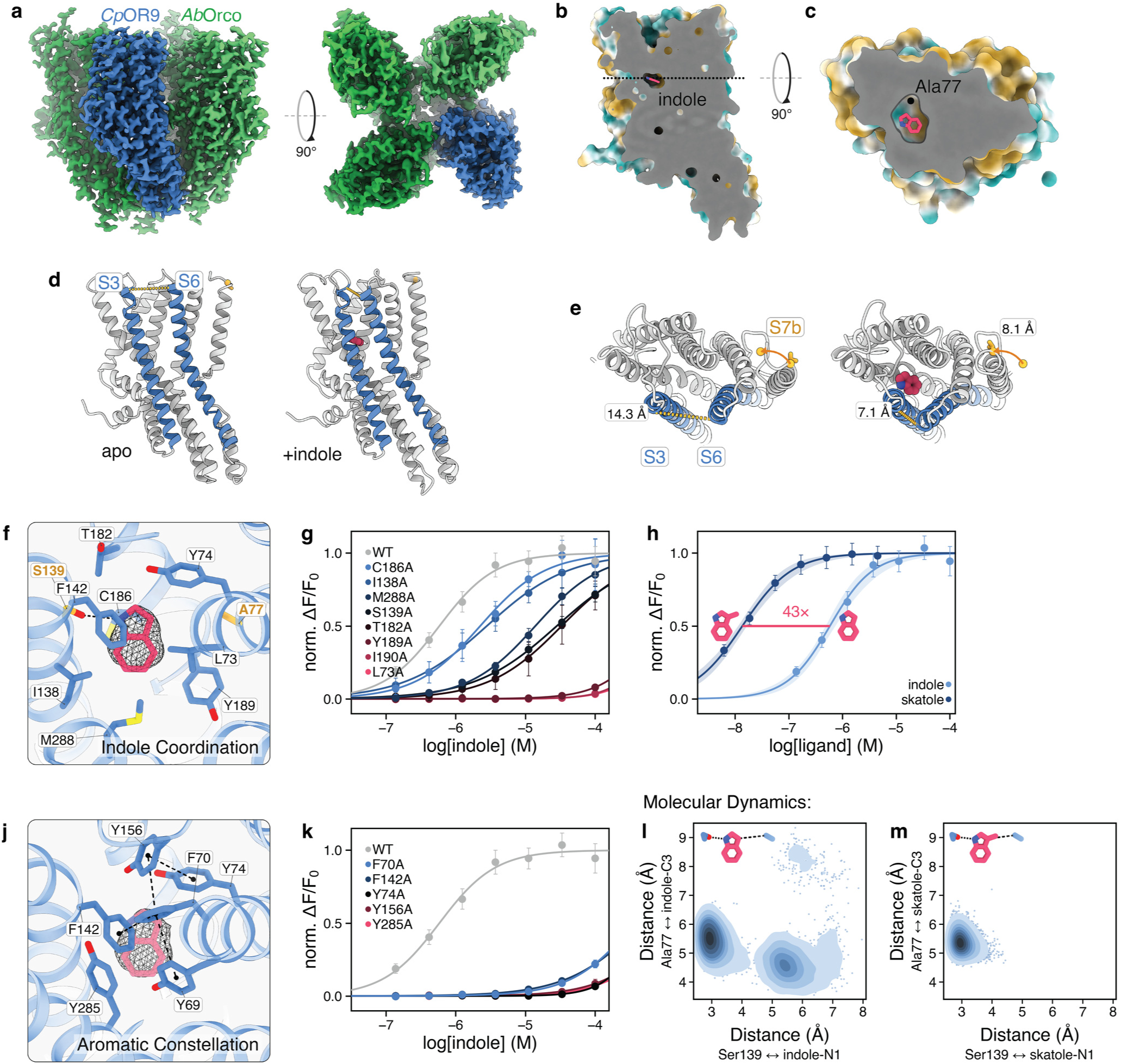
Structural basis of indole recognition and gating by the *Ab*Orco–*Cp*OR9 heteromer. **a**, Cryo-EM density of the 3:1 *Ab*Orco–*Cp*OR9 heteromer viewed from within the membrane plane (side view; left) and extracellular space (top view; right) with Orco colored in shades of green and *Cp*OR9 colored blue. **b**, **c**, Cutaway surface representation of the *Cp*OR9 binding pocket colored by hydrophobicity (teal is hydrophilic; gold is hydrophobic), showing with bound indole shown as side (**b**) and top (**c**) views. In (c) the cavity adjacent to the indole C3 position formed by Ala77 is highlighted. **d**, Side views of the *Cp*OR9 in the ligand-free (apo; left) and indole-bound (right) states with S3 and S6 highlighted in blue. **e**, Corresponding top views of the S7b gate in the apo and indole-bound states. Distances between top of S3 and S6 and across the hydrophobic gate at residue Val374 on S7b are indicated illustrating the conformational changes associated with indole binding. **f**, Coordination of indole within the *Cp*OR9 binding pocket. Cryo-EM density corresponding to indole is shown (mesh), with residues within 4 Å of ligand are shown as sticks. **g**, Dose– response curves for alanine substitutions of residues in the indole coordination shell, listed in order of EC_50_. Hill fits are individually scaled from 0 to 1. Error bars indicate 95% confidence intervals (CIs) of the mean. n=12-52. **h**, Dose–response curves comparing activation of *Ab*Orco-*Cp*OR9 by indole versus skatole, with thumbnails showing the methyl-substitution position on skatole (3-methyl indole). Skatole elicits athe 43-fold lower EC_50_. Each Hill model fits are individually scaled between 0 and 1. Error bars represent 95% CI of the mean scaled by the normalization factor; shaded bands represent the 95% CI envelope of Hill model fits using non-parametric bootstrap resampling of well-level data. n=40-52. **j**, Aromatic residues surrounding the indole-binding site that form a π-interaction network. Predicted interactions (calculated using PLIP) are shown as dashed lines. **k**, Dose-response curves for alanine substitutions of the aromatic residues shown in (**j**) listed in order of EC_50_. n=12-40. **l**, **m**, Molecular dynamics analysis of ligand positioning within the *Cp*OR9 pocket. Kernel density estimate show the joint distributions of the Ser139-to-ligand N1 distance and the Ala77-to-ligand C3 distances for indole (**l**) and skatole (**m**), each aggregated across 6 independent 100 ns simulations. Molecular thumbnails indicate the corresponding measured distances.

Density corresponding to indole was observed within the canonical ligand-binding pocket shared across insect ORs, Orcos, and gustatory receptors (GRs), formed from the splayed arrangement of the S2, S3, S4, and S6 helices (**Fig. 1b, c; Extended Data Fig. 2a, b**). Comparison of the bound and apo state of *Cp*OR9 (**Extended Data Fig. 3**) revealed that indole binding induces a concerted reorganization o**f residues surrounding the pocket, drawing the S3** and S6 helices ∼7 Å closer through their interactions with the ligand (**Fig. 1d, e**). This conformational change propagates through S5 to the central ion-conduction pathway, inducing the pore-lining S7b helix to tilt away from the pore’s central axis—a gating rearrangement broadly conserved among divergent ORs^12,16,17^ (**Extended Data Fig. 3a–c**).

The high resolution of our structure allowed for unequivocal placement of indole within the pocket, shedding light on the highly selective tuning of this receptor. The asymmetric density enabled us to distinguish between indole’s five- and six-membered rings, with its pyrrole nitrogen positioned in hydrogen-bonding distance of Ser139, while its benzene ring extends along a hydrophobic cleft at the base of the pocket (**Fig. 1f**). Consistent with this pose, mutation of Ser139 to alanine reduced the apparent affinity for indole 48-fold, supporting a role for its hydroxyl group in hydrogen bonding to the ligand and orienting it within the pocket (**Fig. 1g**).

Across mosquito species, receptors within the same clade as *Cp*OR9 preferentially detect either indole or skatole (3-methyl indole) with high affinity despite sharing significant sequence conservation^27,32^. Our analyses suggest a structural basis for this selectivity. When bound to *Cp*OR9, the 3-carbon of the indole pyrrole ring points towards a small cavity lined by Ala77 (**Fig. 1c**). Molecular dynamics simulations indicate that while indole remains primarily anchored via its hydrogen bond with Ser139, it can transiently sample this cavity (**Fig. 1l; Extended Data Fig. 4a, c**). By contrast, the methyl at the 3-carbon position in skatole projects into this pocket, constraining the ligand and stabilizing its pose (**Fig. 1m; Extended Data Fig. 4b, d**). Consistent with this model, *Cp*OR9 displayed >40-fold higher apparent affinity for skatole than indole (**Fig. 1h**), whereas related indolergic receptors bearing a bulkier leucine at the equivalent Ala77 position show a strong shift in preferential sensitivity towards indole^33^. Thus, small changes in the volume of the odorant-binding pocket appear sufficient to discriminate between closely related microbial metabolites that convey distinct ecological information.

Beyond this key determinant of ligand specificity, the apo and bound structures of *Cp*OR9 reveal how odorant recognition is organized into two functionally distinct layers that together couple odorant binding to the helical rearrangements underlying gating of the pore. The inner layer is formed by eleven residues within direct contact of indole (<4 Å), comprising a distribution of polar, aromatic, and aliphatic side chains (**Fig. 1f**). Substituting any of these residues with alanine decreased the receptor’s apparent affinity for indole, consistent with a distributed network that stabilizes the ligand within the pocket (**Fig. 1g**). Above this direct coordination shell, six broadly conserved aromatic residues—Tyr69, Phe70, Tyr74, Phe142, Tyr156, and Tyr285—separate the buried binding pocket from the extracellular solution and rearrange upon indole binding to form an extensive π-interaction network (**Fig. 1j**). Although these aromatic residues are not in direct contact with the ligand, mutation of any of them to alanine strongly attenuated indole signaling (**Fig. 1k**), suggesting that perturbing this aromatic layer disrupts the local rearrangements required to couple indole binding to channel opening, thereby destabilizing the open state. The single predominant binding pose observed in *Cp*OR9 indicates that narrowly-tuned indolergic ORs achieve high-affinity discrimination through precise ligand coordination within a layered pocket architecture, a striking contrast to the more permissive pockets of broadly tuned receptors such as *Mh*OR5, which accommodate multiple degenerate ligand poses^12^. Specialized features of the *Cp*OR9 binding pocket therefore contribute to its selective recognition of odorants, providing a direct point of comparison to consider how the homologous pocket in Orco has been adapted to render it odorant-insensitive.

### A phospholipid occupies the binding pocket of Orco

Orco retains an extracellular cavity similar to the canonical ligand-binding pocket of ORs and GRs, raising the possibility that it serves as a conserved site for modulation by synthetic agonists (**Fig. 2a–c**). Indeed, VUAA4 was previously proposed to bind within this cavity^15,34^. Unexpectedly, rather than harboring VUAA4, the *Ab*Orco pocket is occupied by a phospholipid, whose polar headgroup and glycerol backbone reside in the analogous position occupied by volatile odorants in ORs (**Fig. 2c, d; Extended Data Fig. 5a, b**). The lipid’s acyl tails snake toward the membrane through a hydrophobic cleft between the S3 and S6 helices that is lined with conserved residues bearing small side chains—including Ser142, Ser145, Ala380, and Gly384 (**Fig. 2d**). By contrast, bulkier hydrophobic residues at these positions in ORs bridge across the S3-S6 interface, rendering this gap too narrow to allow lipid penetration. Lipid occupancy may therefore occlude the canonical ligand-binding site in Orco, preventing spurious activation by odorants.

**Fig. 2.**
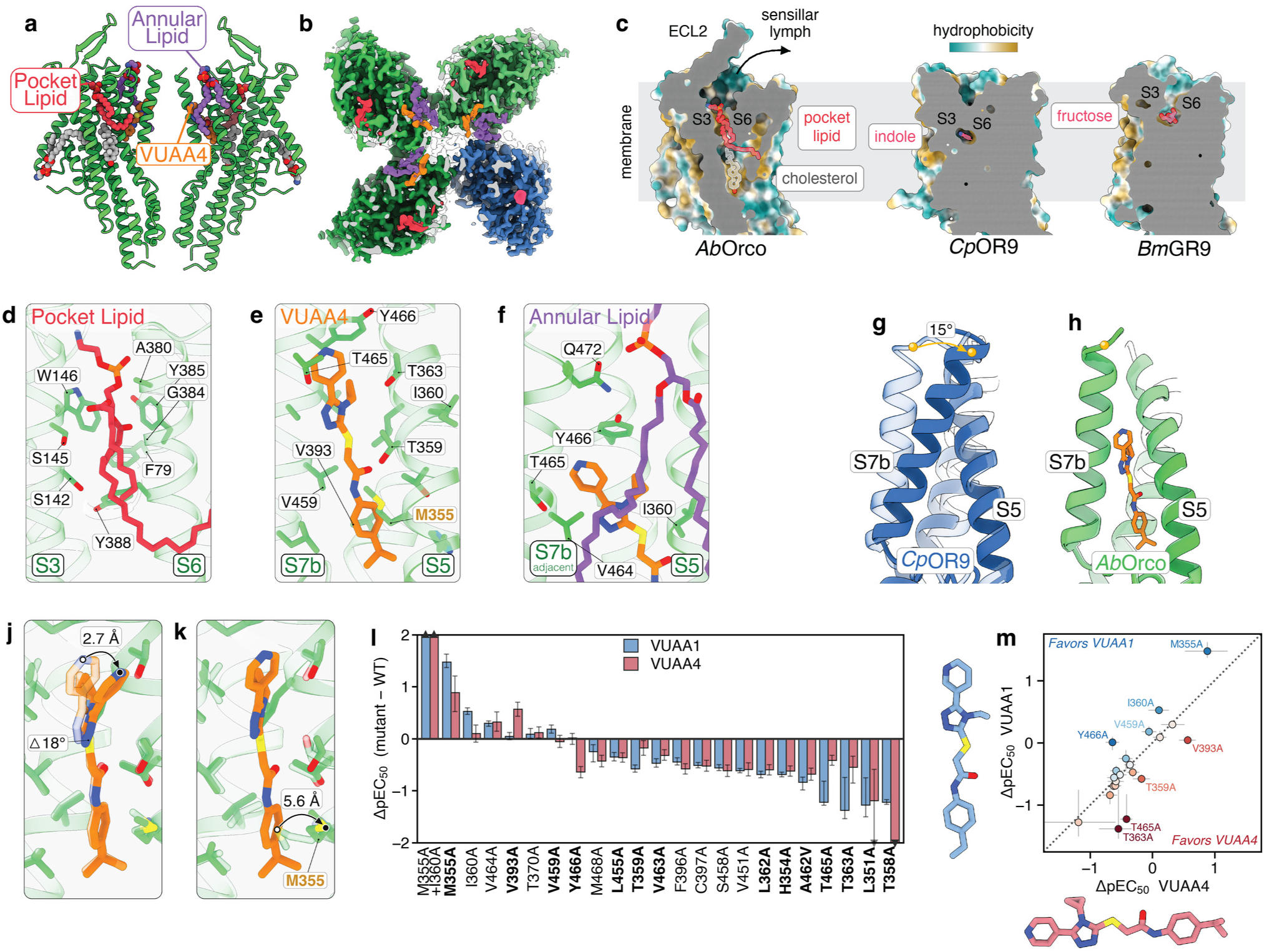
A membrane lipid occupies the Orco binding pocket, while VUAA4 binds to a distinct crevice near the pore. **a**, Overview of the positions of bound phospholipids (red, purple, and gray), cholesterol (gray), and VUAA4 (orange) in models of the two opposing *Ab*Orco subunits from the 3:1 heteromeric structure. **b**, Extracellular cutaway view of the cryo-EM density from the 3:1 heteromer showing bound agonists (indole, magenta; VUAA4, orange), pocket lipid (red), and the annular lipid at the pore (purple). *Cp*OR9 density is blue, the three *Ab*Orco subunits green. **c**, Hydrophobicity colored cutaway representation of the binding pockets of *Ab*Orco (pocket lipid, red; cholesterol, gray), *Cp*OR9 (indole, magenta), and the gustatory receptor *Bm*GR9 (fructose, magenta; PDB: 8UVU), sliced through the pocket with the extracellular space oriented upward and intracellular anchor domain cropped. **d**, Zoomed-in view of the *Ab*Orco pocket lipid with relevant residues and helices labeled. **e**, Zoomed-in view of the VUAA4 binding crevice with relevant residues and helices labeled. **f**, Zoomed-in view of the annular lipid at the pore with relevant residues and helices labeled. **g**, **h**, Comparison of the relative positions of S7b position to S5 from apo (transparent) and indole-bound (opaque) state of *Cp*OR9 (**g**), and the corresponding view for apo (transparent) and VUAA4-bound (opaque) state of *Ab*Orco (**h**) in the 3:1 assembly, highlighting the location of the VUAA4 binding site relative to the helical rearrangements in *Cp*OR9 during gating. **j**, Comparison of the VUAA4 conformation when bound to an *AbOrco* subunit adjacent to another *Ab*Orco subunit (transparent) or adjacent to the *Cp*OR9 (opaque). VUAA4 bends 18° about its sulfur atom leading to the indicated displacement of the pyridyl nitrogen at the end of the agonist. **k**, Comparison of the Met355 side-chain rotamer in the apo (transparent) and VUAA4-bound (opaque) *Ab*Orco, with the measured displacement of the Met355 sulfur atom indicated.. **l**, Changes in agonist potency for homomeric *Ab*Orco due to alanine substitution of residues that directly contact VUAA4 (<4 Å; bold) or nearby in the experimental structure. *n*=8-100. Effects are shown for both the lower- potency agonist VUAA1 (WT EC_50_ = 41 µM; blue) and higher-potency agonist VUAA4 (WT EC_50_ = 2.5 µM; pink) as ΔpEC₅₀ = pEC₅₀(mutant) − pEC₅₀(WT), where pEC₅₀ = −log10(EC₅₀). WT is the zero line, negative values indicate reduced potency. Bars show the bootstrap-median ΔpEC₅₀; error bars are the asymmetric 95% confidence interval of that difference (see Methods). **m**, Scatter plot of data from (**l**) comparing relative effects of each alanine mutation on VUAA1 and VUAA4 potency as measure by ΔpEC₅₀ relative to WT. The iso-potency diagonal separates mutants that differentially impact one ligand over another. Molecular thumbnails of both ligands are shown on their corresponding axes to highlight chemical modifications to the scaffold.

While Orco’s second extracellular loop (ECL2) was previously proposed to block odorant access by capping the binding pocket^15,35^, we find instead that it forms a structured β-hairpin that projects into the extracellular space, leaving the cavity open to the solvent. Likewise, highly conserved aromatic residues lining Orco’s pocket adopt rotamers that maintain continuity with the extracellular solution (**Extended Data Fig. 5a**) in contrast to ORs where bound odorants are fully encased by hydrophobic residues and sequestered from the extracellular solution (**Fig. 2c; Extended Data Fig. 6a, b**). These structural features create an amphipathic cavity in Orco that is ideally suited to stabilize a membrane phospholipid with its polar headgroup remaining solvent exposed while its acyl tails are accommodated within a narrower hydrophobic cleft (**Fig. 2b**). Lipid density is observed in structures of Orco from distantly related species^16^, demonstrating that this pocket architecture is conserved across the co-receptor family. This structural conservation in Orco stands in stark contrast to tuning ORs, where high sequence and structural variability within the canonical pocket underlies their diverse chemical tuning. The evolutionary conservation of this architecture across Orcos indicates that the lipid blockade represents a broadly conserved mechanism for preserving Orco’s insensitivity to environmental odorants.

### VUAA4 binds in a crevice near the pore

With the canonical binding pocket of Orco occluded by a lipid, we reasoned that allosteric modulation by synthetic molecules like VUAA4 must arise from a different site. Indeed, unambiguous density for VUAA4 was tucked into a distinct elongated crevice formed between S5 and S7b near the ion permeation pathway just below the gate (**Fig. 2a, b, e**). To access this site, VUAA4 would first need to partition into the membrane—a route consistent with the compound’s high hydrophobicity (LogP of 4.5).

This membrane-accessible binding mode highlights the unusual architecture of insect chemoreceptors, in which their gating machinery is directly embedded within the lipid bilayer, rather than shielded by peripheral domains as in most other tetrameric ion channels^15^. The pinwheel arrangement of the tetramer creates deep membrane inlets between adjacent subunits that are poised to corral and kinetically trap phospholipids around the pore. Consistent with this organization, well-defined densities for multiple lipids surround the heteromer (**Fig. 2a**), including an annular phospholipid positioned near the pore and directly above VUAA4, cradling the agonist in its binding site (**Fig. 2f**). Molecular dynamics simulations indicate that lipids within these inlets were substantially less mobile than those at the membrane periphery (**Extended Data Fig. 7**), prolonging their residence time near the channel’s gate and the gap between S3 and S6 through which lipids access Orco’s binding pocket. This close spatial association between annular lipids, the pore gate, and VUAA4 suggests an active role for the surrounding bilayer in channel gating. Supporting this hypothesis, mutation of membrane-facing residues adjacent to the annular lipid, such as I360A or V464A, significantly potentiated channel responses to both VUAA1 and VUAA4 (**Fig. 2l**).

VUAA4 adopts an extended conformation spanning 19 Å in its binding crevice, where it is stabilized by van der Waals interactions, forming no hydrogen bonds or π-interactions with Orco residues (**Fig. 2e**). VUAA4 is anchored at one end by its 4-isopropyl aniline ring, which displaces the side chain of Met355 to fit snugly at the base of S5 and S7b (**Fig. 2k**). At the other end, the 4-pyridyl ring contacts Tyr466–a conserved component of the S7b signature sequence (TYhhhhhQF, where h is any hydrophobic amino acid)^36^ that participates in a hydrogen-bond network to stabilize the closed state of the pore across this receptor superfamily (**Extended Data Fig. 2d**). Despite VUAA4 occupancy at all three Orco binding sites, the Orco subunits remain essentially unchanged from the apo state (<1 Å RMSD, **Extended Data Fig. 8**), apart from the localized rotamer shift of Met355. Thus, VUAA4 binding does not stabilize an open state of the pore, suggesting that Orco–OR structures resolved under these conditions may preferentially capture nonconductive states.

Although Orco remains conformationally invariant, the position of VUAA4 reveals why this membrane-facing crevice may represent a critical site for allosteric control. VUAA4 wedges between S5 and S7b, helices whose interface significantly reorganizes during gating in both ORs and GRs (**Fig. 2g, h; Extended Data Fig. 2c**). In *Cp*OR9, for instance, odorant binding draws S5 away from the central pore axis while S7b tilts by ∼15°, forming new contacts one helical turn below those observed in the ligand-free state (**Fig. 2g**). In Orco, VUAA4 sits precisely at the hinge point for these helical movements, ideally positioning the agonist to stabilize analogous gating transitions. Consistent with this model, VUAA4 adopts distinct conformations in its three binding sites, pivoting around its central thioether linkage to accommodate adjacent subunit states—bending by ∼18° when abutting an activated *Cp*OR9 (**Fig. 2j**). The requirement of a flexible central linkage in VUAA4-like agonists^22^ further suggests conformational flexibility is essential for agonist efficacy.

To test the functional importance of this binding crevice, we systematically mutated residues lining the VUAA4 site. Although alanine substitutions of many of the 14 pocket-lining residues reduced VUAA4 efficacy (**Fig. 2l**), the M355A mutation significantly enhanced potency for VUAA4, consistent with its bulky side chain acting as a steric barrier regulating access to the crevice. Combining M355A with the lipid-facing I360A mutation resulted in a ∼260-fold potentiation of Orco activation by VUAA4, emphasizing that ligand access and membrane interactions jointly tune Orco activation.

Comparing the mutational effects on VUAA4 sensitivity with its lower affinity analog, VUAA1^21^, revealed that most pocket mutations affected both agonists similarly, reflecting their common chemical scaffold (**Fig. 2m**). However, four mutations–M355A, T363A, T465A, and Y466A–displayed pronounced differential effects on the EC_50_ values of these two compounds. Mapped onto our structure, these residues cluster directly around the specific chemical substituents that distinguish VUAA1 from VUAA4 (**Fig. 2e; Extended Data Fig. 9d**), validating the orientation and binding pose of the ligand within the density. Because VUAA4 fits tightly within this narrow crevice, subtle variations in ligand geometry would alter steric packing against pocket-lining side chains. These tight geometric constraints explain the stringent structure–activity relationship governing Orco allosteric modulation^22,37–39^, demonstrating how minor alterations to the ligand scaffold produce substantial shifts in agonist potency.

Together, these findings suggest that the spatial separation of the two distinct ligand-binding sites— odorants in the canonical ligand-binding pocket of tuning ORs and synthetic agonists in the membrane-accessible crevice beneath the gate of Orco—provides a structural basis for allosteric communication across subunits within the heteromer. Consistent with functional coupling between these sites, VUAA4 significantly potentiates odorant-evoked signaling, whereas VUAA-derived synthetic antagonists inhibit activation^38–40^ (**Extended Data Fig. 10**). Thus, chemically distinct ligands acting at spatially segregated sites converge on a shared gating apparatus to modulate channel opening in heteromeric insect olfactory receptors.

### Surface complementarity at the anchor domain enables flexible assembly

A defining feature of insect olfactory receptors is their modular architecture, in which a single obligate co-receptor, Orco, can assemble with tens or hundreds of highly divergent tuning OR subunits within a species^23^. Given the functional interchangeability of Orcos across insect lineages^30^, Orco must potentially accommodate millions of distinct OR sequence variants. What structural features endow Orco with this remarkable capacity to assemble and function with such diverse OR partners?

Inspection of the *Ab*Orco-*Cp*OR9 structure reveals that intersubunit contacts are highly concentrated within the intracellular anchor domain, where neighboring subunits display striking shape complementarity (**Fig. 3a**). A primary contact interface is centered on a Trp–His–Tyr (WHY) motif divided across the subunit interface (**Fig. 3a; Extended Data Fig. 11a**). These residues comprise the most conserved sequence element across both OR and Orco anchor domains (**Extended Data Fig. 11c**). At the cytosolic end of S6, the polypeptide chain is partially unravelled, exposing a bulky tryptophan side chain. Four residues away, a conserved tyrosine projects toward the adjacent subunit to form a hydrogen bond with a buried histidine (**Fig. 3b**). Together, these conserved residues interlock into a complementary hydrophobic groove formed by S5 and S7a of the neighboring subunit. This interdigitating arrangement allows divergent ORs and Orco to fit together like complementary puzzle pieces, suggesting that coupling specificity relies on the conserved geometry of complementary surfaces rather than a rigid, sequence-specific network of side-chain interactions.

**Fig. 3.**
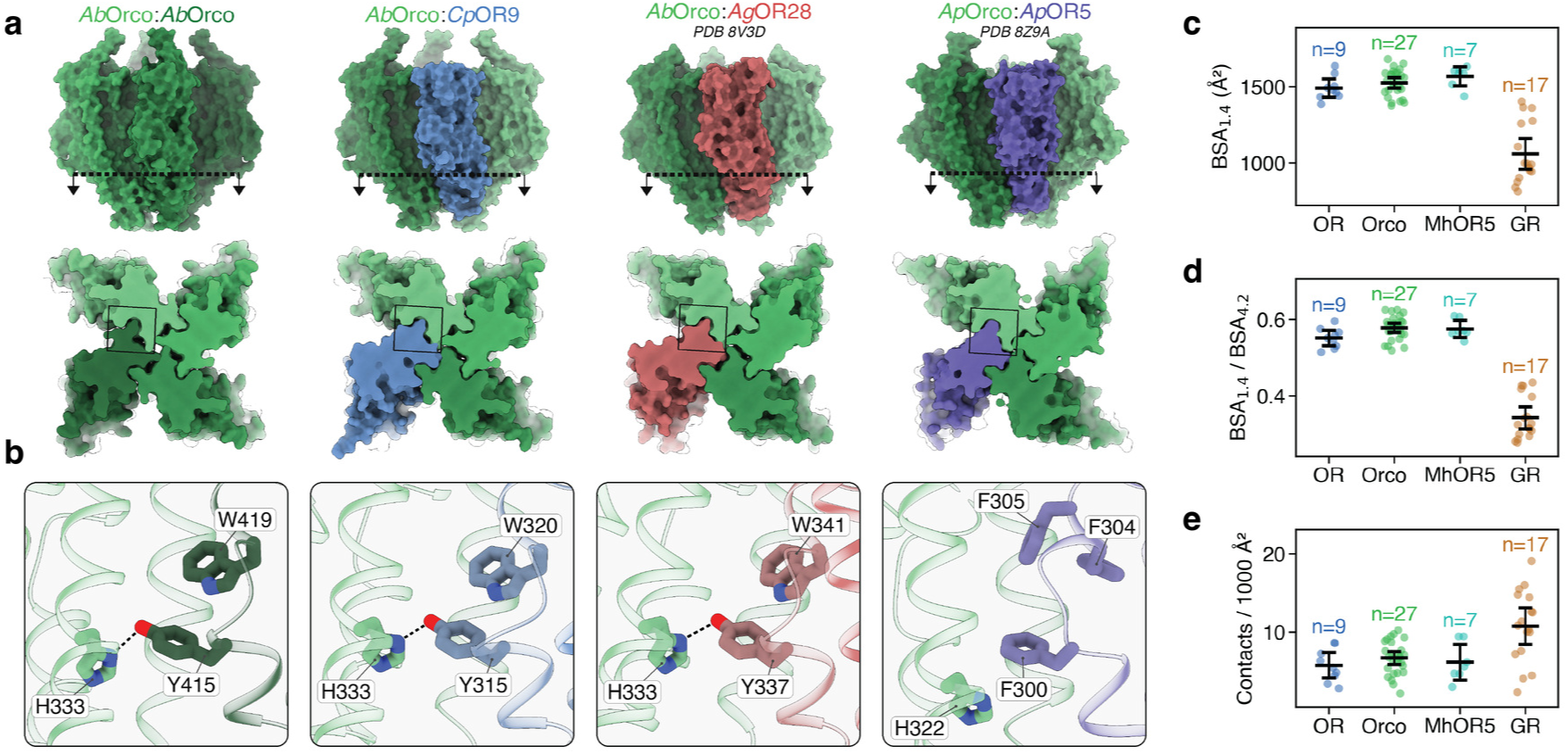
Assembly of ORs and GRs is mediated by shape complementarity in the anchor domain. **a**, Surface representations of homo- and heteromeric receptors, viewed from the side (top row) and from the cytoplasm (bottom row) after sectioning through the anchor domain at the level of the Trp-His-Tyr (WHY) motif, indicated by the dashed line at top. Shown from left: *Ab*Orco homotetramer (this study), *Ab*Orco-CpOR9 heteromer (this study), *Ab*Orco-*Ag*OR28 heteromer (PDB: 8V3D), and *Ap*Orco-*Ap*OR5 (PDB: 8Z9A). In all structures, Orco subunits are colored shades of green. **b**, Zoomed-in view of boxed region in (a) showing the packing interactions between a single Orco-Orco or Orco-OR interface. Residues corresponding to the conserved WHY motif across subunits are shown in stick representation. **c**, Jitter plot of buried surface area (BSA) measured with a water probe (1.4 Å) for all unique states of published heteromeric ORs (blue), Orcos (green), basal OR (*Mh*OR5; teal), and GRs (brown). S7b is excluded from these measurements to isolate anchor domain contacts. Each dot represents a unique chain with group mean and 95% CI shown as lines and hashes. n represents the number of unique chains measured. **d**, Jitter plot of the ratio of BSA measured with fine (1.4 Å) and coarse (4.2 Å) probe to highlight surface convolution for the same receptor groups. Each dot represents a unique chain with group mean and 95% CI shown as lines and hashes. n represents the number of unique chains measured. **e**, Jitter plot of the number of electrostatic contacts per chain (hydrogen bonds and salt bridges) normalized to surface area for the same receptor groups. n represents the number of unique chains measured. The structures analyzed and methodology for **c**, **d**, and **e** are listed in the Methods.

This mode of assembly contrasts with GRs, which form homo- or heteromers^41^, but need not assemble with so many divergent partners. Based on the structures of homomeric GRs^19,20^, the corresponding region of the anchor domain remains helical and neighboring subunits interact in a planar fashion, via specific electrostatic interactions (**Extended Data Fig. 11b**). Surface analysis using either a fine (1.4 Å) or coarse (4.2 Å) probe revealed that Orco-OR interfaces are significantly more convoluted than those of GRs, with the WHY motif contributing substantially to this increased surface complexity (**Fig. 3d**). ORs and Orcos also show >40% greater buried surface area within the anchor domain, whereas GRs exhibit a higher density of electrostatic contacts, consistent with a more geometrically-constrained interaction surface (**Fig. 3c, e**). The basal receptor *Mh*OR5 possesses an incomplete WHY motif, which contributes to a highly interdigitated interface that achieves a similarly large contact area through a distinct convoluted geometry (**Extended Data Fig. 11b**). This partially formed motif may represent an ancestral solution for subunit packing that preceded the emergence of the more stereotyped anchor domain interactions found in Orco-OR heteromers^11,23^.

Together, these observations suggest that Orco-OR assembly is driven primarily by shape complementarity and distributed van der Waals packing, with the WHY motif providing a conserved structural feature that supports interactions with divergent ORs. Consistent with this role, mutation of the conserved tryptophan within the WHY motif in *Drosophila* Orco impairs both receptor trafficking and function^42^, underscoring the importance of this interface for tetramer assembly.

### Orco–OR complexes adopt multiple stoichiometries

If interface recognition is mediated by shape complementarity rather than geometrically constrained polar interactions, heteromeric receptors may be capable of assembling in multiple stoichiometries. Although single-molecule photobleaching of purified heteromers previously suggested variable subunit composition^16^, all structures to date have exclusively resolved 3:1 Orco-OR complexes^16,17^. We therefore used split-GFP complementation^43,44^ as a readout of subunit proximity to assess the range of *Ab*Orco-*Cp*OR9 receptor assemblies formed in our heterologous expression system. We fused either GFP_1–10_ (comprising the first ten strands of the GFP β-barrel) or GFP_11_ (the eleventh strand), to the N-terminus of *Ab*Orco or *Cp*OR9 via a flexible 13-amino-acid linker, such that fluorescence reconstitution should be limited to subunits within a tetramer. Consistent with this structural constraint, co-expression of *Ab*Orco and *Bm*GR9^19^, a homomeric fructose receptor from *Bombyx mori*, yielded no detectable fluorescence, despite both receptors being highly expressed and exhibiting robust complementation within their cognate homotetramers (**Fig. 4a**). Conversely, strong fluorescence reconstitution was observed when *Ab*Orco and *Cp*OR9 were tagged in either reciprocal configuration, confirming intersubunit complementation within heteromers. Critically, co-expression of *Cp*OR9–GFP_1–10_ and *Cp*OR9–GFP_11_ alongside untagged *Ab*Orco also yielded robust fluorescence, demonstrating that heteromeric complexes containing at least two tuning OR subunits assemble in HEK cells.

**Fig. 4.**
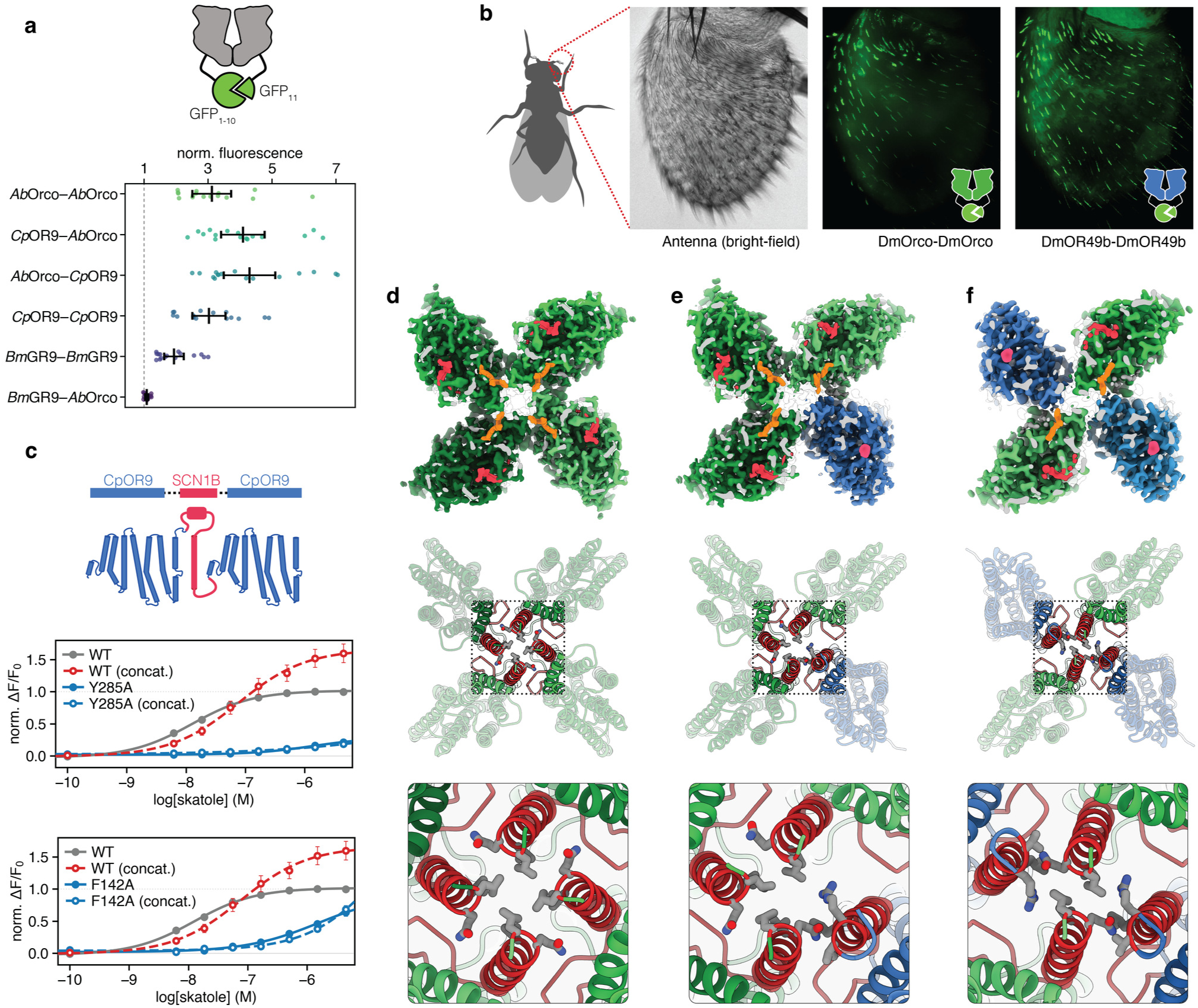
Assembly, function, and structure of alternative heteromeric stoichiometries. **a**, Schematic of the GFP complementation paradigm (top) in which receptors fused to complementary GFP_1- 10_ or GFP_11_ fragments generate fluorescence upon close association (schematic, top). Relative fluorescence of indicated combinations of *A. bakeri* Orco (*Ab*Orco), *C. pipiens* OR9 (*Cp*OR9) and *B. mori* GR9 (*Bm*GR9) normalized to untransfected HEK293T cells on the same plate. Note that *Cp*OR9-*Cp*OR9 constructs were also co-transfected with non-fluorescent *Ab*Orco. Each point represents an independent well; black bars indicate mean ± 95% CI. **b**, Split-GFP complementation in the *Drosophila* antenna reveals association between copies of *Dm*Orco and between copies of the indolergic tuning receptor *Dm*OR49b in native olfactory sensory neurons. Left, bright-field image of the antenna; middle and right, GFP fluorescence for the indicated receptor pairs from top planes of the antenna highlighting sensilla. **c**, Functional analysis of concatenated *Cp*OR9 subunits. Top, schematic of two *Cp*OR9 subunits joined through the single-pass transmembrane protein SCN1B. Dose–response curves for concatenated *Cp*OR9 (dashed lines) or monomeric *Cp*OR9 (solid lines) co-expressed with monomeric *Ab*Orco, comparing the WT receptor (gray and red lines) to either Y285A or F142A mutant *Cp*OR9 (blue lines). Note that in concatenated mutants only one of the two binding pockets has been mutated and the other remains wild-type. Values are mean ± 95% CI with *n*=24-66. **d**–**f**, Top row: Slice through cryo-EM density at height of ligands for 4:0 *Ab*Orco homomer (**d**), 3:1 *Ab*Orco-*Cp*OR9 (**e**) or 2:2 *Ab*Orco-*Cp*OR9 (**f**). Middle row: Corresponding top view shown as cartoon with S7b helices highlighted in red. Bottom row: Zoomed in view of the gate with both hydrophobic and hydrophilic gating residues shown as gray sticks.

To determine whether heteromers containing multiple tuning OR subunits also form in their native physiological environment, we adapted the split-GFP assay to *Drosophila* olfactory sensory neurons. While split-YFP complementation experiments between Orco and *Dm*OR43a previously suggested that heteromers can incorporate multiple copies of either subunit *in vivo*^14^, whether this represents a general property of tuning ORs is not known. We therefore generated transgenic flies in which *D. melanogaster* Orco *Dm*Orco and *Dm*OR49b^45^, the fly ortholog of *Cp*OR9, were tagged with either GFP_1–10_ or GFP_11_ and expressed in the majority of olfactory sensory neurons under the *Orco* promoter^10^. Robust reconstituted fluorescence was observed in sensory dendrites and soma for both Orco–Orco and OR-OR configurations (**Fig. 4b; Extended Data Fig. 12**). Thus, heteromeric receptors do not appear to be restricted to a fixed 3:1 stoichiometry *in vivo* but can incorporate multiple copies of either subunit, consistent with the flexible assembly permitted by shape complementarity at the anchor-domain interface.

### Concatenated ORs are functional

The existence of multiple heteromeric stoichiometries raises a central question: which assemblies can support odorant signaling? Because freely associating monomeric subunits generate a mixed population of receptor compositions, functional responses cannot be assigned to a defined stoichiometry without constraining assembly. We therefore covalently tethered two OR subunits by inserting the single-pass transmembrane protein SCN1B (the structured β-subunit of the voltage-dependent sodium channel)^46^ between the extracellular C-terminus of one *Cp*OR9 subunit and the intracellular N-terminus of a second (**Fig. 4c**). AlphaFold3 predictions indicate that this concatenated *Cp*OR9 construct is compatible with a 2:2 heteromeric assembly containing two individual Orco subunits (**Extended Data Fig. 13**). Consistent with this, co-expressing *Cp*OR9-*Cp*OR9 concatemers with monomeric Orco produced robust skatole-evoked responses, albeit with a modest decrease in potency (log EC50 shifted from -7.8 to -7.0) (**Fig. 4c**). Thus, heteromeric complexes constrained to a 2:2 Orco–OR stoichiometry are functional.

To determine whether *Cp*OR9–*Cp*OR9 concatemers can signal through a single functional OR subunit, which would mimic the architecture of a 3:1 complex containing only one OR, we introduced single point mutations into the binding pocket of one subunit within the linked pair. We selected Y285A, which nearly abolishes *Cp*OR9 activity, and F142A, which strongly attenuates but does not eliminate signaling (**Fig. 1k**). In each case, the single-mutant concatemer recapitulated the functional profile of the monomeric mutant receptor, despite the presence of an intact *Cp*OR9 in the complex (**Fig. 4c**). Thus, within heteromers constrained to contain two ORs, activation appears to depend on the intact functional capacity of both subunits. These data establish that the 2:2 heteromer is a functionally competent assembly but reveal that its activation depends on productive signaling in both OR subunits—either because both must bind odorant or because mutation of one subunit disrupts the concerted conformational changes needed to open the tetrameric gate.

### Cryo-EM captures additional Orco-OR stoichiometries

Evidence that heteromers containing two OR subunits are functional prompted us to re-examine the cryo-EM dataset from which our 3:1 structure was determined in search of additional stoichiometric assemblies. By embedding each particle into a latent ’variability’ space^47^ derived from the consensus image stack, two additional sub-populations emerged: an Orco homomer that co-purified alongside the heteromer and a 2:2 *Ab*Orco–*Cp*OR9 complex. We reseeded these additional references for further classification^48^ (**Extended Data Fig. 1**), yielding three discrete stoichiometries: an Orco homomer at 2.1 Å, the 3:1 heteromer at 2.2 Å, and a 2:2 heteromer at 2.8 Å (**Fig. 4d–f**). In the 2:2 configuration (**Fig. 4f**), the two *Ab*Orco and two *Cp*OR9 subunits alternated around the central pore in C2 symmetry and we did not detect any particles in which the two ORs were adjacent to each other. Although the 2:2 assembly represented only a minor fraction of particles, its recovery from the same dataset as the 3:1 structure further underscores the capacity of ORs and Orcos to flexibly assemble into multiple stoichiometric arrangements in our heterologous expression system. The relative abundance of each class in the cryo-EM dataset likely reflects differential stability during detergent purification rather than their true distribution *in vivo*, where the specialized lipid environment of the sensory neuron may better stabilize otherwise labile receptor assemblies. Consistent with this, *Cp*OR9 and the orthologous indolergic receptor from *D. virilis*, *Dv*OR49b, reliably assembled as homotetramers when heterologously expressed in HEK cells in the absence of Orco, but proved recalcitrant to further biochemical purification (**Extended Data Fig. 14**). Thus, while the anchor domain supports a broad range of receptor assemblies–including homotetramers of both Orco and tuning ORs–which complexes are recovered after detergent solubilization is likely strongly shaped by their dependence on the lipid environment and sensitivity to membrane disruption^15^.

In the 2:2 structure, the indole-bound *Cp*OR9 subunits and VUAA4-bound *Ab*Orco subunits closely resembled their counterparts in the 3:1 heteromer, with individual OR and Orco protomers superimposable at Cα RMSD less than 1 Å (**Extended Data Fig. 8**). A notable difference, however, was the arrangement of subunits within the tetramer. As the two *Cp*OR9 subunits adopted activated conformations, the opposing Orco subunits tilted inward as rigid-bodies toward the pore’s central axis, occupying the space created by the outward displacement of the ligand-bound ORs. This repositioning brings the two Orco subunits ∼6 Å closer together than in the 3:1 heteromer, allowing them to fully occlude the ion-permeation pathway. Thus, although receptors constrained to a 2:2 stoichiometry are functional (**Fig. 4c**), the 2:2 structure represents a nonconductive state in which activated OR subunits are insufficient to open the pore (**Fig. 4f**) unless their conformational changes propagate cooperatively across the tetramer.

The same dataset also yielded a high-resolution *Ab*Orco homotetramer, with VUAA4 occupying each of the four membrane-facing crevices formed between S5 and S7b. Although Orco homomers can function as autonomous ion channels when activated by this synthetic agonist^22^, the conformation of Orco subunits in the homotetramer also remained essentially unchanged. Thus, every Orco structure captured here–representing nine unique chains in total, in apo and VUAA4-bound states and in multiple stoichiometries–was virtually superimposable (**Extended Data Fig. 8**) and displayed none of the pore-domain rearrangements that accompany opening in ORs or GRs (**Extended Data Fig. 2**). These data suggest that all Orco-OR heteromeric structures captured to date^16,17^ are unlikely to represent fully activated channels, but instead correspond to snapshots of inactive, desensitized, or intermediate states along the gating pathway.

### A common conformational landscape links homomeric and heteromeric receptors

The structural invariance of Orco raises the possibility that this co-receptor has evolved to function primarily as a rigid structural scaffold to support divergent ORs. Alternatively, existing heteromeric structures may capture only a subset of the conformations sampled during receptor gating. To distinguish between these possibilities, we turned to *Mh*OR5^12^, a broadly tuned homomeric receptor, reasoning that its high basal activity might allow multiple states along the gating pathway to be resolved within a single sample. Visualizing the ligand-free conformational ensemble of *Mh*OR5 could therefore reveal intermediate states that are challenging to capture in more tightly regulated heteromers and provide a basis for interpreting their nonconductive conformations.

From a single dataset of *Mh*OR5 purified in the absence of exogenous ligand, we produced a consensus set of particles at 2.4 Å and analyzed the underlying conformational variability to recover three stable states: a C4-symmetric closed state, a C2-symmetric closed state, and a C4-symmetric open state (**Fig. 5a–c, Extended Data Fig. 15**). While the C4-closed and C4-opened states resemble the described apo and odorant-bound structures of *Mh*OR5^12^, the previously unobserved C2 state was the most prevalent conformation, indicating it represents a prominent feature of the *Mh*OR5 conformational landscape rather than a rare excursion. In the C2-state, two opposing *Mh*OR5 subunits are partially activated, whereas the other two remain in a closed configuration. In the activated subunits, the S3–S6 gap is narrowed, accompanied by a small unassigned density positioned in the odorant-binding pocket, suggesting that small molecules present during purification may bind to this promiscuously tuned receptor and stabilize this intermediate state.

**Fig. 5.**
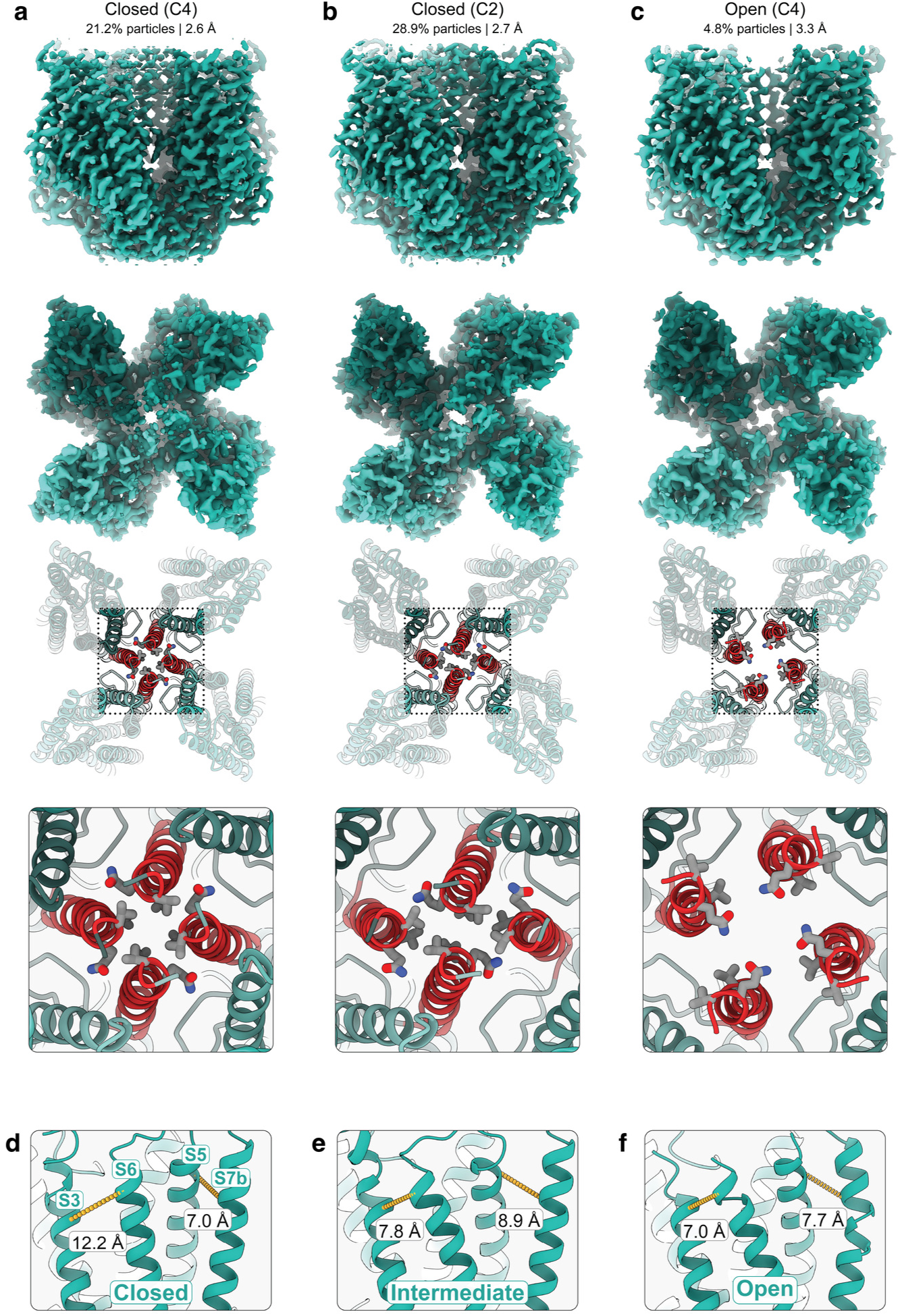
Conformational landscape of the basal homomeric receptor MhOR5. **a**, Representation of the *Mh*OR5 ligand-free C4 closed state showing side view of the cryo-EM density (row 1), extracellular view of the density (row 2), extracellular view of the tetrameric model as cartoon with S7b highlighted in red (row 3), and a zoomed in view of the gate with hydrophobic and hydrophilic gating residues shown (row 4). **b**, Same as (**a**) for *Mh*OR5 ligand-free C2 gating intermediate. **c**, Same layout as (**a**) for the *Mh*OR5 ligand-free C4 open state. **d**, **e**, **f**, Gating transitions for a single protomer from C4 closed (**d**), through an intermediate (**e**), to C4 open (**f**) conformations.

Alignment of the individual *Mh*OR5 protomers suggests a stepwise transition from the C4-closed to C4-opened state that places the observed conformations on a structural continuum. In both C4-states, the S5 and S7b helices remain tightly coupled, but the local interaction network reorganizes as S7b tilts away from the central axis (**Fig. 5a, c, d, f**), converting a narrow hydrophobic constriction at the extracellular gate into a widened, hydrophilic pore. These rearrangements broadly parallel conformational changes observed in heteromeric ORs, including *Cp*OR9 (**Extended Data Fig. 2a–c**). By contrast, the *Mh*OR5 C2-intermediate state appears to capture an intervening conformation in which the movements of the S5 and S7b helices are uncoupled (**Fig. 5d–f**). While the S5 helices of the two active subunits are shifted outward towards their open-state position, their S7b helices remain engaged at the central axis, constricting the pore (**Fig. 5e**). Thus, movement of S5 may precede displacement of S7b, creating an asymmetric intermediate in which just part of the tetramer has progressed along the gating trajectory (**Fig. 5b**). Only when all four S7b helices tilt away from the central axis does the pore form a wider hydrophilic aperture compatible with ion permeation (**Fig. 5c**). Consistent with this broader conformational flexibility, 45% of particles could not be confidently assigned to the three symmetry classes and remained poorly resolved within the pore domain, despite the high resolution of the consensus reconstruction (**Extended Data Fig. 15; Extended Data Table 3**). Individual *Mh*OR5 subunits therefore appear to sample additional asymmetric intermediates between these more stable, structurally resolved conformations.

The *Mh*OR5 C2-intermediate and 2:2 *Ab*Orco-*Cp*OR9 heteromer display striking similarity in their symmetric arrangement of partially opened and closed subunits (**Figs. 4f, 5b**). In the heteromer, the two indole-bound *Cp*OR9 subunits adopt activated conformations, whereas the two *Ab*Orco subunits remain in a closed configuration with their S7b helices occluding the ion conduction pathway. The 2:2 heteromeric structure therefore resembles an asymmetric intermediate in which ligand-induced rearrangements have occurred in only a subset of subunits but have not yet propagated across the tetramer to fully dilate the pore. More broadly, these structures suggest that homomeric and heteromeric odorant receptors sample from a common conformational landscape in which pore opening proceeds through coordinated but non-equivalent movements of individual subunits (**Fig. 6**). Structural analyses may have, thus far, captured only a subset of states within this dynamic gating process.

**Fig. 6.**
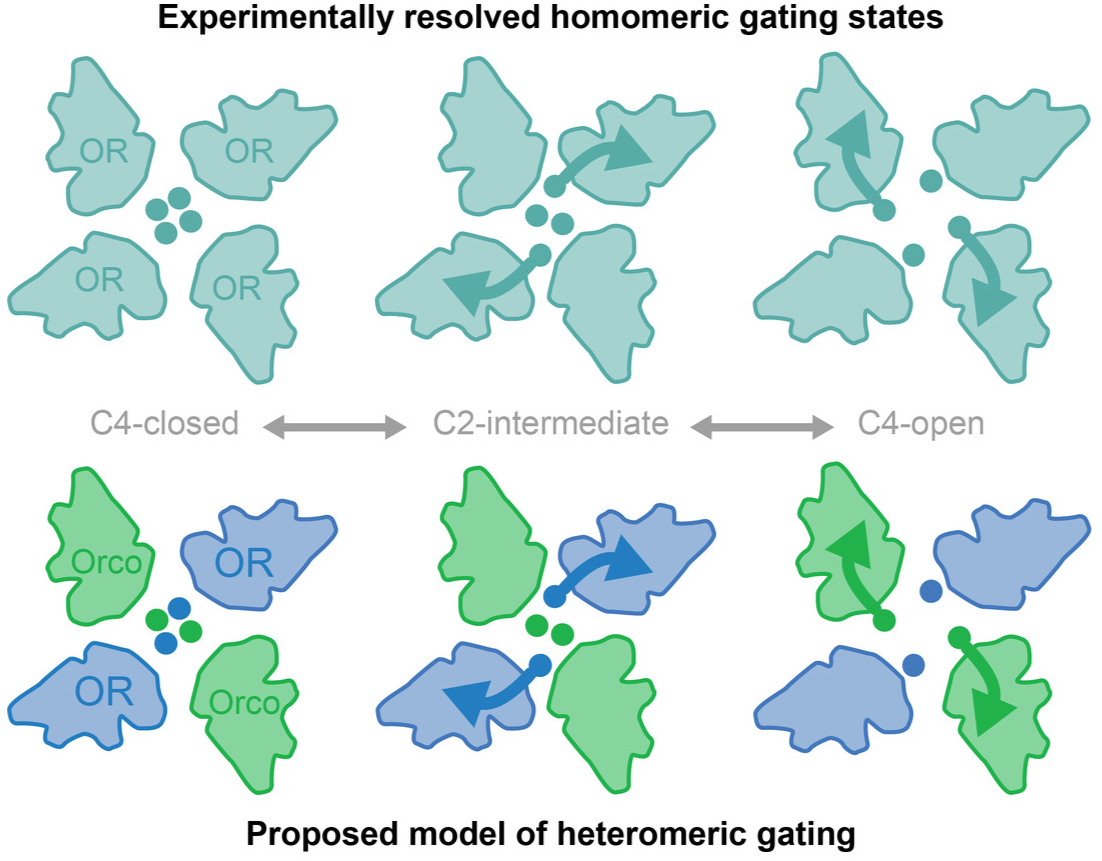
Models of homomeric and heteromeric channel opening. Comparison of experimentally derived gating states resolved for homomeric *Mh*OR5 (top row) and proposed model of heteromeric gating based on our structural analyses (bottom row).

## Discussion

Insect odorant receptors represent one of the largest and most diverse families of ion channels in nature. Their extraordinary diversification imposes two competing demands on receptor architecture: tuning must rapidly evolve to allow each species to adapt to their particular chemical niche while the machinery for assembly and ion conduction must remain reliable and robust. Our analyses show how heteromeric olfactory receptors reconcile these demands through a functional division of labor. Tuning subunits such as *Cp*OR9 govern selective odorant recognition, whereas the obligate co-receptor Orco confers stability to the heteromer, consistent with other recent structural studies of this family^16,17^. In *Cp*OR9, selective recognition arises through precise ligand coordination within a layered pocket architecture, where specific residues orient and constrain a single binding pose that discriminates between closely related odorants. By contrast, although Orco retains a canonical ligand-binding pocket, this cavity is occupied by a membrane phospholipid, providing a mechanism to prevent spurious activation by volatile odorants. Yet, Orco’s odorant insensitivity does not reflect a loss of its intrinsic capacity to gate^21^. Instead, synthetic agonists engage a distinct allosteric site near the pore, revealing a route through which Orco can directly influence gating of the heteromer. These findings suggest that Orco is not simply a conserved structural scaffold for divergent ORs, but an active participant that supports both flexible assembly and cooperative channel gating.

Orco’s ability to partner with an evolving repertoire of tuning ORs relies on an adaptable assembly architecture. Inter-subunit contacts are concentrated within the intracellular anchor domain, where adjacent subunits exhibit pronounced geometric shape complementarity. At the core of this interface lies the highly conserved Trp–His–Tyr (WHY) motif, which acts as a structural puzzle piece whose stacked Trp and Tyr residues on S6 fit into a complementary hydrophobic groove formed by the anchor domain helices S5 and S7a of the neighboring subunit. By maximizing buried surface area and distributed van der Waals packing–rather than relying on geometrically constrained, sequence-specific polar contacts– this interdigitating interface endows Orco with the capacity to pair with countless divergent OR partners without imposing rigid constraints on their amino acid sequences. Intriguingly, the electrostatic contacts that mediate assembly in GRs appear already relaxed in homomeric *Mh*OR5, despite the absence of a fully formed WHY motif. This suggests that a more permissive interface arose prior to the emergence of Orco^11^, perhaps enabling promiscuous interactions among coexpressed ancestral receptors, with subsequent evolution of the WHY motif allowing Orco to accommodate an increasingly diverse repertoire of tuning ORs.

As a consequence of the adaptable Orco-OR interface, the full range of stoichiometries appears structurally accessible, from Orco homotetramers that are insensitive to environmental odorants to OR homotetramers that lack the capacity for stable assembly and trafficking *in vivo*. Which configurations predominate, however, is likely determined by cellular and biochemical context. Assemblies purified from HEK cells may reflect both the permissive environment of this heterologous expression system and their differential stability during purification, enriching stoichiometries most tolerant of detergent solubilization. By contrast, the specialized environment of olfactory sensory neurons may impose distinct constraints through Orco-dependent stabilization^10,14^, regulated transport into sensory cilia^49–51^, and specific lipid composition^52–54^, favoring only a subset of possible assemblies. The 2:2 complex may represent one such solution, balancing chemical responsiveness with structural stability. Indeed, OR-OR complementation demonstrates that receptors containing two tuning subunits can assemble and traffic to the sensory cilia in *Drosophila*, while OR-OR concatenation establishes that this organization is functional. Although these experiments establish the viability of the 2:2 arrangement, the permissive assembly interface leaves open the possibility that other stoichiometries are also present *in vivo*, potentially broadening sensory dynamic range or enabling coincidence detection in neurons that co-express distinct ORs^55–59^.

Flexibility in subunit assembly, however, does not imply that subunits act independently during gating. In concatenated receptors constrained to a 2:2 stoichiometry, mutation of a single OR impairs signaling, indicating that activation requires coordinated engagement across subunits. This dependence may reflect a requirement for ligand binding by both ORs or for cooperative conformational changes across multiple subunits to open the shared gate that are disrupted in the mutant OR. Coordination across subunits of the tetramer does not appear to be restricted to the ORs since VUAA-derived antagonists that bind Orco suppress odorant-evoked signaling through the tuning OR^38,39^ (**Extended Data Fig. 10**). Such cooperativity is consistent with structures of homomeric ORs and GRs, in which all four pore-lining S7b helices symmetrically dilate to open the shared ion conduction pathway^12,19,20^. Yet, Orco remains stationary in all structural snapshots resolved to date, including our 2:2 assembly in which its S7b helices occlude the ion conduction pathway. These data suggest that the asymmetric displacement of a single OR subunit captured in the 3:1 structure does not represent a stable, fully open state, but may instead reflect an intermediate or a nonconductive conformation favored under the conditions used for structural analysis.

The pore architecture provides a structural basis for how such asymmetric gating intermediates arise and why they are preferentially captured. While the enmeshed anchor domain interactions rigidly fixes the four subunits in a pinwheel arrangement, intersubunit contacts within the transmembrane domain are sparse, mediated primarily through conserved residues within the signature sequence of S7b that form the tetrameric gate. Such minimal coupling may allow individual S7b helices to sample distinct conformations, giving rise to lower-symmetry intermediates such as the C2-state of *Mh*OR5 and the 2:2 *Ab*Orco-*Cp*OR9 heteromer. Moreover, the heteromeric gate would be directly exposed to the lipid bilayer rendering its conformational stability inherently sensitive to the membrane environment. The intimate interaction of the receptor gate with membrane lipids may explain why Orco has yet to be captured in an open state despite clear evidence that it gates autonomously when bound to synthetic agonists^21^. The structures of homomeric GRs have also been resolved in a closed conformation despite being bound by tastants, indicating that ligand occupancy alone may be insufficient to stabilize the fully open state in detergent-solubilized samples^20^. Capturing Orco in an open state may depend on specialized lipid constituents or biophysical properties of the ciliary membrane that are not preserved after purification, consistent with the requirement for phospholipid flippases such as ATP8B in Orco-expressing sensory neurons for odorant signaling^52,53^.

Although an open structure of Orco remains elusive, our VUAA4-bound structures nevertheless offer insight into the agonist’s binding mode and potential mechanism of action. VUAA4 occupies a membrane-accessible crevice wedged between S5 and S7b at the pivot point where these elements reorganize during gating in ORs and GRs (**Extended Data Fig. 2**), suggesting that this crevice may represent a privileged site for allosteric control across the broader chemoreceptor family. VUAA4 bends about its central thioether linkage, potentially allowing it to accommodate analogous helical rearrangements during Orco activation. VUAA4 also contacts a well-resolved annular phospholipid adjacent to the gate that participates in an intersubunit hydrogen-bond network stabilizing the closed pore of ORs, GRs, and Orcos (**Extended Data Fig. 2d**). Because these interactions are disrupted during pore opening in ORs and GRs, VUAA4 may promote Orco gating by perturbing the lipid-protein network at the gate, thereby lowering the energetic barrier to pore dilation.

The constrained geometry of the VUAA4-binding crevice provides a structural basis for the stringent structure–activity relationships of Orco agonists. Each moiety of VUAA1 and VUAA4 occupies a distinct sub-pocket, leaving little room for chemical modification, consistent with the limited tolerance for substitutions across their shared scaffold^22,38^. The requirement for the central thioether^37^ further suggests that ligand flexibility is needed to accommodate rearrangements of the S5–S7b interface. More broadly, minor modifications can convert VUAA-derived agonists into antagonists that still bind Orco but suppress odorant signaling through the tuning OR, underscoring the functional coupling of subunits within the heteromer^38,39^.

Together, our data suggest that Orco’s conformational invariance across different structural snapshots does not reflect a loss of the ancestral gating capacity retained by other members of this receptor family. Rather, Orco appears to serve as an integral component of cooperative channel gating, consistent with its strict conservation of residues in S7b signature sequence that contribute to pore opening in ORs and GRs. This modular organization provides a solution to receptor diversification, uncoupling the conserved mechanisms of assembly and gating from rapidly evolving odorant recognition, thus enabling the extraordinary diversification of insect olfactory receptors.

## Acknowledgements

We thank R. Benton, S. R. Datta, A. Haldane, T. Gao and members of the V.R. laboratory for discussions and comments on the manuscript; V. Gomes for initial design of concatenated constructs; M. Ebrahim, J. Sotiris, and H. Ng at The Rockefeller University Evelyn Gruss Lipper Cryo-Electron Microscopy Resource Center for technical support. This work was supported by the Bill and Melinda Gates Foundation (V.R. and J.A.B); NIH NIAID R01AI103171 (V.R.) and NIH NIGMS RM1GM149406 (J.A.B). V.R. is a Howard Hughes Medical Institute Investigator.

## Author contributions

V.R., J.A.B. and N.P. conceived the study, designed experiments and wrote the manuscript with input from all authors. N.P. performed biochemistry, collected and analyzed cryo-EM data, built and refined the models, and performed molecular dynamics. J.A.B. performed initial structural analyses of the VUAA4-bound receptor. D.N. performed biochemistry, designed and generated transgenic flies and complementation constructs with help from C.U., and carried out *in vitro* and *in vivo* complementation assays. D.N., B.G., C.U., M.A.Y. and J.A.R. carried out molecular biology, cell culture, and calcium imaging assays on receptor variants.

## Data availability

The cryo-EM density and models have been deposited to their respective databases under the following accessions: 4:0 *Ab*Orco homomer (ligand-free) [PDB XXXX; EMDB XXXXX]; 3:1 *Ab*Orco:*Cp*OR9 heteromer (ligand-free) [PDB XXXX; EMDB XXXXX]; 4:0 *Ab*Orco homomer (+VUAA4) [PDB XXXX; EMDB XXXXX]; 3:1 *Ab*Orco:*Cp*OR9 heteromer (+VUAA4/indole) [PDB XXXX; EMDB XXXXX]; 2:2 *Ab*Orco:*Cp*OR9 heteromer (+VUAA4/indole) [PDB XXXX; EMDB XXXXX]; MhOR5 C4 closed (ligand-free) [PDB XXXX; EMDB XXXXX]; *Mh*OR5 C2 closed (ligand-free) [PDB XXXX; EMDB XXXXX]; *Mh*OR5 C4 open (ligand-free) [PDB XXXX; EMDB XXXXX]. Cryo-EM data acquisition, processing, and model refinement statistics are provided in Extended Data Tables 1–3. Analysis of raw GCaMP dose-response data are provided in Extended Data Figs. 16–25.

## Code availability

The dose-response analysis code in this study are available from the corresponding authors on request.

## Methods

### Protein purification: ligand-free *Cp*OR9:*Ab*Orco

The apo *Ab*Orco-*Cp*OR9 heteromer was produced by baculovirus (BacMam)^60^ expression in HEK293S GnTI⁻ cells (ATCC CRL-3022) grown in FreeStyle 293 medium (Thermo Fisher Scientific). Cells were transduced with baculovirus encoding N-terminally mVenus-tagged *Apocrypta bakeri* Orco (AbOrco) and N-terminally sfGFP-tagged *Culex pipiens* CpOR9, supplemented with 4.5 mM sodium valproate (Sigma-Aldrich), grown at 37 °C, and harvested 48-72 h post-infection. Frozen pellets from 3 L of culture were thawed in 100 mL of solubilization buffer [2% (w/v) glyco-diosgenin (GDN; Anatrace), 300 mM NaCl, 20 mM HEPES pH 7.5, 1 mM PMSF, 4 µg/mL aprotinin, 2 µg/mL leupeptin, 2 µM pepstatin A, 2 µM E-64, and DNase I] and solubilized for 1-2 h at 4 °C with gentle stirring. The lysate was clarified by ultracentrifugation and batch-bound to GFP-nanobody affinity resin for 3-4 h at 4 °C. The resin was transferred to a column and washed with 200 mL of gel-filtration buffer [150 mM NaCl, 50 mM Tris-HCl pH 8.0, 0.005% (w/v) GDN, 0.1 mg/mL lipids, 2 mM DTT]; the lipid supplement was a 5:1:1:5 (w/w) mixture of POPC:POPE:POPS:brain total lipid extract (Avanti Polar Lipids), pre-solubilized at 10 mg/mL via sonication in 35 mM CHAPS (Anatrace), 150 mM NaCl, 50 mM Tris pH 8.0 before dilution into the buffer. Protein was released from the immobilized mVenus/sfGFP-nanobody complex by on-resin cleavage with HRV 3C protease, concentrated, and polished by size-exclusion chromatography on a Superose 6 Increase 10/300 GL column (Cytiva) on an ÄKTA system in the same gel-filtration buffer. Peak fractions were pooled and concentrated to ∼10 mg/mL for grid preparation.

### Protein purification: ligand-bound *Cp*OR9:*Ab*Orco

Expression and affinity capture of the ligand-bound *Ab*Orco-*Cp*OR9 heteromer were performed as for the apo complex (HEK293S GnTI⁻; mVenus-*Ab*Orco + sfGFP-*Cp*OR9), except that all buffers were supplemented with indole and VUAA4. Ligands were dissolved into the pre-mixed buffer base to nominal concentrations of 1 mM indole and 100 µM VUAA4. Frozen pellets from 3 L of culture were thawed in 100 mL of solubilization buffer [2% (w/v) GDN (Anatrace), 100 mM NaCl, 100 mM KCl, 50 mM Tris-HCl pH 8.0, 3 mM CaCl_2_, 3 mM MgCl_2_, 1 mM indole, 100 µM VUAA4, 2 mM DTT, 1 mM PMSF, 4 µg/mL aprotinin, 2 µg/mL leupeptin, 2 µM pepstatin A, 2 µM E-64, 1 mM benzamidine, and DNase I] and solubilized for 1-2 h at 4 °C with gentle stirring. The clarified lysate was batch-bound to GFP-nanobody resin, and the resin was washed on a column with gel-filtration buffer [100 mM NaCl, 100 mM KCl, 50 mM Tris-HCl pH 8.0, 3 mM CaCl_2_, 3 mM MgCl_2_, 1 mM indole, 100 µM VUAA4, 0.1 mg/mL lipids (5:1:1:5 POPC:POPE:POPS:brain total lipid extract; Avanti Polar Lipids), 2 mM DTT, and 0.02% (w/v; ∼170 µM) GDN carried over from the GDN-solubilized lipid stock]. Lipids were pre-solubilized at 10 mg/mL via sonication in high-detergent extraction buffer before dilution. Protein was liberated by on-resin HRV 3C protease cleavage, concentrated, and polished on a Superose 6 Increase 10/300 GL column (Cytiva) on an ÄKTA system in the gel-filtration buffer. Peak fractions were concentrated to ∼10 mg/mL for grid preparation.

### Protein purification: ligand-free *Mh*OR5

*Machilis hrabei* OR5 (MhOR5) was expressed by baculovirus (BacMam) infection of HEK293F cells (Thermo Fisher Scientific) grown in SMM293-TII medium (Sino Biological), using the baculovirus described previously ^12^. Cells (2.8 L) were harvested by centrifugation, washed in PBS, and lysed in 1% (w/v) n-dodecyl-β-D-maltoside (DDM; Anatrace), 0.2% (w/v) cholesteryl hemisuccinate (CHS; Anatrace), 100 mM NaCl, 100 mM KCl, 1 mM PMSF, 1x Halt Protease Inhibitor Cocktail (Thermo Fisher Scientific), and DNase I (Sigma-Aldrich). The lysate was clarified by centrifugation and bound to GFP-nanobody affinity resin. The resin was washed on a column with 100 mL of 100 mM NaCl, 100 mM KCl, 20 mM HEPES pH 7.1, 0.02% (w/v) DDM, 0.004% (w/v) CHS, 0.1 mg/mL lipids (5:1:1:5 POPC:POPE:POPS:brain total lipid extract; Avanti Polar Lipids), and 2 mM DTT. Lipids were pre-solubilized at 10 mg/mL via sonication in 35 mM CHAPS (Anatrace), 150 mM NaCl, 50 mM Tris pH 8.0 before dilution into the buffer. Protein was eluted by HRV 3C protease cleavage of the LEVLFQGP site immediately C-terminal to the N-terminal superfolder-GFP tag, and polished on a Superose 6 Increase 10/300 GL column (Cytiva) on an ÄKTA system, which produced a single sharp, monodisperse peak. Peak fractions were concentrated to ∼10 mg/mL for grid preparation.

### Cryo-EM data processing

Data collection and image processing were performed using the same workflow previously described^48^. The strategy is summarized here; dataset-specific acquisition and refinement statistics are reported in Extended Data Tables 1–3.

Movies were acquired on a Titan Krios (CEMRC), using a multi-shot, multi-hole beam-image-shift strategy^61^ such that exposures could subsequently be assigned to discrete optical groups for aberration and CTF refinement. Movies were motion-corrected and dose-weighted, and per-micrograph CTF parameters were estimated in cryoSPARC^62^.

Particles were selected by first generating a small set of high-fidelity picks to train a Topaz model^63^, then curating the resulting particles by iterative decoy classification without imposed symmetry, which removes damaged particles, contaminants, and empty micelles while retaining rare compositional states^48^. Curated particles were refined by non-uniform refinement^62^. Enforced C4 symmetry gave the highest-resolution alignment and was therefore used for reference-based Bayesian polishing in RELION^64^. Polished particles were divided into their optical groups for CTF refinement^65^ in cryoSPARC, and the symmetric consensus map was symmetry-expanded and treated in C1 for all subsequent heterogeneity analysis. Performing classification and variability analysis on symmetry-expanded particles allows symmetric signals to be multiplied when they are present without limiting identification of asymmetric states.

Conformational and compositional heterogeneity was resolved by cryoSPARC 3D variability analysis (3DVA)^47^ with 3 Å low-pass filtering across 3 variability modes, clustering the latent space into discrete states along each mode. Whole-tetramer local refinements along the relevant modes yielded the final maps used for model building and analysis. For the 2:2 and 4:0 AbOrco:CpOR9 states, specifically, the maps derived from this approach were reseeded for a second round of decoy classification yielding significantly greater particle numbers and clarity for the rare stoichiometries. Final maps were sharpened with density modification^66^, and models were built iteratively in coot^67^, ISOLDE^68^, and Phenix^69,70^ using standard practices.

### Molecular dynamics simulations

Two systems were simulated: the 3:1 *Ab*Orco-*Cp*OR9 heterotetramer with indole or skatole in the *Cp*OR9 pocket and VUAA4 in each *Ab*Orco crevice, and the 4:0 *Ab*Orco homotetramer with four VUAA4. Starting coordinates came from the cryo-EM structures, with all modeled lipids and cholesterol deleted. Ligands were parameterized with GAFF2/AM1-BCC^71–73^. Each complex was embedded in a POPC bilayer with packmol-memgen^74^, solvated with TIP3P water, and neutralized to 150 mM NaCl. Proteins used AMBER ff14SB (heteromer)^75^ or ff19SB (homomer)^76^, lipids Lipid21^77^; hydrogen-mass repartitioning enabled a 4-fs timestep^78^.

Systems were energy-minimized and equilibrated through an eight-stage restrained protocol (∼3.25 ns) with position restraints released stepwise, then run unbiased in the NPT ensemble in GROMACS 2023.2 (4-fs timestep, velocity-rescaling thermostat at 310 K, Parrinello-Rahman barostat at 1 bar, PME, 1.0-nm cutoffs, LINCS on bonds to hydrogen)^79^, writing coordinates every 20 ps. Replicates were launched with randomized initial velocities: six 100 ns replicates per odorant for the heterotetramer and four 300 ns replicates for the homotetramer.

Odorant pose in the *Cp*OR9 pocket was scored per frame (MDAnalysis)^80^ by two distances: *Cp*OR9 Ser139 to the odorant ring nitrogen (indole N1 or its skatole equivalent), and the odorant 3-position ring carbon (the methyl-bearing carbon in skatole) to the CpOR9 Ala77 side chain. Annular versus bulk POPC mobility was compared in the outer leaflet of the four homotetramer trajectories. Annular lipids were the two nearest outer-leaflet POPC to each of the four Q472 gates by headgroup center-of-mass distance, resolved to eight non-overlapping lipids per replicate; a count-matched bulk set was drawn from the outermost 15% lateral shell with ≥10 Å spacing.

### Anchor domain analysis

Buried surface area (BSA) and polar-contact analyses were performed in UCSF ChimeraX 1.11^81^ on all PDB-deposited heteromeric insect OR and GR structures and the new additions from this work, with pore domain (S7b) and non-protein heteroatoms removed and polar hydrogens added. ORs (PDB code): *A. aegypti* OR10 (8V00, 8V02); *A. gambiae* OR28 (8V3C, 8V3D); *A. pisum* OR5 (8Z9A, 8Z9Z); *C. pipiens* OR9 (this work); *M. hrabei* OR5 (this work). Orcos (PDB code): *A. bakeri* Orco (8V00, 8V02, 8V3C, 8V3D, this work); A. pisum Orco (8Z9A, 8Z9Z). GRs (PDB code): *B. mori* Gr9 (8UVT, 8UVU, 8VV3, 8VC1, 8VC2); *D. melanogaster* Gr43a (8JM9, 8JMA); *D. melanogaster* Gr43a I418A (8X82, 8X83, 8X84); *D. melanogaster* Gr64a (8JME, 8JMH, 8JMI, 8ZE0, 8ZE2); D. mojavensis Gr43a (8ZDZ, 8ZE3). For every unique chain, buried surface area was computed against the remainder of the protein at probe radii of 1.4 Å and 4.2 Å, with electrostatic contacts (hydrogen bonds calculated with ChimeraX *hbonds* and salt-bridges within 4 Å) counted and normalised to BSA_1.4_ per 1000 Å^2^.

### Split GFP constructs

Gene fragments encoding TwinStrepII–split GFP_1–10_ followed by a GGLEVLFQGPGRA linker containing the rhinovirus 3C protease recognition sequence, and FLAG–split GFP_11_ followed by the same linker, were synthesized (GeneScript) and cloned in-frame into the BspDI and AscI restriction sites of pCAG vectors containing either *Ab*Orco, *Cp*OR9, or *Bm*GR9^19^. The split GFP sequences (GFP_1–10_ and GFP_11_) were based on the previously described superfolder GFP split system^43^ .

### Concatenated receptor constructs

To generate plasmids expressing concatenated *Cp*OR9 and/or *Ab*Orco, synthetic gene fragments encoding *Cp*OR9–ENLYFKSGGGS–SCN1B(1–218)–GGGSENLYFKSPPGRA or *Ab*Orco–ENLYFKSGGGS–SCN1B(1–218)–GGGSENLYFKSPPGRA were designed based on the previously reported Orco-Orco concatemer^46^ . The gene fragments were synthesized (GeneScript) and cloned into the AscI site at the 5′ end of *Cp*OR9 or *Ab*Orco in pME18st vectors already containing one copy of the corresponding receptor.

### Site-directed mutagenesis

Point mutations were introduced by PCR-based site-directed mutagenesis using complementary oligonucleotide primer pairs containing the desired nucleotide substitutions (Integrated DNA Technologies). PCR amplification of the mutant plasmid was performed using PfuTurbo DNA Polymerase (Agilent Technologies). The parental plasmid template was digested with DpnI before transformation into competent E. coli cells. All mutations were confirmed by whole-plasmid sequencing (Plasmidsaurus or Genewiz). For concatenated constructs, mutations were first introduced into the pME18st vector containing a single copy of CpOR9. After sequence verification, the mutated fragment was transferred into the corresponding concatenated construct by Gibson assembly.

### Cell-based GCaMP fluorescence calcium flux assay

Receptor function was assessed using a GCaMP6s-based calcium fluorescence assay in HEK293 cells, adapted from previously described protocols^15^. Briefly, cells were maintained in high-glucose DMEM with 10% (v/v) FBS and 1% (v/v) GlutaMAX at 37°C in 5% CO_2_, transiently co-transfected with GCaMP6s (Addgene #40753) and the receptor construct(s) of interest using Lipofectamine 2000 (Invitrogen), and plated into 384-well CELLSTAR plates (Greiner). All receptor constructs were cloned into a modified pME18ST vector without a fluorescent marker; for heteromeric receptors, Orco and OR plasmids were co-transfected at a 1:1 ratio while holding total receptor DNA constant. Fluorescence was recorded on a Hamamatsu FDSS μCell plate reader (470 nm excitation/540 nm emission) for 30 sec prior to ligand addition and 120 sec after, allowing most responses to reach a plateau. ΔF/F₀ was calculated for each well using the mean fluorescence during the 30 sec preceding ligand addition as F₀ and the mean fluorescence during the final 10 sec as F. Each 384-well plate (biological replicate) contained an *n*=4 technical replicates per titration. Custom Python code was written to analyze the resulting data, fitting a four-parameter Hill model and performing bootstrap resampling of the well-level data to produce 95% confidence bounds for the fitted parameters. The reported *n* measures well-level counts for total biological and technical replicates per titration.

### Generation of transgenic flies

Plasmids used for generating transgenic split GFP flies were constructed by synthesizing codon-optimized gene fragments encoding TwinStrepII–split GFP_1–10_, FLAG–split GFP_11_, *Drosophila melanogaster* Or49b, and *Drosophila melanogaster* Orco (GeneScript). These fragments were cloned into the XhoI and XbaI sites of the pJFRC7-20XUAS-IVS vector using Gibson assembly.

Transgenic flies were generated by ΦC31 integrase-mediated transgenesis (BestGene Inc.) into distinct attP sites. Constructs expressing GFP_1–10_-tagged *Dm*Or49b or *Dm*Orco were inserted into attP2 (Bloomington stock #8622), whereas constructs expressing GFP_11_-tagged *Dm*Or49b or *Dm*Orco were inserted into the VK27 attP site (Bloomington stock #9744). Transformants were selected and balanced over TM6b,Tb.

To generate flies expressing complementary split GFP fragments, third chromosome recombinants were generated and subsequently crossed to Orco-Gal4 flies (Bloomington stock #26818) to drive expression in all olfactory sensory neurons of the antennae and maxillary palps^3^ .

The resulting experimental genotypes were:

**Orco-Orco:** yw; Orco-Gal4/*CyO*; P{20xUAS-sGFP_1-10_::*Dm*Orco}attP2, P{20xUAS-sGFP_11_::*Dm*Orco}attVK27/TM6B

**OR-OR:** yw; Orco-Gal4/CyO; P{20xUAS-sGFP_1-10_::*Dm*OR49b}attP2, P{20xUAS-sGFP_11_::*Dm*OR49b}attVK27/TM6B

**GFP-*Ab*Orco** flies used as a positive control for split-GFP complementation: w; P{20xUAS-sfGFP::*Ab*Orco}attP40,Orco-Gal4/CyO; TI{w[+mW.hs]=TI}Orco[1]/TM6B.

### Whole-mount antennal imaging

Split GFP complementation in transgenic flies was examined by imaging whole-mount antennae at the Rockefeller University Bio-Imaging Resource Center using an instant structured illumination microscope (iSIM). Because cuticular autofluorescence increased with age and interfered with GFP detection, only newly eclosed adult female flies were used. Antennae were dissected in pH 7.5 buffered saline containing 108 mM NaCl, 5 mM KCl, 5 mM HEPES, 5 mM trehalose, 10 mM sucrose, 4 mM NaHCO_3_, 1 mM NaH_2_PO_4_, 2 mM CaCl_2_, and 8.2 mM MgCl_2_, washed twice in PBT (0.03% Triton X-100 in PBS), mounted in SlowFade Glass Antifade Mountant (Thermo Fisher Scientific, Cat. No. S36917) and covered with a #1.5 (18×18 mm) coverslip.

All images were acquired under identical conditions using a VisiTech/BioVision/Leica/Mizar iSIM microscope equipped with a 63×/1.3 NA glycerol immersion objective. GFP fluorescence was excited using a 488-nm laser and collected through a 525-nm emission filter with 70% laser power and 100-ms exposure time. Z-stacks were acquired with a step size of 100 nm using an ORCA Fusion sCMOS camera (Hamamatsu) controlled by VisiView software.

Image stacks were 3D-deconvolved in Huygens Professional (Scientific Volume Imaging, Hilversum, NL) using a theoretical point-spread function computed from the recorded acquisition parameters. For the depth-coded images in **Extended Data Fig. 12**, each deconvolved stack was converted into a maximum-intensity projection (MIP). In parallel, the axial index of the brightest voxel at each lateral position (the arg-max plane) was recorded to encode depth. To restrict the color scale to the signal-bearing portion of the stack (rather than the arbitrary acquisition bounds), a per-slice signal profile was computed as the summed voxel intensity of each plane; the uniform background baseline (the minimum across planes) was subtracted, and the cumulative signal distribution along z was formed. The depth color scale was then anchored between the planes corresponding to the 20th and 80th percentiles of cumulative signal (planes outside this range were clamped to the endpoints). The MIP intensity was normalized independently for each image to give comparable appearance and maximal detail. Edge detail was enhanced with an unsharp mask (radius 2 px, amount 1.6; scikit-image v0.24) applied to the intensity channel.

### HEK293 cell transfection and GFP quantification

HEK293 cells (1x10^6^ cells) were transiently transfected with the indicated plasmid combinations using Lipofectamine 2000 Transfection Reagent according to the manufacturer’s protocol. Four technical replicates from each transfection were seeded into 96-well plates.

For all experiments, equal amounts of plasmids encoding *Cp*OR9 (1 μg) and *Ab*Orco (1 μg) were maintained. When necessary, a plasmid encoding non-fluorescent Halo-tagged *Ab*Orco was included to maintain constant total amounts of *Ab*Orco DNA across conditions. Halo-*Ab*Orco alone served as the negative control, whereas full-length sfGFP-CpOR9 coexpressed with Halo-*Ab*Orco served as the positive control. To determine whether GFP complementation occurred specifically within OR-Orco complexes, *Bm*Gr9 was substituted for *Cp*OR9 as an additional negative control.

Forty-eight hours after transfection, GFP fluorescence (488-nm excitation, 525-nm emission) was measured using a Synergy Neo2 microplate reader (BioTek) at the Rockefeller University Drug Discovery Resource Center.

### Fluorescence-detection size exclusion chromatography (FSEC)

HEK293S GnTI^-^cells (ATCC CRL-3022) were cultured in DMEM supplemented with 10% fetal bovine serum and transiently transfected with pEGBM plasmids encoding StrepII-sfGFP-PPX-*Cp*OR9 and/or mVenus-PPX-*Ab*Orco using FuGENE HD transfection reagent. Forty-eight hours after transfection, cells were washed with PBS, harvested, pelleted by centrifugation, flash-frozen in liquid nitrogen, and stored at -20°C until use.

Frozen pellets were resuspended in buffer containing 375 mM NaCl and 20 mM HEPES (pH 7.5) supplemented with 1 mM PMSF and EDTA-free Complete Protease Inhibitor Cocktail (Roche). Membrane proteins were solubilized for 2 h at 4°C in 0.75% (w/v) n-dodecyl-β-D-maltoside (DDM; Anatrace) supplemented with 0.15% (w/v) cholesterol hemisuccinate (CHS; Sigma-Aldrich). Insoluble material was removed by centrifugation (15 kxg, 15 min, 4°C), followed by clarification through Spin-X centrifuge filters (15 kxg, 10 min, 4°C).

Clarified lysates were loaded into a Shimadzu SIL-20AC HT autosampler and injected onto a Superose 6 Increase size-exclusion column (Cytiva) preequilibrated in FSEC buffer (150 mM NaCl, 20 mM HEPES pH 7.5, 0.025% DDM, 0.005% CHS) and connected to an ÄKTA Pure chromatography system (Cytiva). Fluorescence was monitored using a Shimadzu dual fluorescence detector configured for GFP (450-nm excitation, 505-nm emission) and mVenus (520-nm excitation, 545-nm emission).

### Multiple sequence alignments and conservation analysis

Multiple sequence alignments (MSAs) were produced from a curated list of ORs and Orcos^15^ using MAFFT v7.526 (FFT-NS-2, BLOSUM62) with the individual reference columns for CpOR9 and AbOrco used to identify the WHY-motif in ORs and Orcos respectively. The values from this analysis appear in **Extended Data Fig. 6a** and **11c**. The ORs are derived from four species: 221 Nasonia vitripennis, 72 Anopheles gambiae, 61 Drosophila melanogaster, 7 Pediculus humanus corporis. The Orco list represents 115 genera, with each Orco from a unique species, except *Thermobia domestica* (a basal species with three putative Orco sequences).

### Structural illustrations and plotting

Structural figures were rendered in UCSF ChimeraX 1.11 from the coordinates and maps described above; structural superimposition was calculated with *matchmaker* (ChimeraX), interactions were calculated with *hbonds* (ChimeraX) for hydrogen bonds, and PLIP^82^ for ᴨ-interactions; quantitative panels were plotted in Python 3 (matplotlib) from the source data; panels were assembled in Adobe Illustrator CC. Rendering, plotting and layout scripts were drafted with assistance from Claude Code (Claude Opus 4.8, Anthropic) and were reviewed, executed and verified by the authors. All displayed data derive from the experimental maps, models and measurements reported here; schematics were hand-drawn by the authors, no image content was generated using generative AI.

---

**Extended Data Table 1.** Cryo-EM data collection, refinement and validation statistics *Ab*Orco–*Cp*OR9 with VUAA4 and indole.

|  | 4:0 *AbOrco*<br>(EMDB-xxxx)<br>(PDB xxxx) | 3:1 *AbOrco*–*CpOR9*<br>(EMDB-xxxx)<br>(PDB xxxx) | 2:2 *AbOrco*–*CpOR9*<br>(EMDB-xxxx)<br>(PDB xxxx) |
| --- | --- | --- | --- |
| **Data collection and processing** |  |  |  |
| Magnification | 105,000× | 105,000× | 105,000× |
| Voltage (kV) | 300 | 300 | 300 |
| Electron exposure (e<sup>−</sup>/Å<sup>2</sup>) | 55 | 55 | 55 |
| Defocus range (μm) | 0.84–2.01 | 0.84–2.01 | 0.84–2.01 |
| Pixel size (Å) | 0.847 | 0.847 | 0.847 |
| Symmetry imposed | C4 | C1 | C2 |
| Micrographs (no.) | 20,789 | 20,789 | 20,789 |
| Initial particle images (no.) | 3,298,267 | 3,298,267 | 3,298,267 |
| Final particle images (no.) | 325,790 | 682,644 | 58,308 |
| Map resolution (Å) | 2.15 (1.95) | 2.25 (2.01) | 2.82 (2.56) |
| FSC threshold | 0.143 | 0.143 | 0.143 |
| **Refinement** |  |  |  |
| Map sharpening B factor (Å<sup>2</sup>) | −60.7 | −60.8 | −68.3 |
| **Model composition** |  |  |  |
| Non-hydrogen atoms | 13,652 | 13,291 | 12,990 |
| Protein residues | 1,612 | 1,586 | 1,560 |
| Ligands | CLR 4, PTY 12, VU4 4 | CLR 3, IND 1, PTY 9, VU4 3 | CLR 2, IND 2, PTY 6, VU4 2 |
| **B factors (Å<sup>2</sup>)** |  |  |  |
| Protein | 44.2 | 55.0 | 54.0 |
| Ligand | 56.2 | 68.2 | 62.3 |
| **R.m.s. deviations** |  |  |  |
| Bond lengths (Å) | 0.0080 | 0.0141 | 0.0188 |
| Bond angles (°) | 1.20 | 1.85 | 1.08 |
| **Validation** |  |  |  |
| MolProbity score | 0.83 | 0.53 | 1.07 |
| Clashscore | 0.54 | 0.07 | 2.23 |
| Poor rotamers (%) | 1.56 | 0.72 | 1.24 |
| **Ramachandran plot** |  |  |  |
| Favored (%) | 99.62 | 98.16 | 98.39 |
| Allowed (%) | 0.38 | 1.71 | 1.61 |
| Disallowed (%) | 0.00 | 0.13 | 0.00 |

**Extended Data Table 2.** Cryo-EM data collection, refinement and validation statistics *Ab*Orco–*Cp*OR9, ligand-free (apo)

|  | 4:0 *AbOrco*<br>(EMDB-xxxx)<br>(PDB xxxx) | 3:1 *AbOrco*–*CpOR9*<br>(EMDB-xxxx)<br>(PDB xxxx) |
| --- | --- | --- |
| **Data collection and processing** |  |  |
| Magnification | 105,000× | 105,000× |
| Voltage (kV) | 300 | 300 |
| Electron exposure (e<sup>−</sup>/Å<sup>2</sup>) | 60 | 60 |
| Defocus range (μm) | 1.18–3.14 | 1.18–3.14 |
| Pixel size (Å) | 0.847 | 0.847 |
| Symmetry imposed | C4 | C1 |
| Micrographs (no.) | 17,232 | 17,232 |
| Initial particle images (no.) | 830,921 | 830,921 |
| Final particle images (no.) | 60,845 | 70,929 |
| Map resolution (Å) | 2.85 (2.73) | 2.95 (2.79) |
| FSC threshold | 0.143 | 0.143 |
| **Refinement** |  |  |
| Map sharpening B factor (Å<sup>2</sup>) | −83.9 | −64.1 |
| **Model composition** |  |  |
| Non-hydrogen atoms | 13,560 | 13,289 |
| Protein residues | 1,612 | 1,584 |
| Ligands | CLR 4, PTY 12 | CLR 4, CPL 3, PTY 7 |
| **B factors (Å<sup>2</sup>)** |  |  |
| Protein | 59.7 | 57.6 |
| Ligand | 75.0 | 56.0 |
| **R.m.s. deviations** |  |  |
| Bond lengths (Å) | 0.0042 | 0.0146 |
| Bond angles (°) | 0.80 | 0.82 |
| **Validation** |  |  |
| MolProbity score | 1.20 | 1.32 |
| Clashscore | 3.09 | 4.11 |
| Poor rotamers (%) | 1.35 | 1.44 |
| **Ramachandran plot** |  |  |
| Favored (%) | 98.93 | 98.98 |
| Allowed (%) | 1.07 | 1.02 |
| Disallowed (%) | 0.00 | 0.00 |

**Extended Data Table 3.**
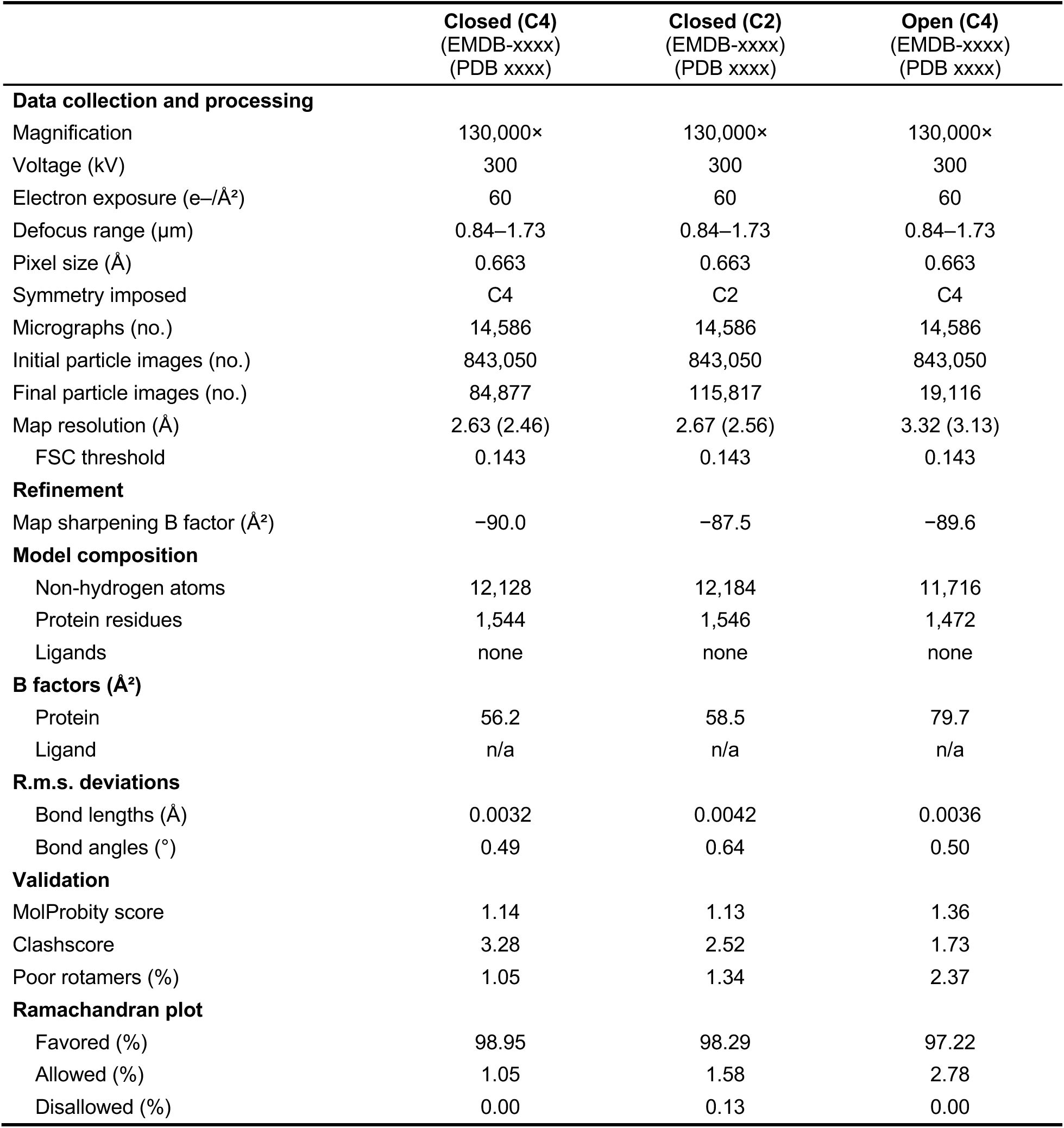
Cryo-EM data collection, refinement and validation statistics *Mh*OR5, ligand-free (apo)

|  | Closed (C4)<br>(EMDB-xxxx)<br>(PDB xxxx) | Closed (C2)<br>(EMDB-xxxx)<br>(PDB xxxx) | Open (C4)<br>(EMDB-xxxx)<br>(PDB xxxx) |
| --- | --- | --- | --- |
| **Data collection and processing** |  |  |  |
| Magnification | 130,000× | 130,000× | 130,000× |
| Voltage (kV) | 300 | 300 | 300 |
| Electron exposure (e<sup>-</sup>/Å<sup>2</sup>) | 60 | 60 | 60 |
| Defocus range (μm) | 0.84–1.73 | 0.84–1.73 | 0.84–1.73 |
| Pixel size (Å) | 0.663 | 0.663 | 0.663 |
| Symmetry imposed | C4 | C2 | C4 |
| Micrographs (no.) | 14,586 | 14,586 | 14,586 |
| Initial particle images (no.) | 843,050 | 843,050 | 843,050 |
| Final particle images (no.) | 84,877 | 115,817 | 19,116 |
| Map resolution (Å) | 2.63 (2.46) | 2.67 (2.56) | 3.32 (3.13) |
| FSC threshold | 0.143 | 0.143 | 0.143 |
| **Refinement** |  |  |  |
| Map sharpening B factor (Å<sup>2</sup>) | −90.0 | −87.5 | −89.6 |
| **Model composition** |  |  |  |
| Non-hydrogen atoms | 12,128 | 12,184 | 11,716 |
| Protein residues | 1,544 | 1,546 | 1,472 |
| Ligands | none | none | none |
| **B factors (Å<sup>2</sup>)** |  |  |  |
| Protein | 56.2 | 58.5 | 79.7 |
| Ligand | n/a | n/a | n/a |
| **R.m.s. deviations** |  |  |  |
| Bond lengths (Å) | 0.0032 | 0.0042 | 0.0036 |
| Bond angles (°) | 0.49 | 0.64 | 0.50 |
| **Validation** |  |  |  |
| MolProbity score | 1.14 | 1.13 | 1.36 |
| Clashscore | 3.28 | 2.52 | 1.73 |
| Poor rotamers (%) | 1.05 | 1.34 | 2.37 |
| **Ramachandran plot** |  |  |  |
| Favored (%) | 98.95 | 98.29 | 97.22 |
| Allowed (%) | 1.05 | 1.58 | 2.78 |
| Disallowed (%) | 0.00 | 0.13 | 0.00 |

**Extended Data Fig. 1.**
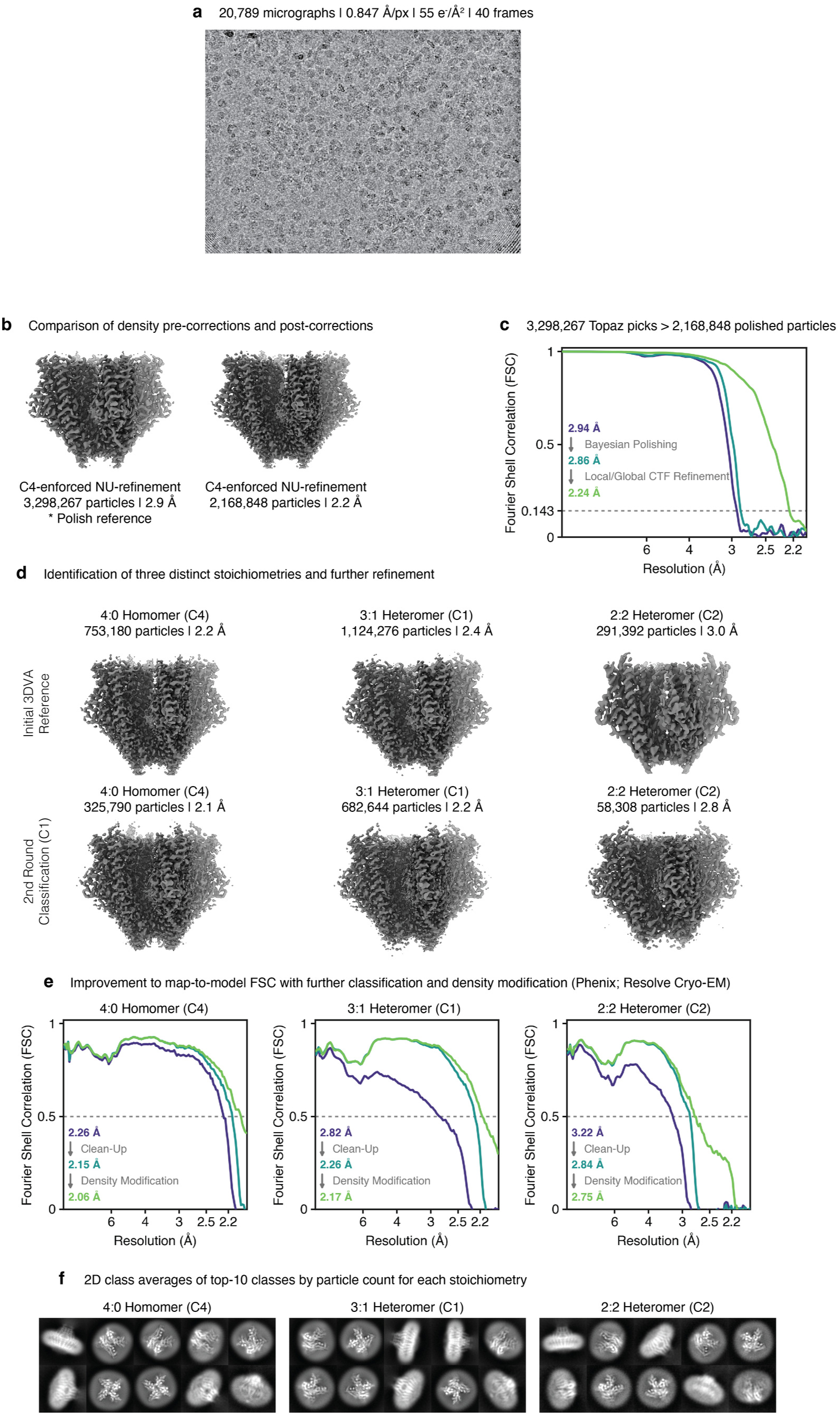
Cryo-EM analysis and model building workflow for the indole and VUAA4 bound CpOR9:AbOrco heteromer structures. **a**, Representative micrograph and acquisition specifications. **b**, The consensus particle stack and resultant refinement (left) used for Bayesian polishing, and local/global CTF refinements to yield the final refinement stack (right). **c**, Improvements to initial map through Bayesian Polishing followed by local/global CTF refinement (FSC=0.143). **d**, Initial identification of three distinct states with 3DVA (top row) and resultant final refinements after classification (bottom row). **e**, Improvements to map-to-model fit (FSC=0.5) through classification and Phenix density modification (Resolve Cryo-EM). **f**, 2D class averages from the final particle stacks for the top-10 classes by occupancy out of 100.

**Extended Data Fig. 2.**
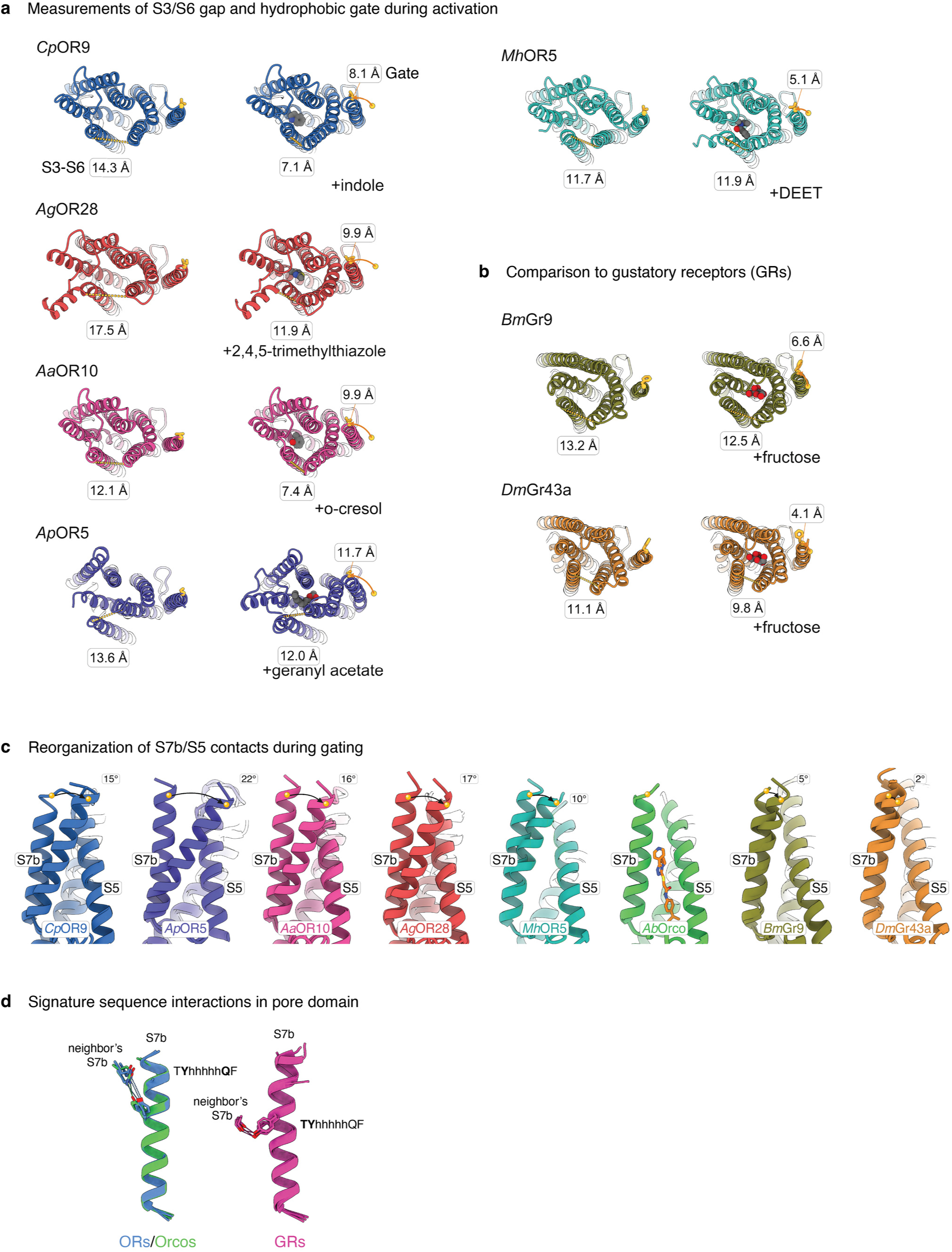
Gating rearrangements across ORs and GRs. **a**, Comparison of monomeric ligand-free and ligand-bound conformations highlights conserved movements associated with gating across heteromeric and homomeric ORs, with narrowing of S3:S6 gap and tilting of S7b. Cα distances between S3 and S6 and movement of S7b gate shown. **b**, Monomeric movements during gating across homomeric GRs. **c**, Conserved repacking between S7b and S5 during gating for ORs and GRs, with angular sweep of S7b measured. **d**, Conservation of hydrogen bond interactions between signature sequence residues (TYhhhhhQF) for ORs (left; blue), Orcos (left, green), and GRs (right, magenta) in apo/closed pore. ORs/Orcos form interactions between Y and Q, GRs form interactions between Y and T.

**Extended Data Fig. 3.**
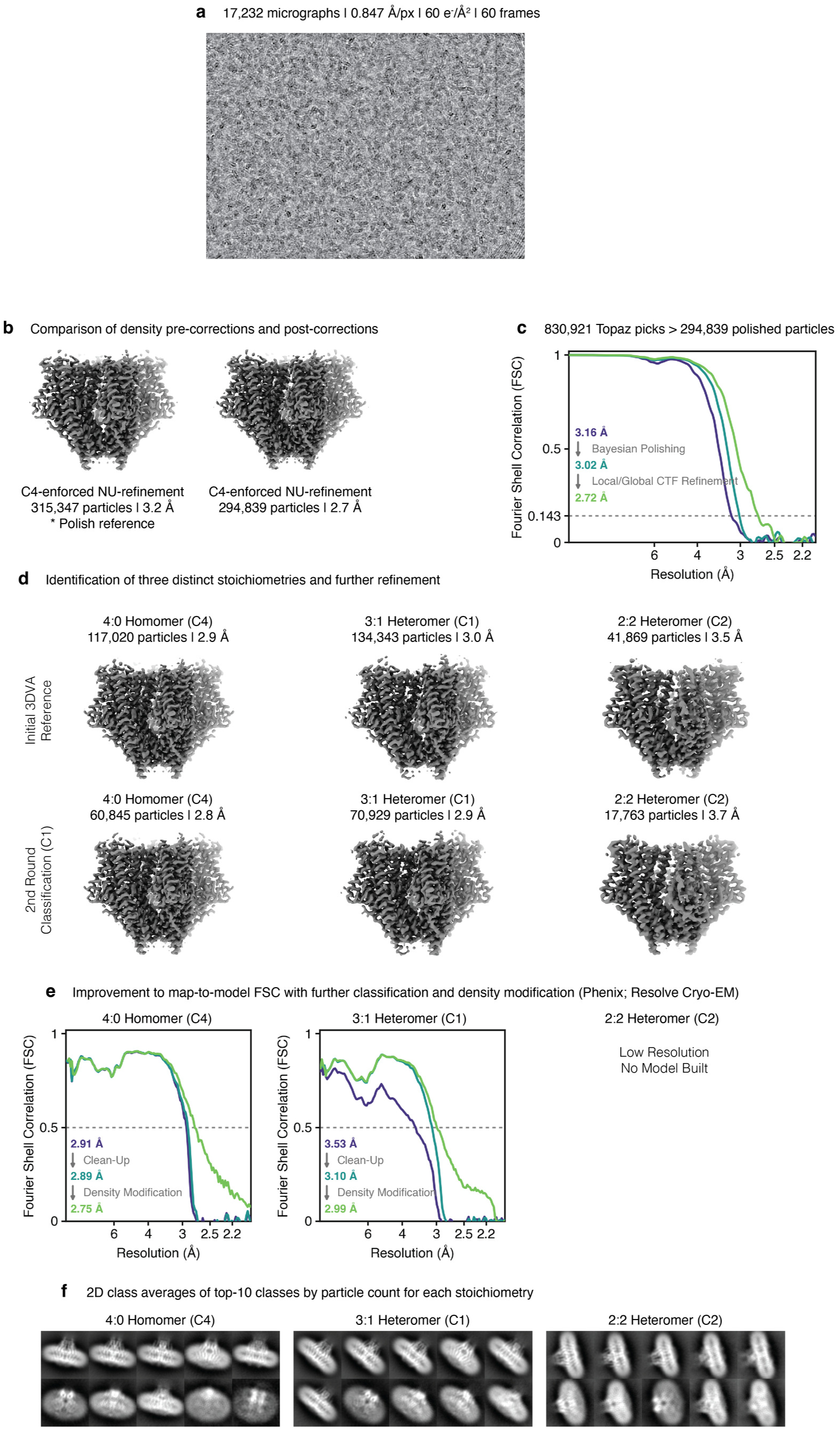
Cryo-EM analysis and model building workflow for the ligand-free CpOR9:AbOrco heteromer structures. **a**, Representative micrograph and acquisition specifications. **b**, The consensus particle stack and resultant refinement (left) used for Bayesian polishing, and local/global CTF refinements to yield the final refinement stack (right). **c**, Improvements to initial map through Bayesian Polishing followed by local/global CTF refinement (FSC=0.143). **d**, Initial identification of three distinct states with 3DVA (top row) and resultant final refinements after classification (bottom row). **e**, Improvements to map-to-model fit (FSC=0.5) through classification and Phenix density modification (Resolve Cryo-EM). **f**, 2D class averages from the final particle stacks for the top-10 classes by occupancy out of 100.

**Extended Data Fig. 4.**
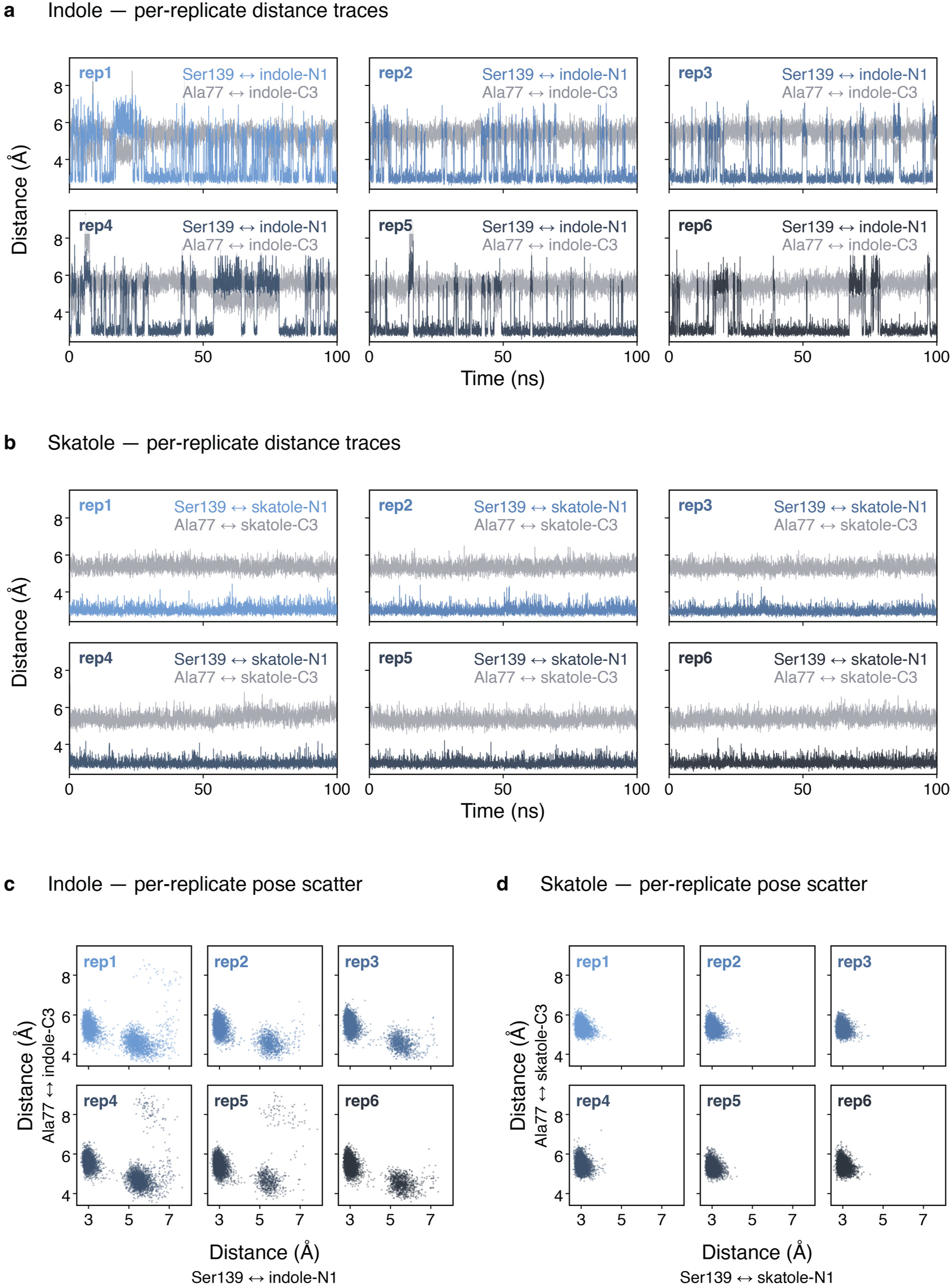
Molecular dynamics simulations of CpOR9 capture bistability of indole pose and monostability of skatole pose in the binding pocket. **a**, Indole per-replicate distance traces: distance between Ser139 and indole nitrogen-1 in each replicate’s color and Ala77 and indole carbon-3 in graphite, versus time. **b**, Skatole traces showing the same measurements as in (**a**). **c**, Indole per-replicate pose scatter for the two distance metrics. **d**, Skatole per-replicate pose scatter.

**Extended Data Fig. 5.**
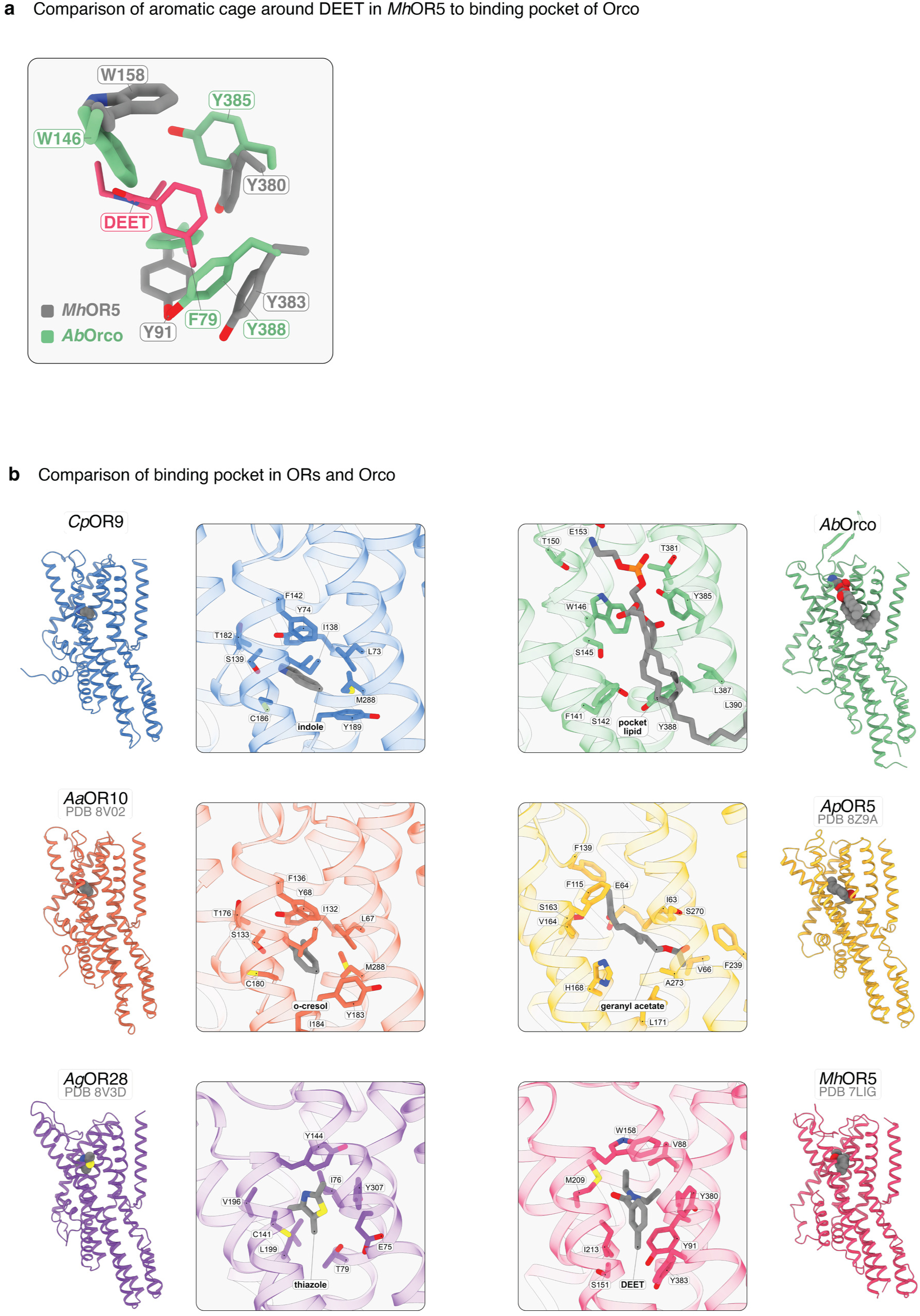
Comparison of lipid-bound pocket in Orco to odorant-bound pockets in ORs. **a**, Comparison of the conserved aromatic cage between the ancestral receptor *Mh*OR5 (gray) bound to DEET (magenta) and *Ab*Orco (green) highlighting how Trp146 in *Ab*Orco adopts a different rotamer to maintain continuity of the pocket with the extracellular solvent whereas in *Mh*OR5 Trp158 shields DEET. **b**, Binding pockets occupied by odorants for *Cp*OR9, *Aa*OR10, *Ag*OR28, *Ap*OR5, and *Mh*OR5 compared to lipid-bound state in *Ab*Orco.

**Extended Data Fig. 6.**
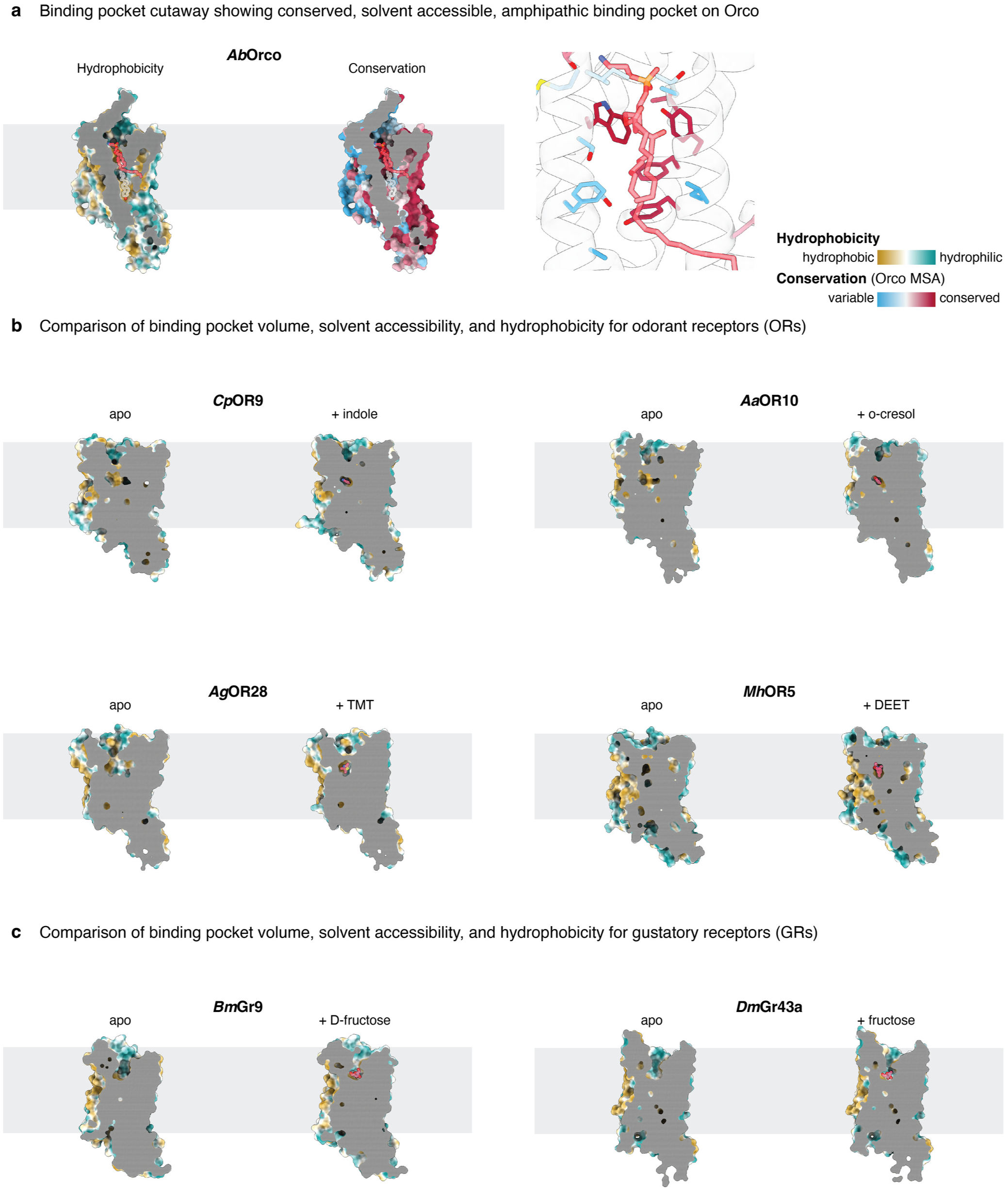
Location, conservation and chemical nature of binding pockets in Orco, ORs, and GRs. **a**, Hydrophobicity (left) and conservation (right) of Orco’s binding pocket occupied by a membrane phospholipid. **b**, Hydrophobicity and solvent-accessibility of OR binding pockets in ligand-free (left) and odorant-bound (right) states for *Cp*OR9, *Aa*OR10, *Ag*OR28, and *Mh*OR5. **c**, Hydrophobicity of GR binding pockets in ligand-free (left) and tastant-bound (right) states for *Bm*Gr9 and *Dm*Gr43a.

**Extended Data Fig. 7.**
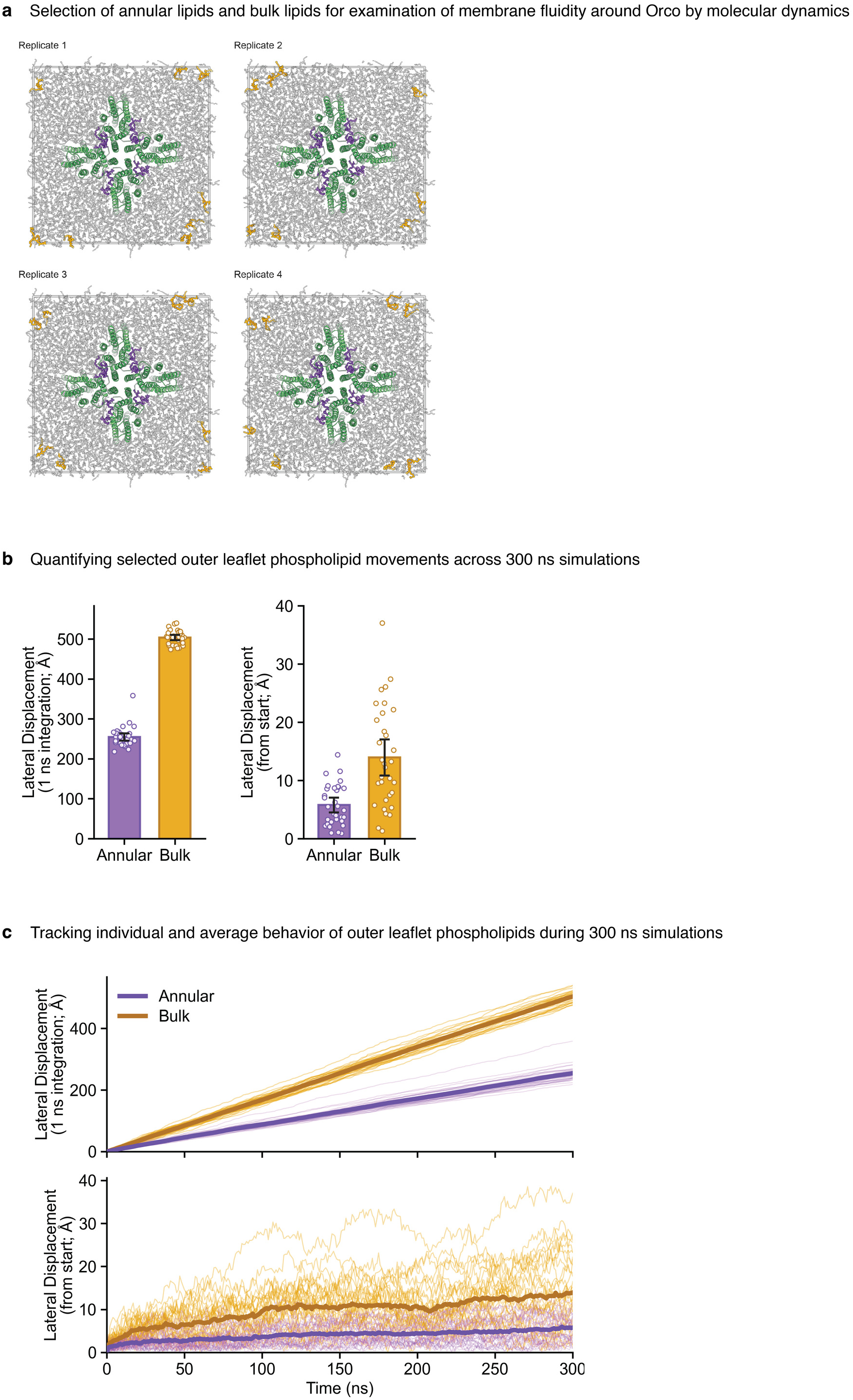
Molecular dynamics simulations quantifying displacement of annular lipids corralled between subunits. **a**, Top view of four simulated replicates of the *Ab*Orco homomer with selected outer-leaflet annular lipids (purple) and bulk lipids (amber) shown. **b**, Integrated path (left) and total displacement from start (right) of annular lipids (purple) versus bulk lipids (amber). Each point represents one lipid, error bars represent 95% confidence interval of the mean. **c**, Cumulative integrated path (top) or displacement from start (bottom) for annular lipids (purple) versus bulk lipids (amber), plotted for each individual lipid measured in a thin line, and the average for the entire cohort in the thick line.

**Extended Data Fig. 8.**
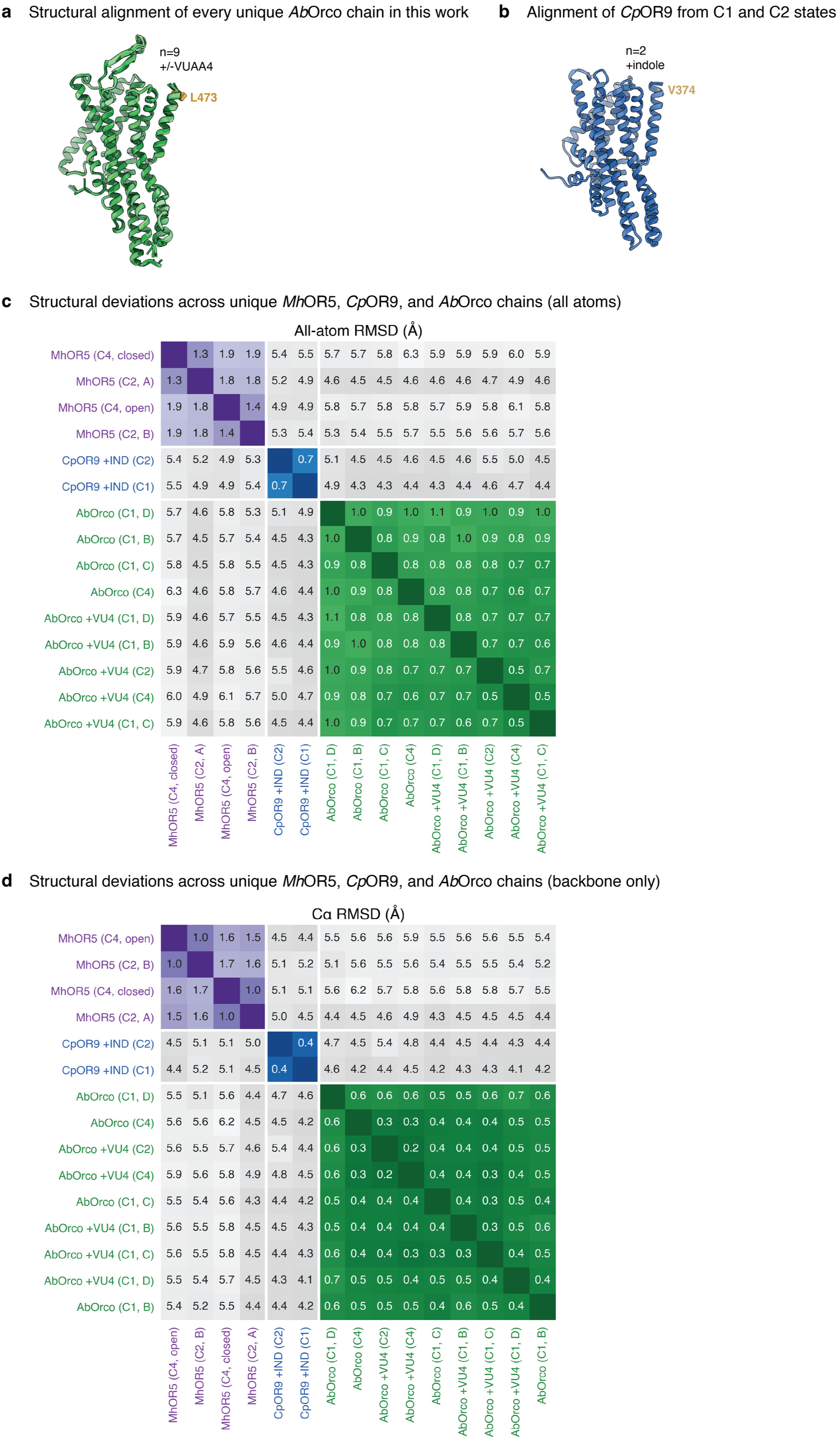
Superimposition of AbOrco, CpOR9, and MhOR5 structures across datasets. **a**, Superimposition of all nine unique *Ab*Orco protomers from apo and VUAA4-bound states across three stoichiometries. The hydrophobic gate (Leu473) shown as amber stick for a fiducial marker. **b**, Superimposition of two unique *Cp*OR9 protomers bound to indole as blue cartoons with the hydrophobic gate (Val374) shown as amber sticks. **c**, All-atom pairwise RMSD matrix over 15 unique protomers: 9x *Ab*Orco (green), 2x CpOR9 (blue), *4x* MhOR5 (purple), aligned by ChimeraX matchmaker algorithm, and hierarchically clustered. **d**, The corresponding Cα-only matrix.

**Extended Data Fig. 9.**
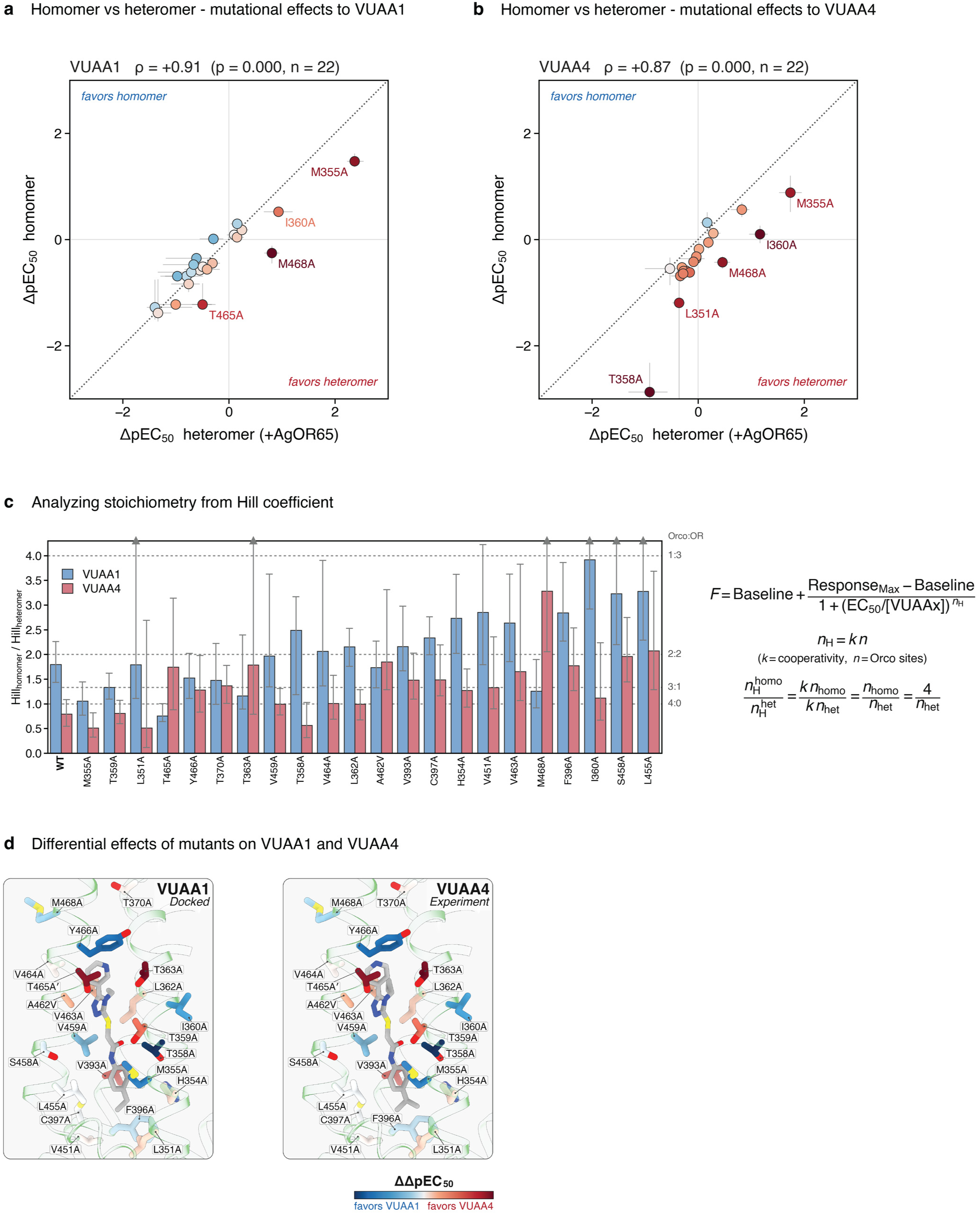
Further analysis of VUAA1 versus VUAA4, comparing across homomeric and heteromeric receptors. **a**, Differential effects of mutations in a homomeric (*Ab*Orco) versus heteromeric (*Ab*Orco+*Ag*OR65) context for VUAA1 revealed by deviations from the isoaffinity line. **b**, Differential effects of mutations in a homomeric (*Ab*Orco) versus heteromeric (*Ab*Orco+*Ag*OR65) context for VUAA1 revealed by deviations from the isoaffinity line. **c**, Investigating the relationship between Hill coefficients and subunit composition. The Hill model used throughout the work is shown on the right, where we fit both the minimum and maximum plateau and Hill coefficient to the underlying dose-response data to calculate EC_50_. By deconvolving the Hill coefficient into two terms, *n* for the number of Orco binding sites per tetramer, and *k* for the cooperativity across subunits, assuming cooperativity does not vary, we can approximate the number of Orco subunits from the Hill coefficient by dividing the homomeric and heteromeric responses. We observe variation in the modeled number of Orco binding sites across mutants for VUAA1 and VUAA4, suggesting that the Hill coefficient is not a reliable indicator of subunit composition for heteromeric Orco-OR systems. **d**, Differential effects of VUAA1 and VUAA4 from Fig. 2l, m, mapped onto the structures of docked VUAA1 (left) and experimental VUAA4 (right), highlighting how differential mutants cluster around the sites of molecular variance between the two agonists.

**Extended Data Fig. 10.**
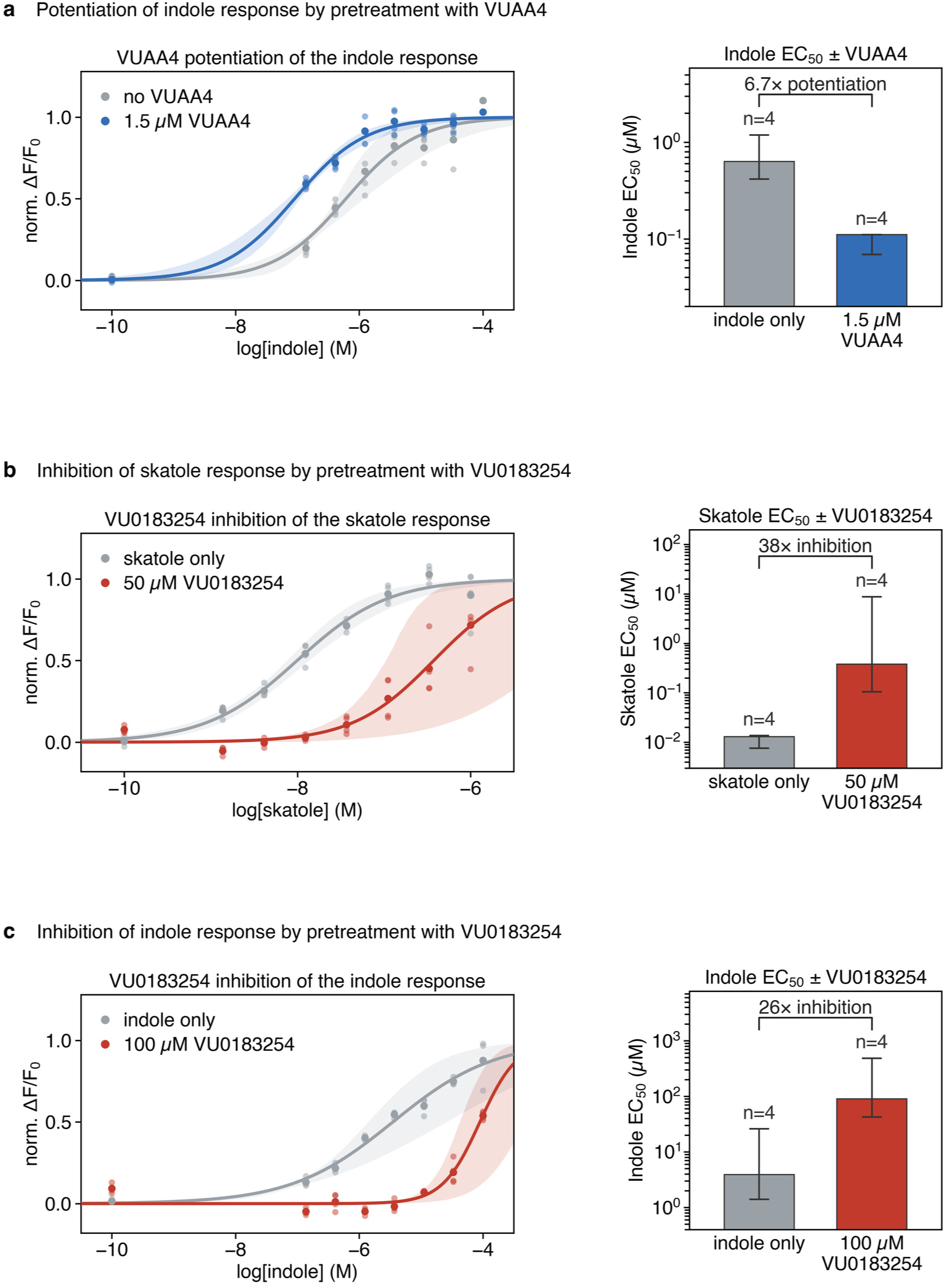
Synergistic effects of Orco agonists and antagonists on odor-mediated signaling. **a**, Pretreatment of *Ab*Orco-*Cp*OR9 heteromers with Orco-agonist (1.5 µM VUAA4) increases the apparent potency of subsequent indole response by 6.7-fold. **b**, Pretreatment of *Ab*Orco-*Cp*OR9 heteromers with Orco-antagonist (50 µM VU0183254) decreases the apparent potency of subsequent skatole response by 38-fold. **c**, Pretreatment of *Ab*Orco-*Cp*OR9 heteromers with Orco-antagonist (100 µM VU0183254) decreases the apparent potency of subsequent indole response by 26-fold. For (**a**–**c**), the dose-response curves show every underlying point as a small dot, and the concentration-average as a large dot. The transparent envelope is derived from 500 bootstrap fits of the Hill model to the underlying well-level measurements. Curves are normalized between 0 and 1 by the Hill model fit. Error bars on the bar plots represent the 95% confidence interval derived from 2000 bootstrap fits of the underlying data. *n*=4 represents total number of replicates per condition.

**Extended Data Fig. 11.**
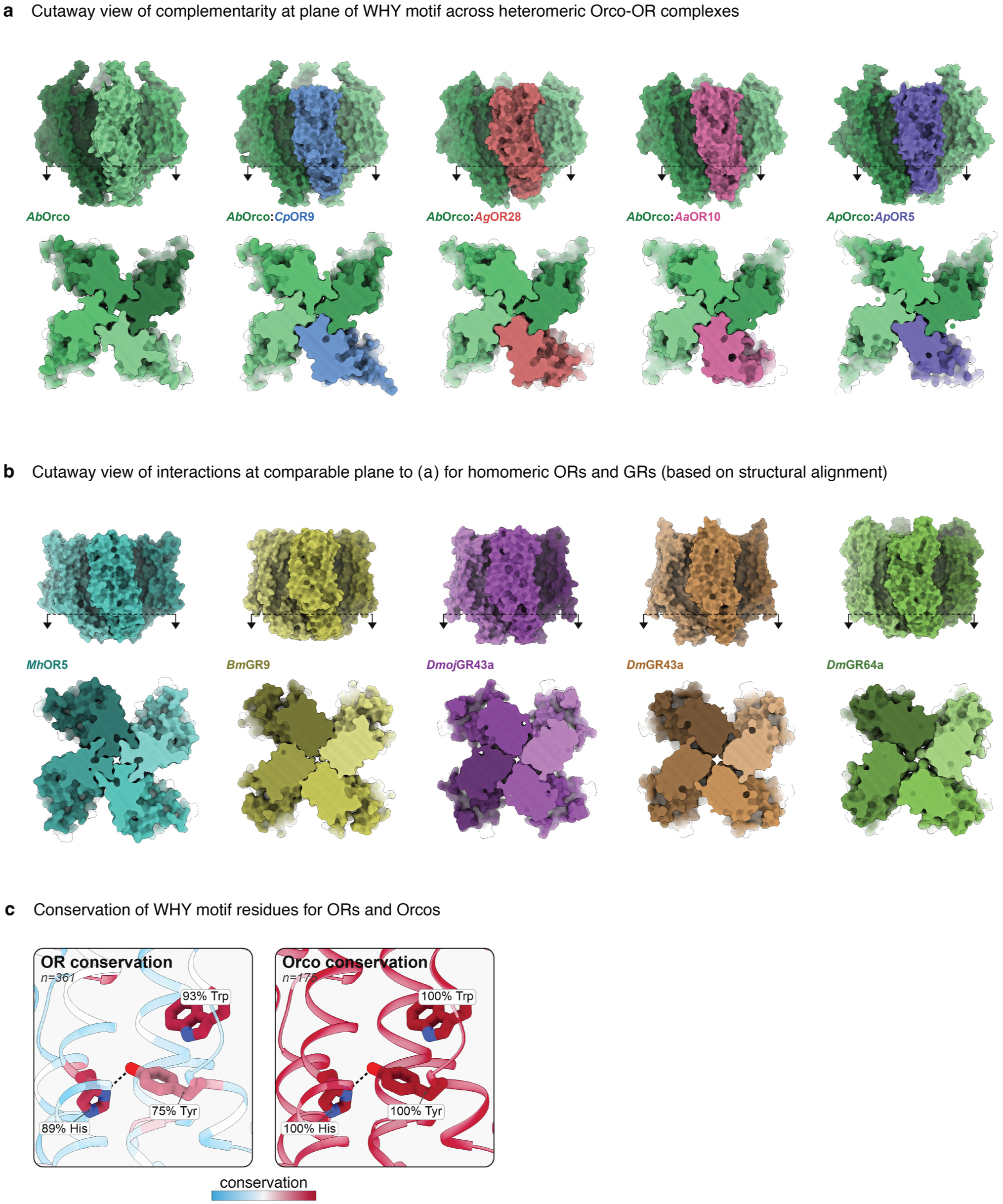
Comparison of the anchor domain assembly in Orco-OR heteromers, OR homomers, and GR homomers. **a**, Side view (top) showing the height of the WHY motif and corresponding slice through the WHY motif (bottom) for divergent 3:1 Orco-OR heteromers. **b**, Side view (top) showing the height of the WHY motif and corresponding slice through the WHY motif (bottom) for the basal homomeric MhOR5 (left, teal) and various homomeric GRs. **c**, Conservation of WHY motif in ORs and Orcos highlights the distinctive local conservation of WHY motif residues in ORs, with percent identity labeled for each. See Methods for details on multiple sequence alignments used to produce the conservation values.

**Extended Data Fig. 12.**
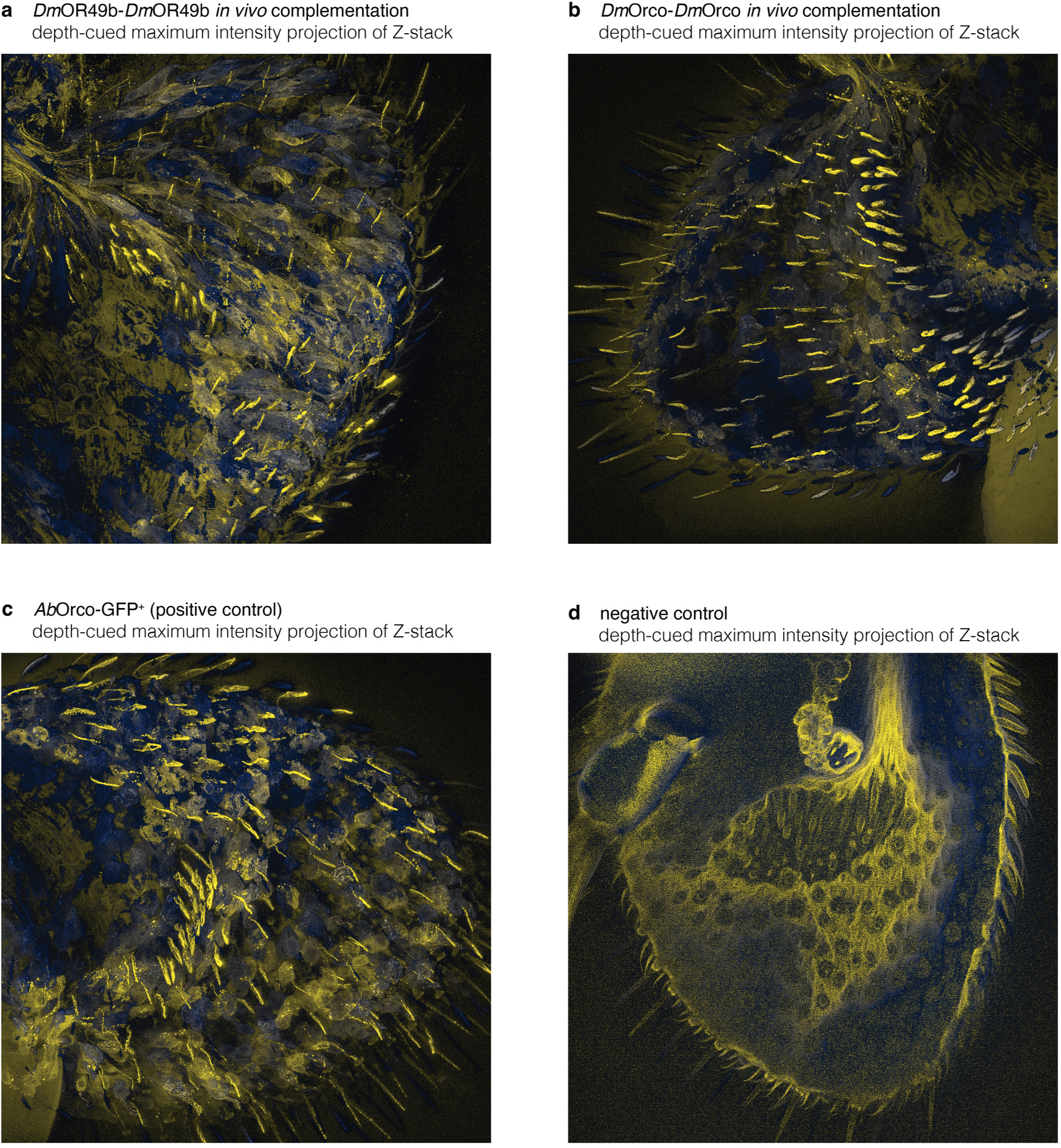
*In vivo* split-GFP-complementation in *Drosophila* antennae. **a**, GFP_1-10_:*Dm*OR49b and GFP_11_:*Dm*OR49b co-expressed in *D. melanogaster* under the *Orco* promoter, imaged via confocal Z-stack, maximum-intensity projected and pseudo-colored by depth on a scale from yellow (near) to dark blue (far) to highlight sensillar signal amongst cell soma. **b**, GFP_1-10_:*Dm*Orco and GFP_11_:*Dm*Orco co-expressed in *D. melanogaster*, imaged via confocal Z-stack, maximum-intensity projected and pseudo-colored by depth on a scale from yellow (dorsal) to dark blue (ventral) to highlight sensillar versus somatic signal. **c**, Expression of sfGFP:*AbOrco* under the *Orco* promoter in *D. melanogaster* as positive control. **d**, Negative control (wild-type animal, no exogenous fluorescent marker) imaged and presented under identical treatment for comparison.

**Extended Data Fig. 13.**
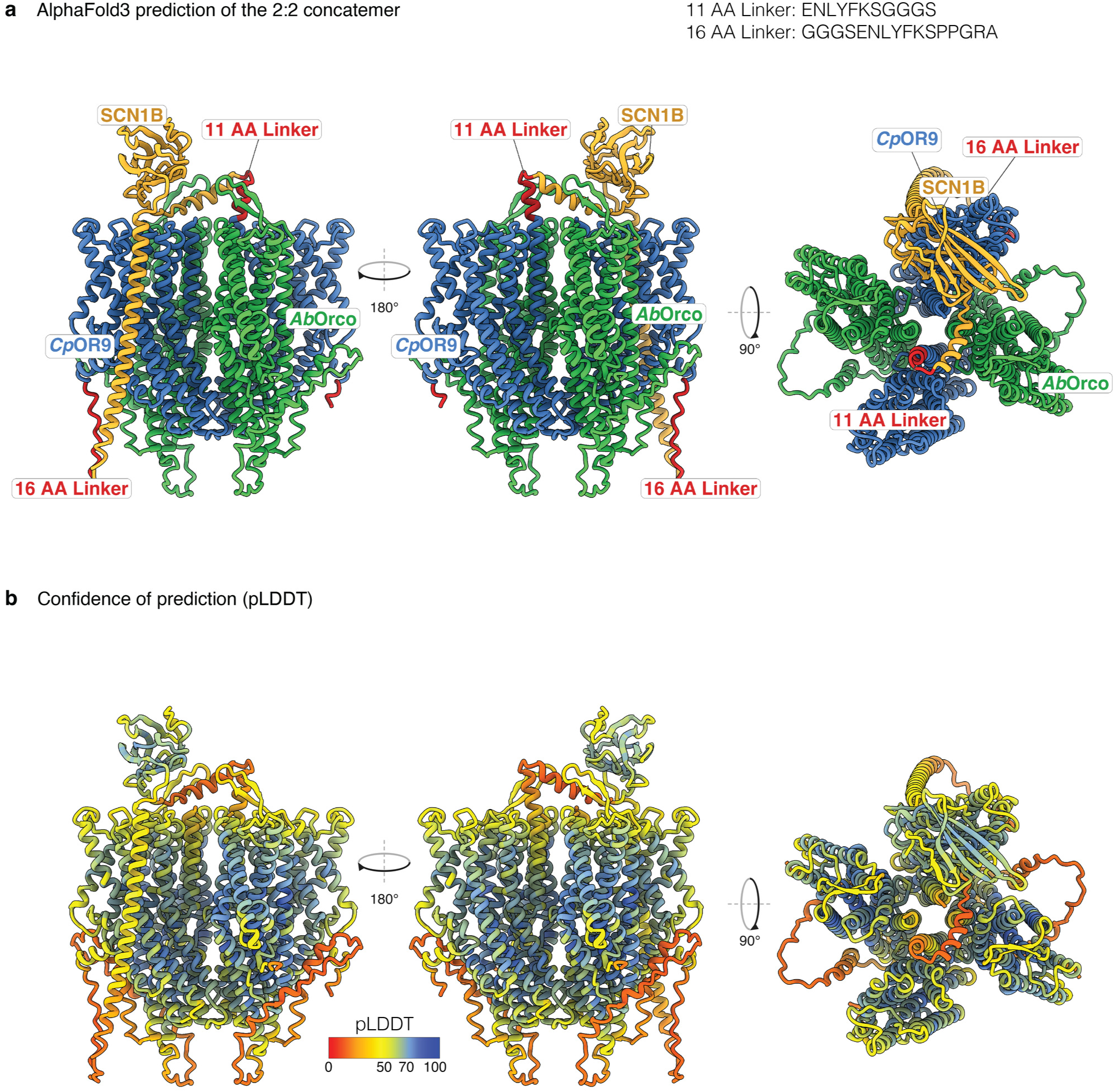
AlphaFold3 prediction of the concatenated receptor assembly mediated by SCN1B. **a**, AlphaFold3 prediction of the *Cp*OR9-SCN1B-*Cp*OR9 concatenated receptor in complex with two *Ab*Orco monomers. *Cp*OR9 sequences colored blue, *Ab*Orco sequences colored green, SCN1B sequences colored amber, and synthetic linkers colored red. **b**, Same complex as (**a**) colored by AlphaFold3 confident metric pLDDT, where pLDDT<70 has uncertain Cɑ placement.

**Extended Data Fig. 14.**
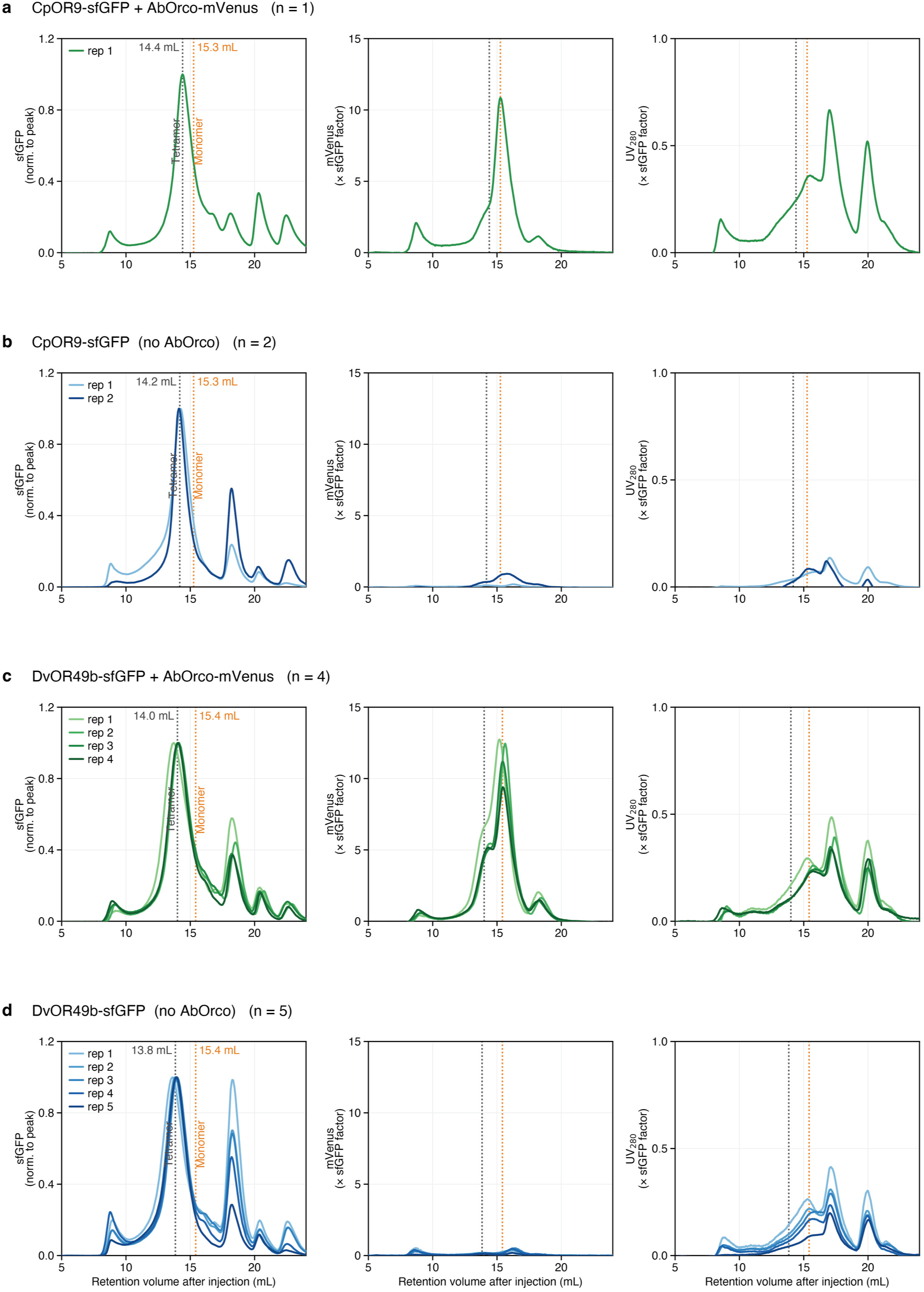
OR homotetramers form in HEK cells as indicated by fluorescence size exclusion chromatography. **a**, Fluorescence size exclusion chromatography (FSEC) of HEK cell lysates expressing sfGFP:*Cp*OR9 tagged with mVenus*:Ab*Orco run on Superose 6 Increase 10/300 GL column. For all panels, the left column shows sfGFP fluorescence (450 nm excitation/505 nm emission) to detect the OR, the middle column shows mVenus fluorescence (520 nm excitation/545 nm emission) to detect Orco, and the right column shows absorbance at 280 nm to detect total protein. **b**, FSEC of sfGFP:*Cp*OR9 expressed in the absence of Orco (n = 2 replicates), showing an OR-containing peak at the expected tetrameric elution volume despite the absence of an Orco signal. **c**, FSEC of sfGFP:*Dv*OR49b co-expressed with mVenus*:Ab*Orco (n = 4). **d**, FSEC of sfGFP:*Dv*OR49b expressed in the absence of Orco (n = 5), again showing an OR-containing peak at the expected tetrameric elution volume in the absence of Orco. For each replicate, each channel was normalized to the same factor used to align sfGFP. Vertical dotted lines indicate the approximate expected elution volumes of tetrameric and monomeric receptor species.

**Extended Data Fig. 15.**
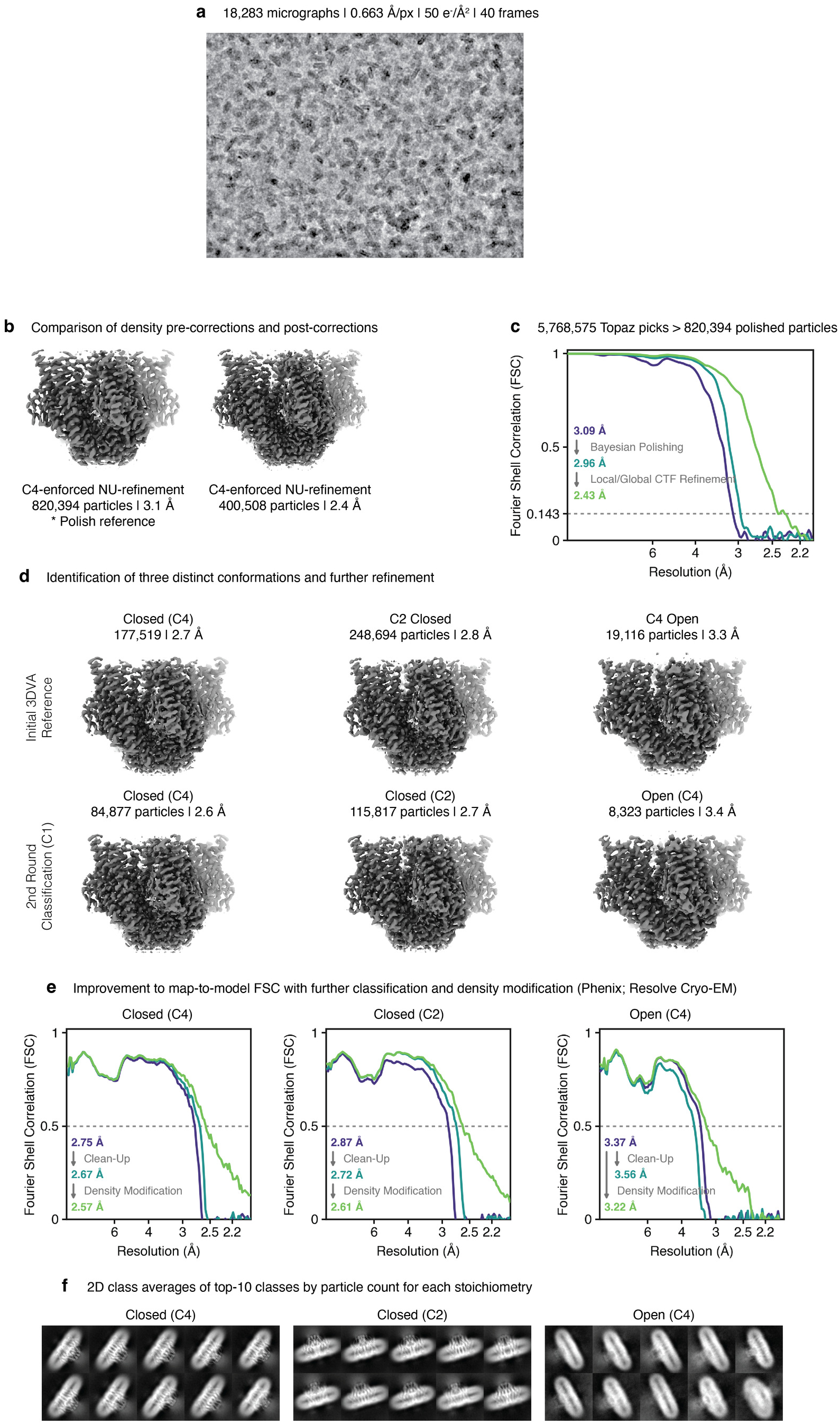
Cryo-EM analysis and model building workflow for the ligand-free MhOR5 structures. **a**, Representative micrograph and acquisition specifications. **b**, The consensus particle stack and resultant refinement (left) used for Bayesian polishing, and local/global CTF refinements to yield the final refinement stack (right). **c**, Improvements to initial map through Bayesian Polishing followed by local/global CTF refinement (FSC=0.143). **d**, Initial identification of three distinct states with 3DVA (top row) and resultant final refinements after classification (bottom row). **e**, Improvements to map-to-model fit (FSC=0.5) through classification and Phenix density modification (Resolve Cryo-EM). **f**, 2D class averages from the final particle stacks for the top-10 classes by occupancy out of 100.

**Extended Data Fig. 16.**
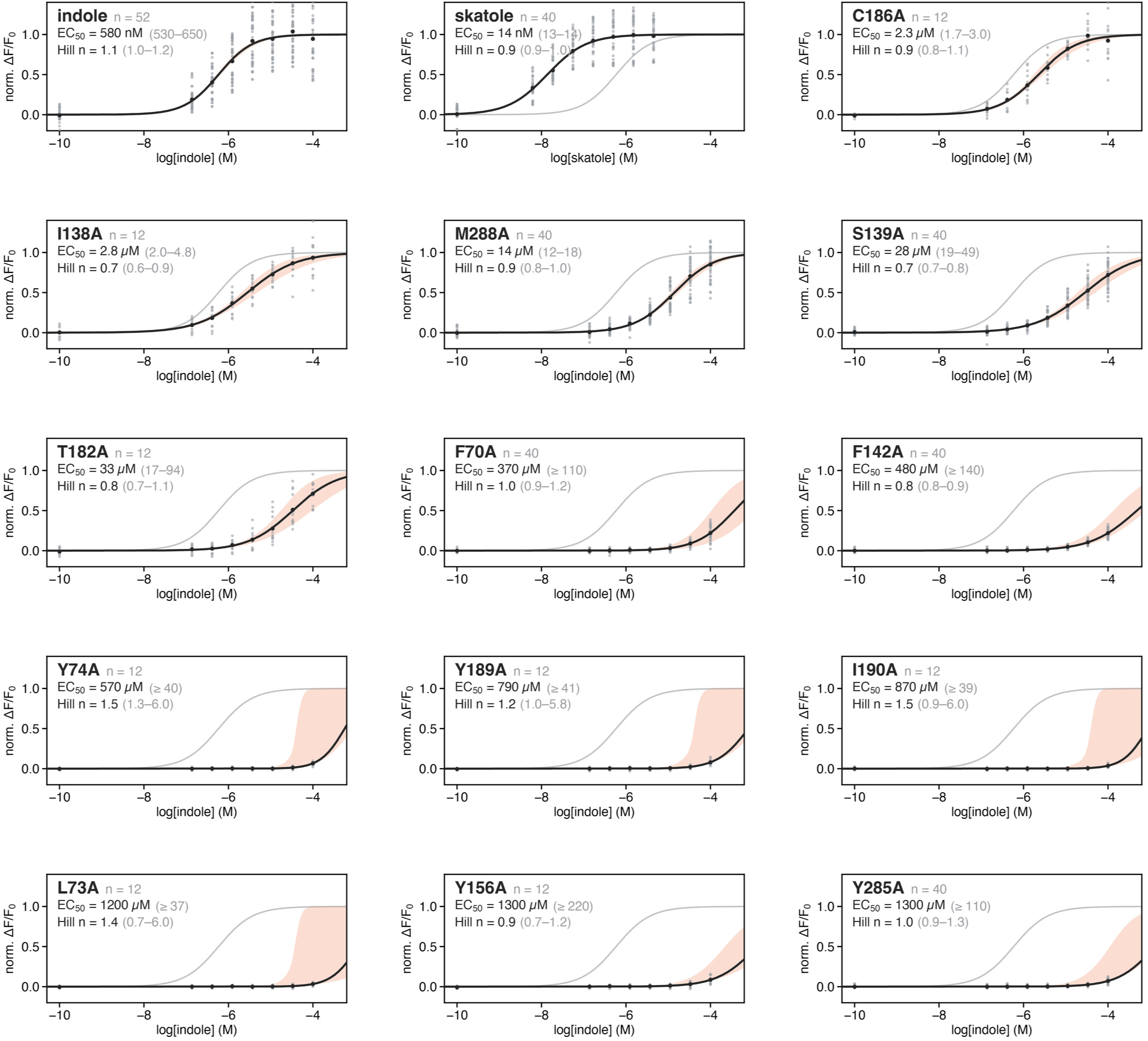
Normalized dose-response data for CpOR9 (+AbOrco) WT and mutant responses to indole and skatole for calculating EC50. The WT or mutants are designated on the top left, with each raw fluorescence ΔF/F_o_ shown as points. A four-parameter Hill model (see Extended Data Fig. 9c for exact equation) was fit to the underlying data for each, and shown as a black line. 95% confidence interval of bootstrap resampling (B=500) of the underlying data was mapped as an orange envelope around the mean curve fit, highlighting points of uncertainty. The curves were scaled between 0 and 1 (baseline and maximum response), such that 0.5 represents the EC_50_. The top left plot is the reference plot, and is recapitulated as a gray line in subsequent plots for comparison. *n* represents the number of replicate dose-series, EC_50_ and Hill coefficient (Hill *n*) are provided as mean with error bounds calculated from bootstrapping (95% CI).

**Extended Data Fig. 17.**
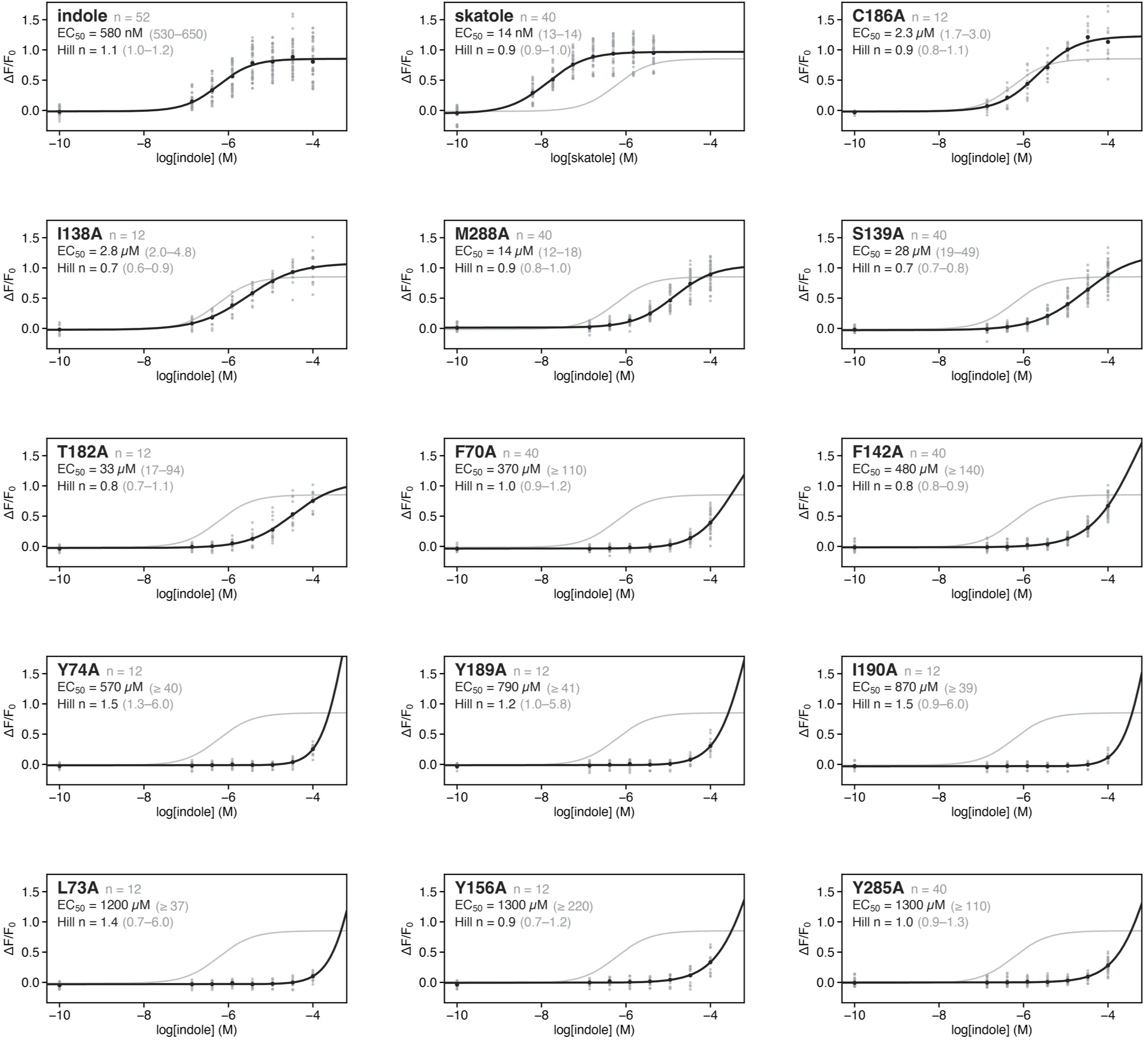
Raw dose-response data for CpOR9 (+AbOrco) WT and mutant responses to indole and skatole for calculating EC50. The WT or mutants are designated on the top left, with each raw fluorescence ΔF/F_o_ shown as points. A four-parameter Hill model (see Extended Data Fig. 9c for exact equation) was fit to the underlying data for each, and shown as a black line. The curves were scaled between 0 and 1 (baseline and maximum response), such that 0.5 represents the EC_50_. The top left plot is the reference plot, and is recapitulated as a gray line in subsequent plots for comparison. *n* represents the number of replicate dose-series, EC_50_ and Hill coefficient (Hill *n*) are provided as mean with error bounds calculated from bootstrapping (95% CI).

**Extended Data Fig. 18.**
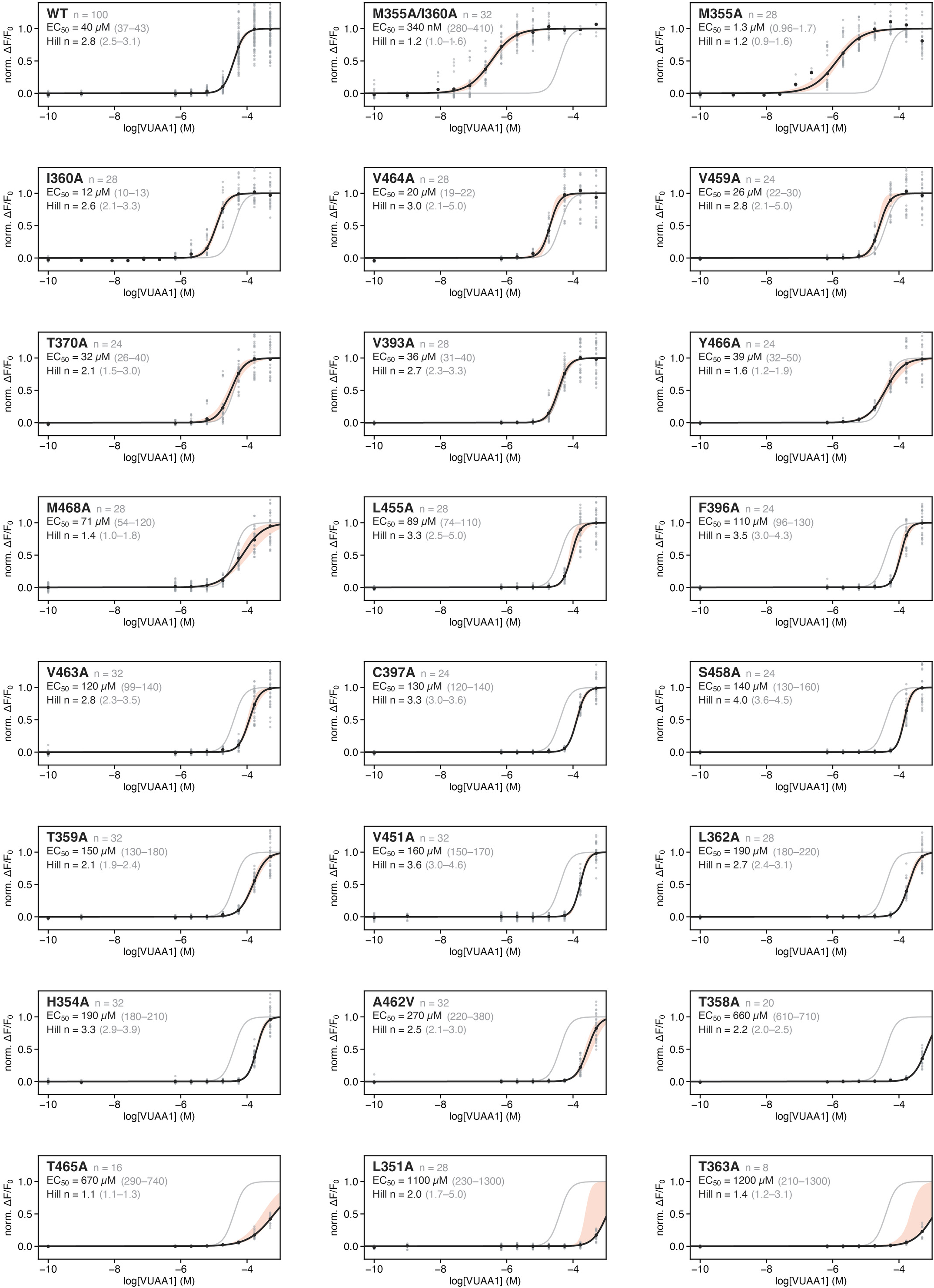
Normalized dose-response data for AbOrco WT and mutant responses to VUAA1 for calculating EC50. The WT or mutants are designated on the top left, with each raw fluorescence ΔF/F_o_ shown as points. A four-parameter Hill model (see Extended Data Fig. 9c for exact equation) was fit to the underlying data for each, and shown as a black line. 95% confidence interval of bootstrap resampling (B=500) of the underlying data was mapped as an orange envelope around the mean curve fit, highlighting points of uncertainty. The curves were scaled between 0 and 1 (baseline and maximum response), such that 0.5 represents the EC_50_. The top left plot is the reference plot, and is recapitulated as a gray line in subsequent plots for comparison. *n* represents the number of replicate dose-series, EC_50_ and Hill coefficient (Hill *n*) are provided as mean with error bounds calculated from bootstrapping (95% CI).

**Extended Data Fig. 19.**
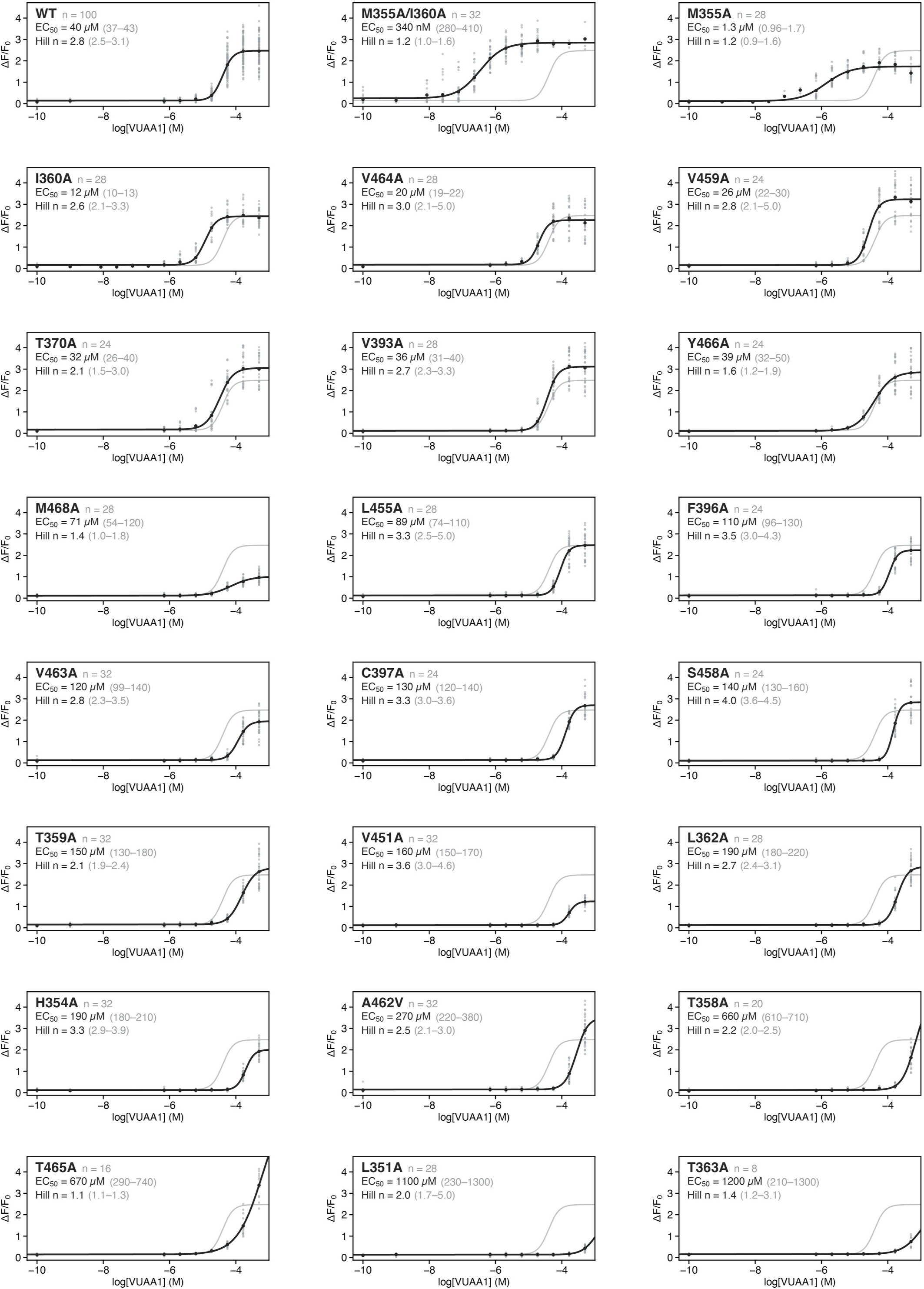
Raw dose-response data for AbOrco WT and mutant responses to VUAA1 for calculating EC50. The WT or mutants are designated on the top left, with each raw fluorescence ΔF/F_o_ shown as points. A four-parameter Hill model (see Extended Data Fig. 9c for exact equation) was fit to the underlying data for each, and shown as a black line. The curves were scaled between 0 and 1 (baseline and maximum response), such that 0.5 represents the EC_50_. The top left plot is the reference plot, and is recapitulated as a gray line in subsequent plots for comparison. *n* represents the number of replicate dose-series, EC_50_ and Hill coefficient (Hill *n*) are provided as mean with error bounds calculated from bootstrapping (95% CI).

**Extended Data Fig. 20.**
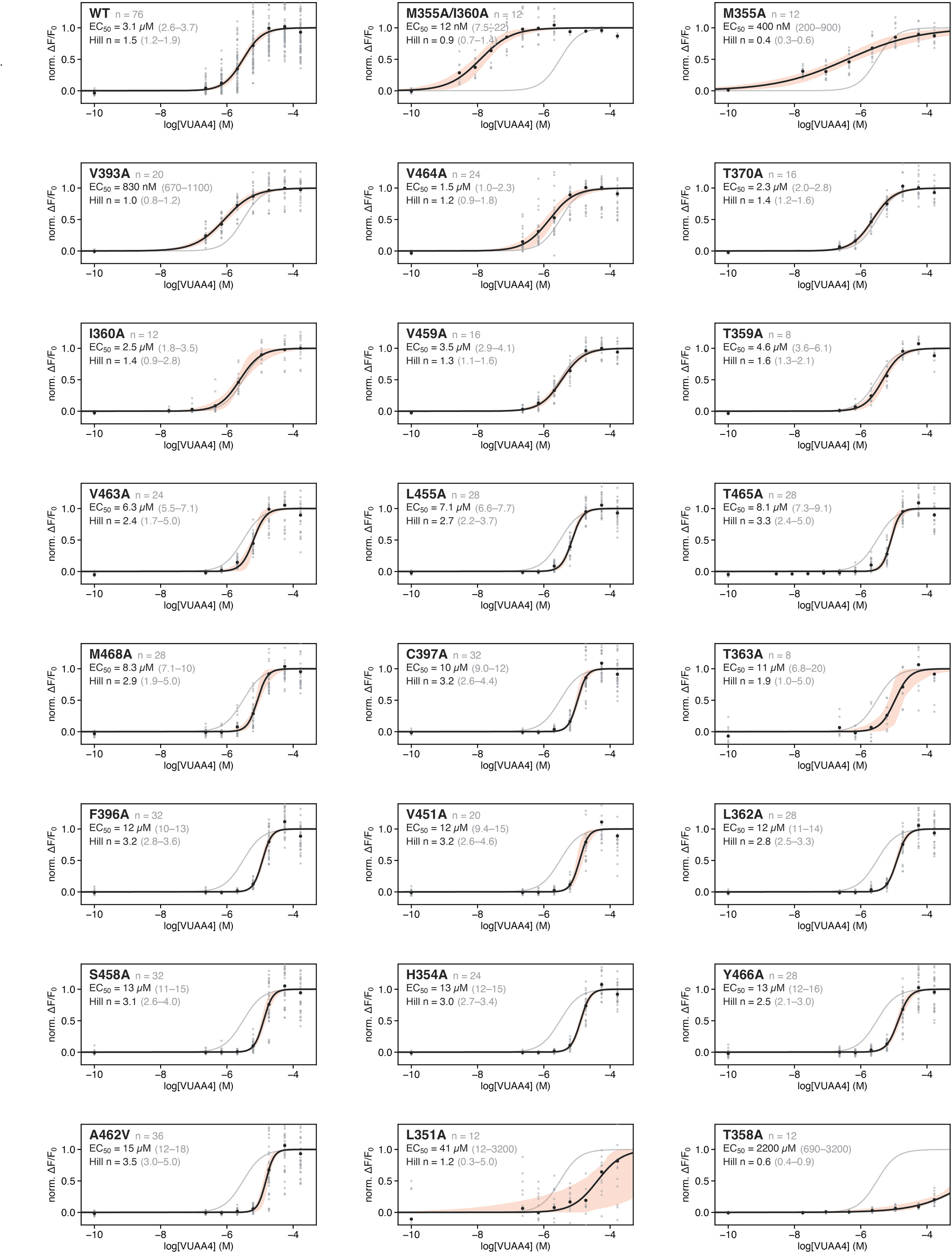
Normalized dose-response data for AbOrco WT and mutant responses to VUAA4 for calculating EC50. The WT or mutants are designated on the top left, with each raw fluorescence ΔF/F_o_ shown as points. A four-parameter Hill model (see Extended Data Fig. 9c for exact equation) was fit to the underlying data for each, and shown as a black line. 95% confidence interval of bootstrap resampling (B=500) of the underlying data was mapped as an orange envelope around the mean curve fit, highlighting points of uncertainty. The curves were scaled between 0 and 1 (baseline and maximum response), such that 0.5 represents the EC_50_. The top left plot is the reference plot, and is recapitulated as a gray line in subsequent plots for comparison. *n* represents the number of replicate dose-series, EC_50_ and Hill coefficient (Hill *n*) are provided as mean with error bounds calculated from bootstrapping (95% CI).

**Extended Data Fig. 21.**
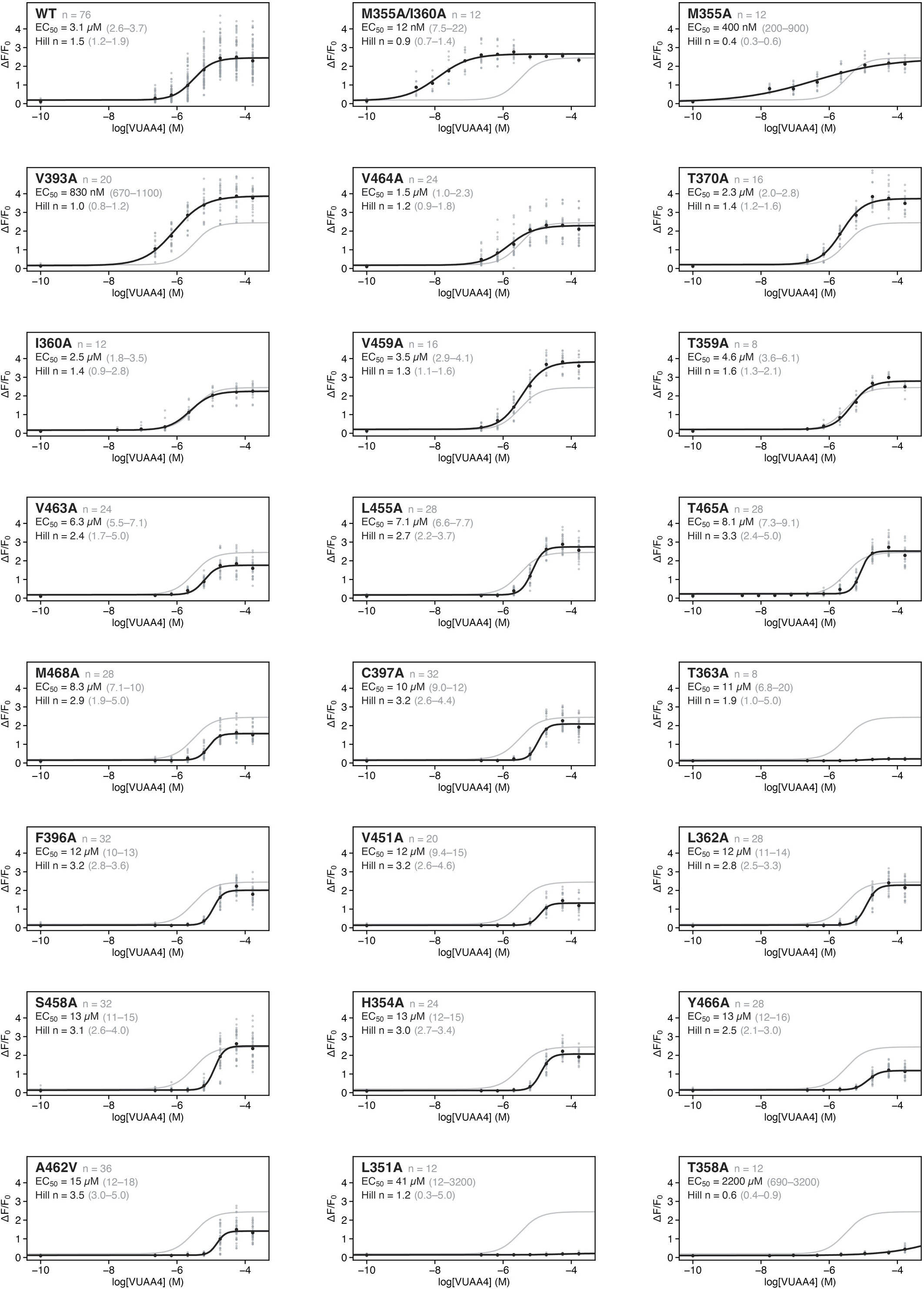
Raw dose-response data for AbOrco WT and mutant responses to VUAA4 for calculating EC50. The WT or mutants are designated on the top left, with each raw fluorescence ΔF/F_o_ shown as points. A four-parameter Hill model (see Extended Data Fig. 9c for exact equation) was fit to the underlying data for each, and shown as a black line. The curves were scaled between 0 and 1 (baseline and maximum response), such that 0.5 represents the EC_50_. The top left plot is the reference plot, and is recapitulated as a gray line in subsequent plots for comparison. *n* represents the number of replicate dose-series, EC_50_ and Hill coefficient (Hill *n*) are provided as mean with error bounds calculated from bootstrapping (95% CI).

**Extended Data Fig. 22.**
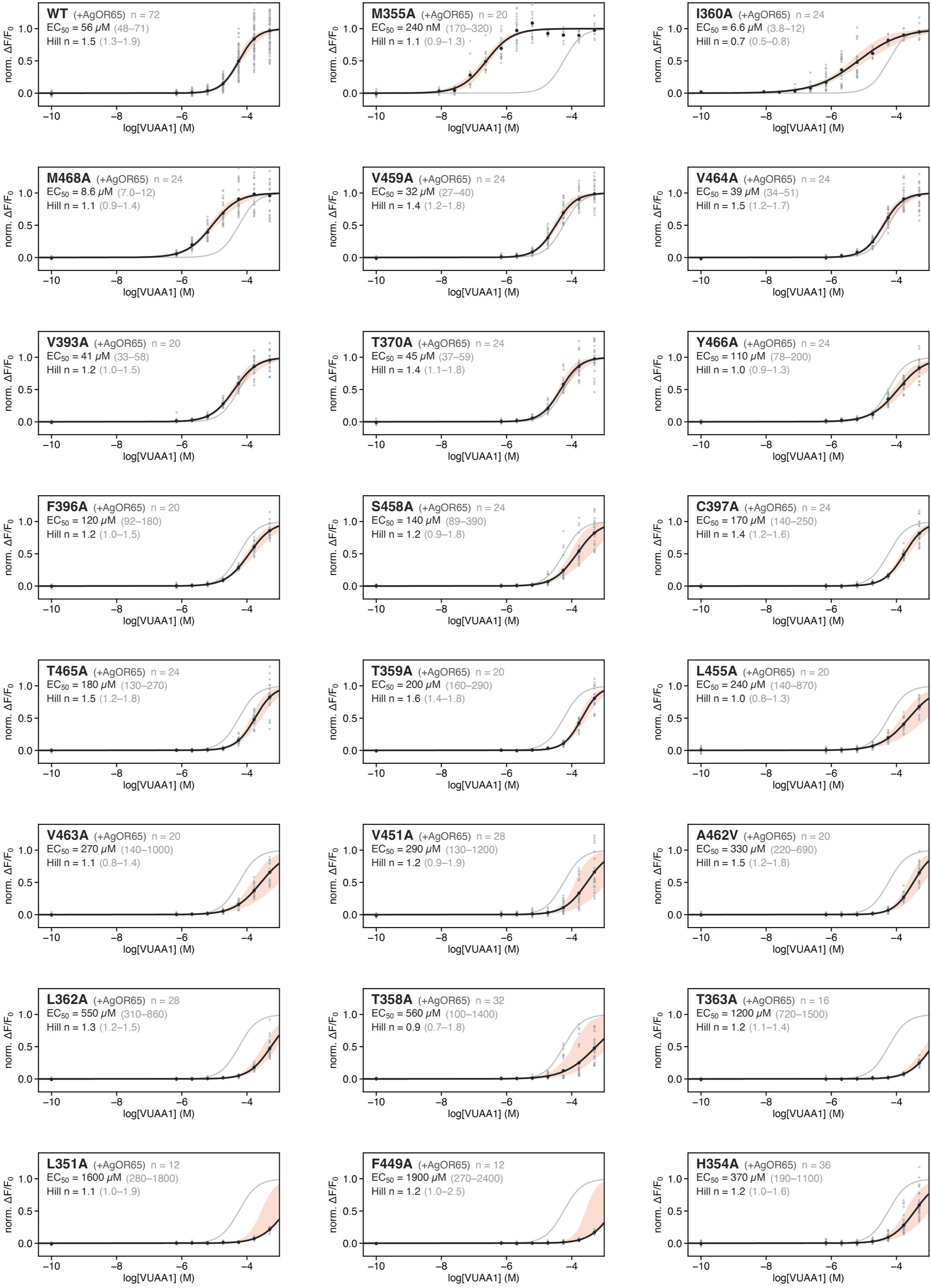
Normalized dose-response data for AbOrco (+AgOR65) WT and mutant responses to VUAA1 for calculating EC50. The AbOrco WT or mutants are designated on the top left, with each raw fluorescence ΔF/F_o_ shown as points. These measurements were performed in the presence of co-expressed *Ag*OR65 for heteromeric responses. A four-parameter Hill model (see Extended Data Fig. 9c for exact equation) was fit to the underlying data for each, and shown as a black line. 95% confidence interval of bootstrap resampling (B=500) of the underlying data was mapped as an orange envelope around the mean curve fit, highlighting points of uncertainty. The curves were scaled between 0 and 1 (baseline and maximum response), such that 0.5 represents the EC_50_. The top left plot is the reference plot, and is recapitulated as a gray line in subsequent plots for comparison. *n* represents the number of replicate dose-series, EC_50_ and Hill coefficient (Hill *n*) are provided as mean with error bounds calculated from bootstrapping (95% CI).

**Extended Data Fig. 23.**
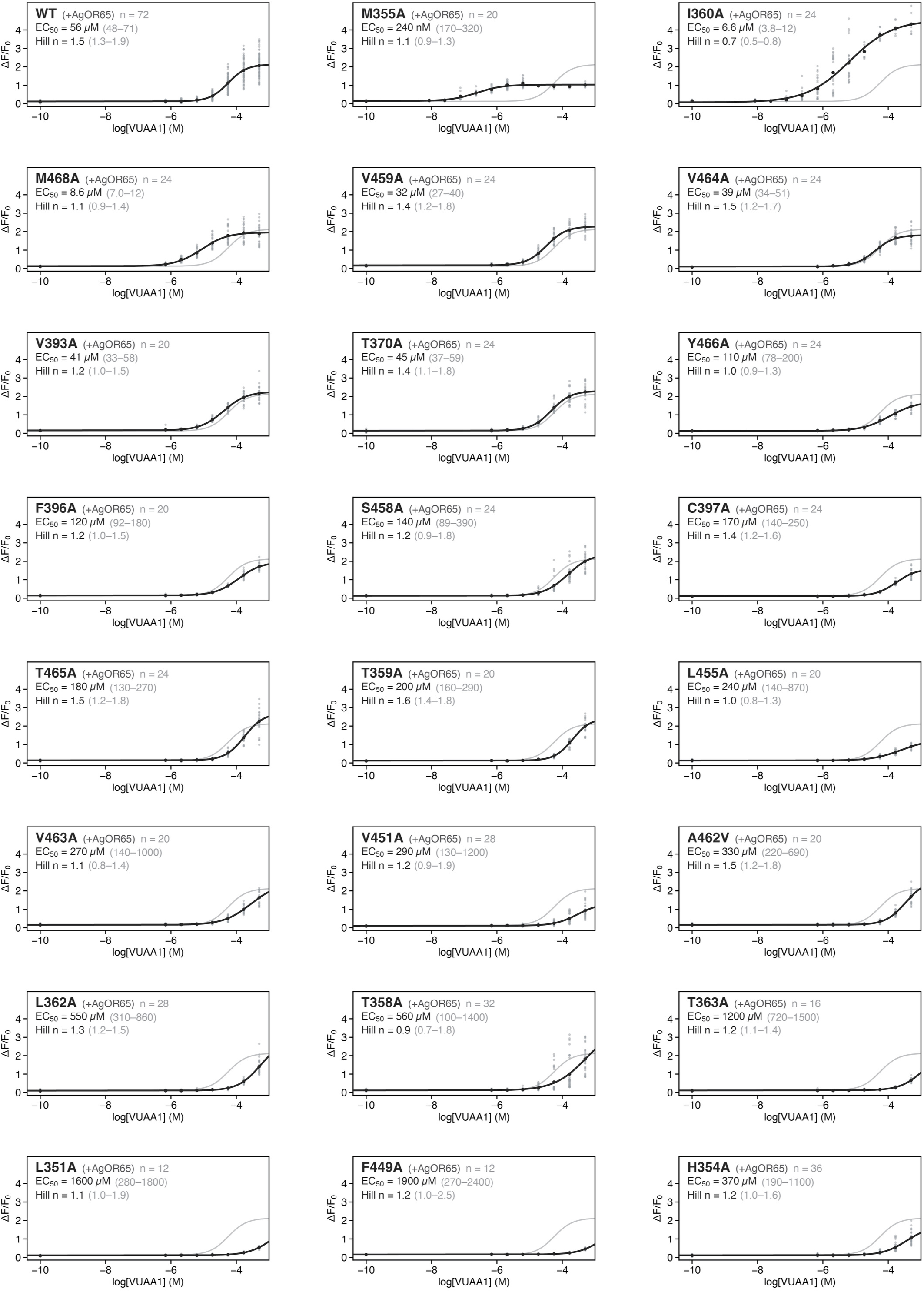
Raw dose-response data for AbOrco (+AgOR65) WT and mutant responses to VUAA1 for calculating EC50. The *Ab*Orco WT or mutants are designated on the top left, with each raw fluorescence ΔF/F_o_ shown as points. These measurements were performed in the presence of co-expressed *Ag*OR65 for heteromeric responses. A four-parameter Hill model (see Extended Data Fig. 9c for exact equation) was fit to the underlying data for each, and shown as a black line. The curves were scaled between 0 and 1 (baseline and maximum response), such that 0.5 represents the EC_50_. The top left plot is the reference plot, and is recapitulated as a gray line in subsequent plots for comparison. *n* represents the number of replicate dose-series, EC_50_ and Hill coefficient (Hill *n*) are provided as mean with error bounds calculated from bootstrapping (95% CI).

**Extended Data Fig. 24.**
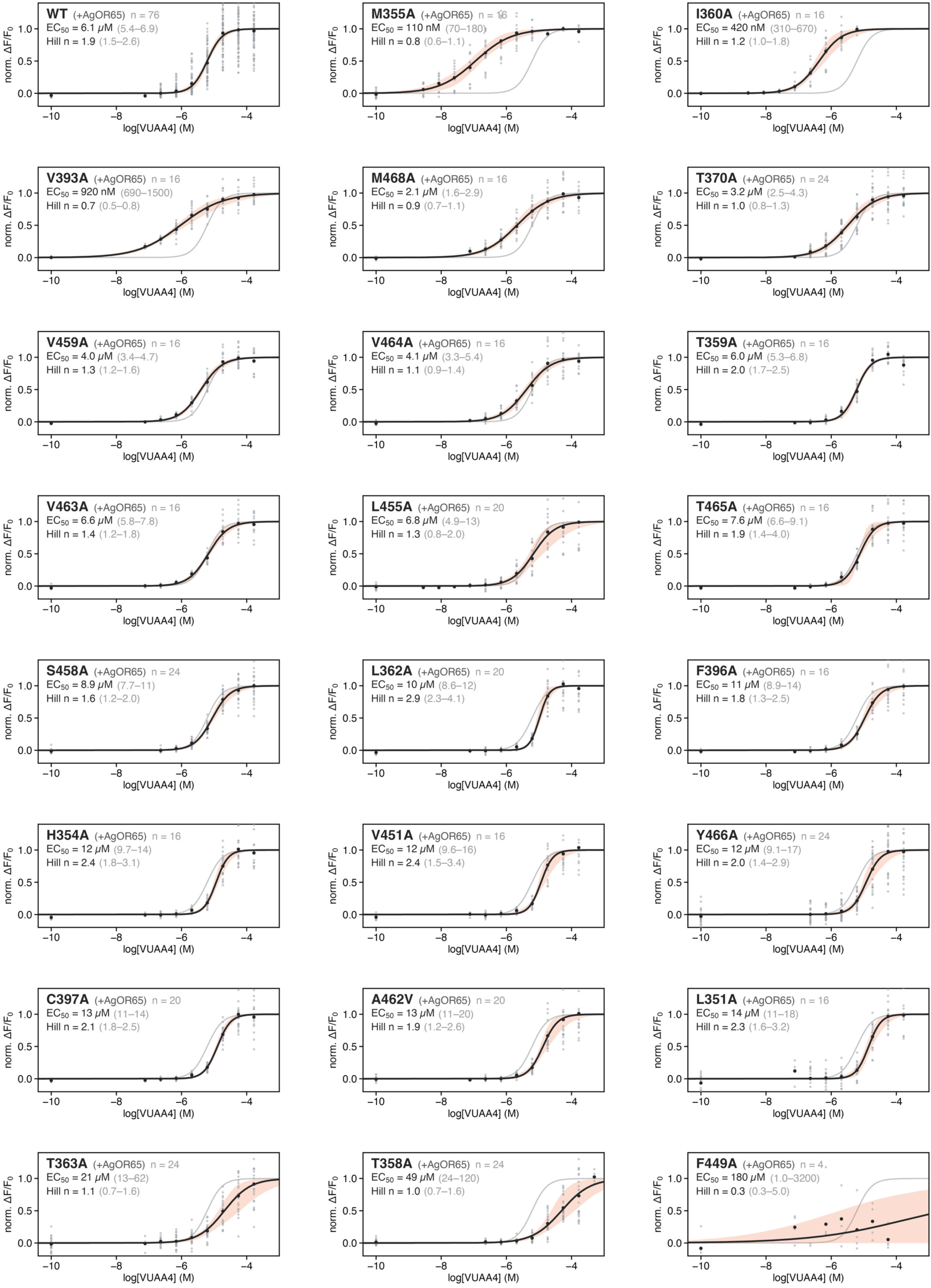
Normalized dose-response data for AbOrco (+AgOR65) WT and mutant responses to VUAA4 for calculating EC50. The *Ab*Orco WT or mutants are designated on the top left, with each raw fluorescence ΔF/F_o_ shown as points. These measurements were performed in the presence of co-expressed *Ag*OR65 for heteromeric responses. A four-parameter Hill model (see Extended Data Fig. 9c for exact equation) was fit to the underlying data for each, and shown as a black line. 95% confidence interval of bootstrap resampling (B=500) of the underlying data was mapped as an orange envelope around the mean curve fit, highlighting points of uncertainty. The curves were scaled between 0 and 1 (baseline and maximum response), such that 0.5 represents the EC_50_. The top left plot is the reference plot, and is recapitulated as a gray line in subsequent plots for comparison. *n* represents the number of replicate dose-series, EC_50_ and Hill coefficient (Hill *n*) are provided as mean with error bounds calculated from bootstrapping (95% CI).

**Extended Data Fig. 25.**
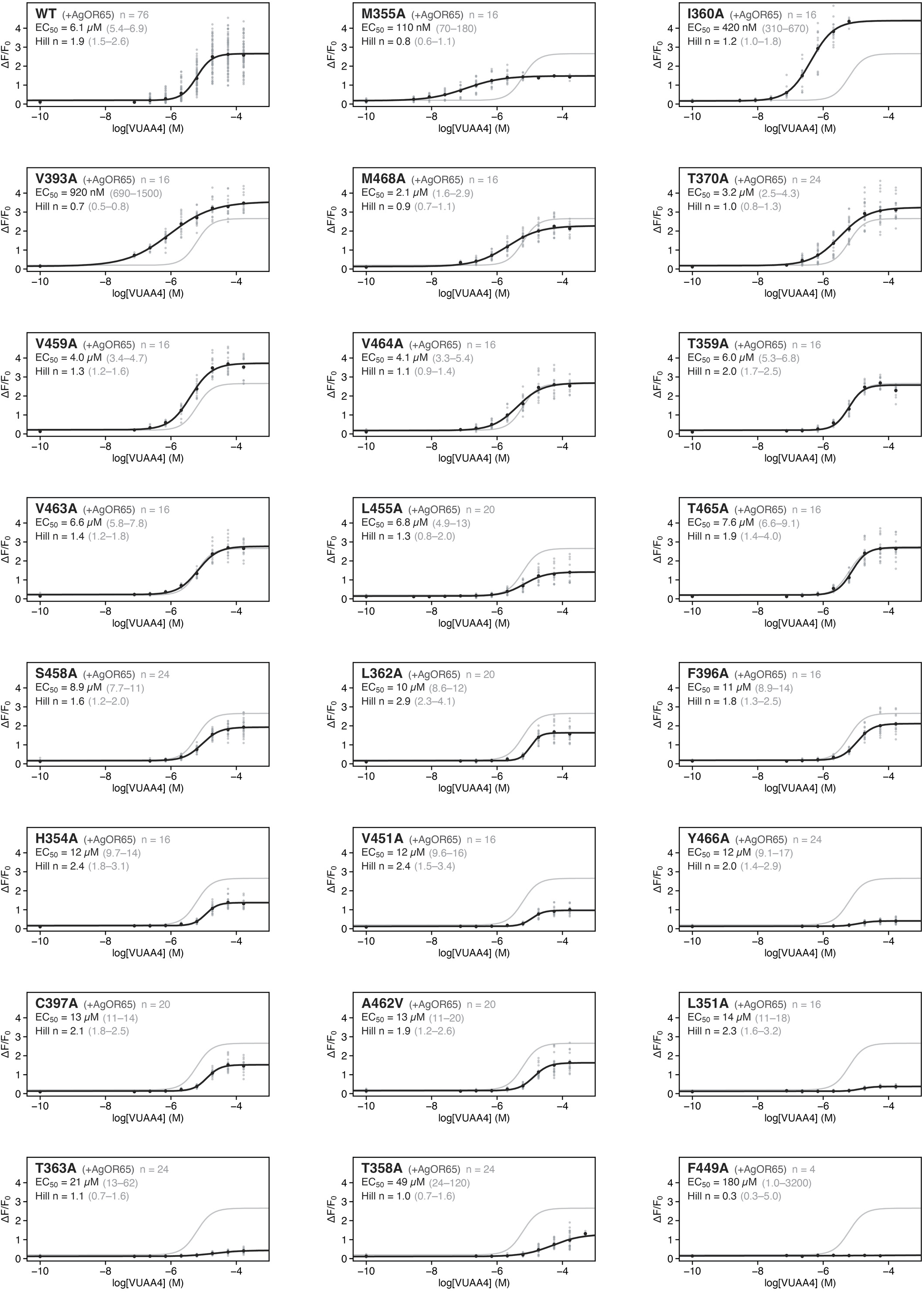
Raw dose-response data for AbOrco (+AgOR65) WT and mutant responses to VUAA4 for calculating EC50. The *Ab*Orco WT or mutants are designated on the top left, with each raw fluorescence ΔF/F_o_ shown as points. These measurements were performed in the presence of co-expressed *Ag*OR65 for heteromeric responses. A four-parameter Hill model (see Extended Data Fig. 9c for exact equation) was fit to the underlying data for each, and shown as a black line. The curves were scaled between 0 and 1 (baseline and maximum response), such that 0.5 represents the EC_50_. The top left plot is the reference plot, and is recapitulated as a gray line in subsequent plots for comparison. *n* represents the number of replicate dose-series, EC_50_ and Hill coefficient (Hill *n*) are provided as mean with error bounds calculated from bootstrapping (95% CI).

